# Anticancer, anti-inflammatory, and immune modulatory activity of ferulic acid fructo-oligosaccharide conjugated microparticle

**DOI:** 10.1101/2023.05.05.539559

**Authors:** Eldin M Johnson, Rasu Jayabalan, Samir Kumar Patra, Joo-Won Suh

## Abstract

**Background and purpose:** Ferulic acid exhibit anticancer activity but almost most of the free ferulic acid taken orally are absorbed in the stomach and extensively metabolised by the liver and hence hardly any free ferulic acid reach the large intestine to exert its beneficial activity. Fructo-oligosaccharide (dietary fibre) are resistant to gastro-intestinal enzymes and are poorly absorbed by the stomach but bioavailable in the large intestine where they are digested by gut microbiota. Ferulic acid fructo-oligosaccharide conjugate was synthesized which could self-assemble in to disc shaped amorphous microparticles, it was found to be resistant to gastro-intestinal enzymes and digestion by gut microbiota. The synthesized microparticles could be used for targeted delivery to the colon and accessed for its ability to ameliorate colo-rectal cancer and inflammation.

**Experimental approach:** The anti-cancer activity of the FA FOS microparticle (FA FOS I) was tested in human colon cancer cell lines HT29, LoVo and compared with the toxicity to normal human colon fibroblast CCD18-Co, relative to that of conventional chemotherapeutic colon cancer drug oxaliplatin. The apoptosis induction by FA FOS I was assessed by TUNNEL (Terminal deoxynucleotidyl transferase mediated dUTP Nick-end Labelling) and FACS. The ability of the FA FOS microparticle to induce cell cycle arrest was determined. The gene expression profiling of both apoptosis related genes and cell cycle arrest related genes were analysed by using RT-PCR analysis of an array of apoptosis related genes and cell cycle related genes. In-vivo pre-clinical anti-colorectal cancer studies of FA FOS I microparticle were carried out in AOM-DSS mediated colitis associated colon cancer mice model (AOM DSS CAC) to determine its anti-cancer efficacy in the physiological, immunological and innate host microbiota setting.

**Key results:** The in-vitro studies in colon cancer and normal colon cells exhibited selective cytotoxicity and apoptosis induction in colon cancer cells. The microparticle arrested the cell cycle in the G_0_-G_1_ phase. There was a reduction in 60.83% of tumour lesions in FA FOS I treated group compared to control group. The H&E histochemistry of the colon tissue revealed that there was 48.27% reduction in the malignant cell or tumour cells in the colon tissue on treatment with FA FOS I. The FA FOS conjugate treatment enhanced the gut barrier function and tight junction with the intestinal barrier guarded by the mucosal lining. The immunohistochemistry (IHC) and the immunofluorescence of the mouse colon tissue revealed the suppression of inflammation and related inflammatory cytokines in the colon. The inhibition of cell proliferation, up-regulation of tumour suppressor protein and apoptosis of the malignant or tumour cells were detected and quantified by IHC and TUNEL staining. The evaluation of immune status of the AOM DSS CAC mouse treated with FA FOS I microparticle was determined using haematological analysis of the blood lymphocytes which revealed a 9% increase in WBC count and the multiplex immunofluorescence of the colon tissue revealed an increase in the infiltration of T-helper cells and cytotoxic T-cells into the tumour microenvironment followed by the cells of the innate immune system. There was a considerable decrease in the expression of tumour suppressing PD-L1 by the tumour cells on four weeks treatment with FA FOS I microparticle.

**Conclusion and implications:** All these data implicate better efficacy of the FA FOS I microparticle delivery to colon and amelioration of colo-rectal cancer, inflammation, and positive immune modulation of tumour microenvironment against tumour proliferation.

## 1. Introduction

In-vitro screening and biosafety test is the first step in validating any therapeutic molecule (1). When screening for compounds with good anti-cancer activity and efficacy it is quiet quintessential to identify molecules with high selectivity towards cancer cells and thereby less toxic to non-cancerous or normal cells at the therapeutically relevant concentration (2). If a drug or therapeutic is capable of killing cancer cells at a lower concentration but also targets the normal or healthy cells at a similar or lower concentration then in such instances the highest tolerable dose by the patients will be insufficient to reach the target site at a therapeutically relevant drug concentration to eliminate the disease, which renders the drug useless and irrelevant (2). This present research is attempted to identify the anti-cancer activity and the mechanism of anti-proliferative function by FA FOS I microparticle synthesised by us (3) as a candidate biotherapeutic for targeted delivery to colon and amelioration of colon carcinogenesis. The FA FOS I conjugate is composed of ferulic acid (FA) esterified onto the terminal glucose unit of the fructo-oligosaccharide (FOS) followed by esterification on to every fourth fructose unit of FOS. The resultant FA FOS I conjugate self-assembled into amorphous disc shaped microparticles of 2 µm in diameter, which are insoluble in water but formed a highly homogenous suspension and resistant to gastro-intestinal enzyme and digestion by the gut microbiota (3). These characteristics makes it a suitable molecule for targeted delivery to large intestine whereby the beneficial antioxidant, anti-inflammatory and anti-cancer efficacy of FA could be exerted in the colon.

The in-vivo efficacy studies and evaluation of its safety in animal models are essential in developing a new drug candidate. Both in-vitro and in-vivo validation are carried out throughout all the stages of drug development. The selection of appropriate models and its application as well as appropriate data interpretation are important to yield basic knowledge about the drug pharmacodynamics, metabolism, and its efficacy along with the possible adverse reactions or off-targets if any, in physiological system. These in-vivo results serves as an essential step in critical decision making for the further advancement of the drug candidate for clinical trial (4). Colon cancer is the third most common cause of cancer and second most common cause of death among cancer. Even though the common causes of colorectal cancer are: (i) chromosomal instability (5), (ii) microsatellite instability and associated dysfunction of DNA mismatch repair genes and genetic hypermutability, (iii) CpG island methylation pathway, almost 30% of all colorectal cancers (CRC)(6) are found to be originating from activation of MAPK pathway by the mutually exclusive presence of BRAF mutations or KRAS mutations (7,8). The mutational landscape alone does not contribute to carcinogenesis, instead it depends on the close interaction of these mutated cells with their tumour microenvironment (TME)(9), inflammation and chronic oxidative stress (10,11). Tumour microenvironment (TME) defines the complex interaction between the immune cells and the tumour, extracellular matrix, interleukins, and other cytokines present in and around the tumour. This network of interaction determines the susceptibility of the tumour cells to the engagement of immune system and its effective function to supress or control the tumour growth. These immune cells in the tumour microenvironment also can undergo neoplastic re- programming in response to the tumour microenvironment which makes them less functional or supress their activity. Hence the success of the therapeutic intervention and the therapy itself is dependent on the tumour microenvironment its interaction with the cellular and non-cellular counterparts and the ability of the drug to positively modulate the tumour microenvironment or negative modulation suppressing the immune system and exhausting the immune system aiding in the tumour proliferation and progression to cancer (12). The non-malignant stromal cells in the periphery of tumour are found to be contributing substantially to carcinogenesis by driving an inflammatory response(8). Almost all of the CRC tumours are found to be associated with an inflammatory environment (11). The AOM DSS induced CAC mice model is an excellent model of chronic inflammation associated tumorigenesis which simulates the clinical conditions of inflammation, intact immune system and its interactions in TME, gut barrier disruption and involvement of microbiota and their metabolites contributing to tumorigenesis such as in the real-world clinical setting (13). The FA FOS I microparticle were tested for its efficacy in azoxymethane (AOM) and dextran sodium sulphate (DSS) induced colitis associated colon cancer (AOM DSS CAC) model by oral administration. In this model the immune system of the model organism is intact which enables the study of immunomodulation and recruitment of the immune cells on to the tumour microenvironment on treatment with FA FOS I and enables the study of the effective delivery of the active molecule to the target site “colon” and its efficacy in ameliorating colon cancer and its associated morbidity like inflammation. The present study is designed to evaluate the efficacy of the FA FOS I microparticle in ameliorating CRC with evaluation of the inflammatory profile, tumour cell proliferation, infiltration of immune cells in to TME and anatomical or intestinal barrier function replenishment and repair.

## 2. Materials and Methods

### 2.1 Cell culture and conditions

All cell lines were obtained from KTCC, South Korea and cultured in the prescribed cell culture media and splitting ratio. The primary cell lines were maintained in aliquots and used for the experiments which is 10 PDL (population doubling level) lesser than that prescribed by the culture collection center. CCD 18-co is human normal colon fibroblast primary cells, which were cultured in Eagle’s minimal essential medium, low glucose supplemented with 10% fetal bovine serum in T-flask. HT-29 cell line is human colorectal adenocarcinoma derived from colon epithelium of colorectal cancer patient, they were maintained in McCoy’s 5a modified medium supplemented with 10% fetal bovine serum, LoVo cell line is colorectal adenocarcinoma of human colon, which is of Duke’s type C, grade IV isolated from colorectal cancer patient, they were maintained in Ham’s F-12K medium supplemented with 10% fetal bovine serum. All the mammalian cells were maintained at 37° C at 5% CO_2_.

### 2.2 Quantitative analysis of apoptosis by Fluorescence Activated cell sorting (FACS)

To further investigate the apoptosis in the cells quantitatively, Fluorescence Activated cell sorting (FACS) analysis is performed after treating HT-29 and LoVo cells with FA FOS I at the indicated concentration for 72 hr and the cells were harvested and TUNEL reaction is performed using APO BrdU TUNEL apoptosis detection kit of Thermofisher Scientific according to the manufacturer’s instructions. FACS analysis was performed using BD LSR II instrument and the events were detected by using FACS DIVA software. The gating and analysis were performed using FlowJo V10 software.

### 2.3 Gene expression analysis for the determination of the mechanism of apoptosis induction

To investigate the mechanism of apoptosis induction by FA FOS I conjugate gene expression was profiled in LoVo cell line after the treatment of the compound at 0.198 µg/ml for 12 hr. The total m- RNA from the treated cells were extracted by using Qiagen m-RNA extraction and purification kit and after the quality of the m-RNA is tested using Qiagen RT2 PCR Array Human RNA QC, c-DNA is synthesized and the expression of 84 key apoptosis related genes were profiled by using Qiagen RT2 Profiler PCR array-Human apoptosis kit and the data was analysed by using Gene globe analysis platform of Qiagen.

### 2.4 Determination of the cell cycle arrest by FA FOS I microparticle

In order to investigate the function of FA FOS I in the cell cycle arrest, LoVo cell line was first seeded in to the culture plates and synchronized by starving them for 48 h by using culture media devoid of FBS and followed by treatment with FA FOS I and FA for 48 h for LoVo cell line and 72h for HT-29 cell line and then the population of cells which proliferated were quantitatively determined by using FACS analysis by DAPI staining followed by univariate cellular DNA content analysis. DAPI stains the DNA of the cells and depending on the DNA content the cell cycle is staged and the quantitative determination of the count of the population of the cells at each stage of the cell cycle is determined by using BD LSR II instrument and the events were analysed by using FACS DIVA software. The data was gated and plotted by using FCS express plus 6 software and the relative population of the cells at each stage were determined. To validate the FACS analysis data and to evaluate the process of cell cycle arrest, western blot analysis of the expression of cell cycle associated proteins cyclins B1, E and cyclin dependent kinase inhibitor protein p21 were determined from 72h FA FOS I treated, the total protein extract of LoVo cell line and densitometry analysis were performed by using Image J software.

### 2.5 Gene expression profiling of cell cycle related genes

Investigation of the cell cycle related genes in LoVo cell line on treatment with FA FOS I was performed as described above by using the synchronized and 24 h FA FOS I (198 µg/ml) treated cells profiled by using RT^2^ Prolifer PCR Array- Human Cell cycle from Qiagen. The data was analysed by using Gene globe platform and reactome pathway analysis.

### 2.6 In-vivo anti-colon cancer activity in AOM-DSS induced mice

The strain of mice used in induced colitis associated colon cancer mice model is C57BL/6, female mice of 5-6 weeks old were randomly assigned to six groups of seven mice each. Initially the mice were injected a single intraperitoneal injection with azoxymethane (AOM) 8 mg/Kg bodyweight of mice. Following AOM administration the mice were fed with 2% DSS for a period of one week, dissolved in drinking water (14,15). After a brief resting period of one week the mice were fed with 1.5% DSS in drinking water for another one week followed by a week rest and subsequent administration of 1% DSS for another week. This is followed by a week’s rest and the FA FOS conjugates were administered at the indicated dosage below for four weeks. The schematic representation for inducing colon cancer in mice were as depicted in figure below:

i. Naïve group (Mice without colon cancer induction)
ii. Control group (AOM-DSS induced tumour bearing mice),
iii. Oxaliplatin treatment group- 30 mg/Kg, P.O (oral administration) dissolved in vehicle- Water
iv. FA treatment group – 100 mg/Kg, P.O, vehicle- Water
v. FA FOS I treatment group- 500 mg/kg (equivalent to 100 mg/Kg free FA), vehicle- as suspension in water

After the termination of 4 weeks the mice was anaesthetized under isoflurane and the blood was extracted through cardiac puncture into heparin coated tubes, the mice were euthanized by cervical dislocation and the organs like kidney, liver, spleen, thymus, colon and femoral bone were extracted and used for further analysis. This experimental procedure is done according to the regulation and approval of Institutional Animal Care and Use Committee (IACUC) of Gachon Medical Research Institute, Le Gil Ya cancer research centre, Incheon, Korea. The colon was flushed with PBS and the number of tumour lesions in each colon were evaluated. The thymus index, spleen index, colon index was calculated from their respective organ weight and body weight. The body weight was monitored every 4 days during the treatment and the body weight gain rate per day was calculated for the treatment regime of four weeks. The total White Blood Cell (WBC) count was determined using the whole blood and the remaining blood were spun down and the plasma fraction were collected and the quantitation of interleukin 3 (IL-3) and interferon gamma (IFN-γ) were assayed by using ELISA kit from Cloud-Clone Corporation, U.S.A. The bone marrow cells were extracted by using the protocol of Liu and Ning et al., 2015(16) and the total number of cells in the femoral bone marrow of each mouse were counted. The plasma biochemistry of the mice was determined by using Fuji Dri Chem NX500 and appropriate Dri Chem slides (Fujifilm, Japan) for the respective parameter.

**Fig. 1.**
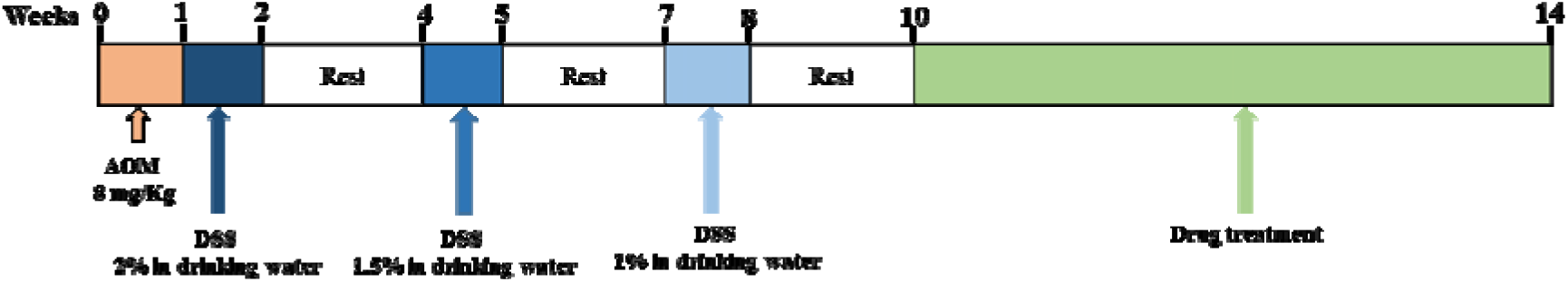
Induction of colon cancer in mice and treatment schedule The experimental groups were:

### 2.7 Histopathological analysis, Immuno histochemistry and Immunofluorescence analysis

The dissected colon from each experimental group (3 nos., from each) of AOM DSS colitis induced colon cancer mice model were rolled using swiss rolling technique of Riley and Kirsner (17) such that the full length of the colon can be visualized and examined in the histological examination. Briefly the resected colon was flushed with PBS to remove all the faecal matter and fixed in 4% formaldehyde (Sigma Aldrich) for overnight followed by paraffin embedding. The tumor tissue from the xenograft model were surgically removed and washed in PBS prior to formalin fixation and paraffin embedding. The paraffin block sections of 5 µm thickness were made on slides. The hematoxylin and eosin (H&E) staining, immuno histochemistry and immunofluorescence staining were done using BenchMark ultra, Ventana IHC/ISH auto-Stainer system of Roche diagnostics, Switzerland in a well standardised automated condition so that all the staining, counter staining and washing steps are all homogenous for all the samples. The slide was scanned using Pannoramic 250 Flash III, 3DHISTECH Ltd., Budapest, Hungary in an multifocus mode at 200x and analysed using Phenochart 1.0.12 software to determine the Dab staining, fluorescence intensity and other histopathological parameters. The tumour stromal ratio was determined from H&E staining by marking the area occupied by tumour cells and stromal cells. The analysis of epithelial lining of the intestine and connective tissues with respect to inflammation and colitis were determined by using masson’s trichrome staining.

### 2.8 Opal multiplex immunohistochemistry and cell phenotyping of immune cells in tumour microenvironment

The Swiss rolled colon from the AOM DSS mice model were subjected to Opal multiplex IHC with CD4, CD8, FOXP3, PD-L1, CD31, CTLA4 antibody counter stained with Opal 480, DIG Opal 780, Opal 620, 690, 520, 570 secondary antibody respectively along with DAPI staining of nucleus using BOND RX, automated IHC and ISH system of Leica biosystems followed by imaging using Vectra 3 system, Akoya biosciences, Delaware, USA equipped with Nunance VIS-FLEX which has liquid crystal tunable filter (LCTF), Cambridge Research and Instrumentation, Inc Massachusetts, USA. The analysis procedure is composed of the following steps: (i) Multiplex staining, (ii) Image processing involving spectrum unmixing, staining verification through pathology view, tissue segmentation and cell segmentation, (iii) Multispectral imaging scan followed by ROI- region selection, (iv) Scoring using cell phenotyping, expression level and optical density section of the specific marker, co-expression analysis and cell interaction metrics determination by inForm® 2.0 image analysis software, Akoya biosciences. Delaware.

## 3 Results and Discussion

### 3.1 Quantitative determination of apoptosis by FACS analysis

The quantitative estimation of the apoptosis induction in the CRC cell lines were performed by TUNEL apoptosis detection kit followed by FACS analysis. The genomic DNA fragmentation by cellular nucleases is one of the hallmarks of cells which undergoes apoptosis. These large number of genomic DNA fragments in the apoptotic cells results in the formation of numerous free 3’- hydroxyl terminals in them, which are labelled by bromolated deoxyuridine triphosphate nucleotides (Br-dUTP) by the enzyme deoxynucleotidyl transferase (TdT). The TdT is found to catalyse the incorporation of the deoxyribonucleoside triphosphates to the 3’ hydroxyl ends of the single stranded or double stranded DNA with either overhanging, blunt or recessed ends. It is also found that Br-dUTP is more readily incorporated into apoptotic cells’ fragmented genome compared to that of regular deoxynucleotide triphosphates which are labelled with fluorescent tags or ligands like fluorescein, digoxigenin or biotin due to stearic hindrance. Hence greater incorporation rate of Br-dUTP in the apoptotic cells and the subsequent detection of Br-dUTP by fluroscein tagged anti-BrdU monoclonal antibody serves as a suitable method for differentiating apoptotic cells from non-apoptotic cells.

It was found that upon treatment with FA FOS I greater number of cells underwent apoptosis and DNA fragmentation than free FA treatment of higher molar concentration equivalent (Fig 2). FA FOS I at 984 µg/ml showed 73.47% of cells undergoing apoptosis in HT-29 cells, while FA at 194 µg/ml (equivalent to free FA content of 984 µg/ml present in FA FOS I) showed only 7.98% cells undergoing apoptosis. This shows that FA FOS I has intrinsic anti-cancer activity independent of the FA content which may be due to its mechanosensitizing function (18) of the self-assembled microparticle, with its unique morphology and polarity. The apoptotic induction by FA FOS I increases linearly in a dose dependent manner. FA at 97 µg/ml showed 3% apoptosis while FA FOS I at 198 µg/ml showed 25.21% apoptotic cells.

**Fig. 2.**
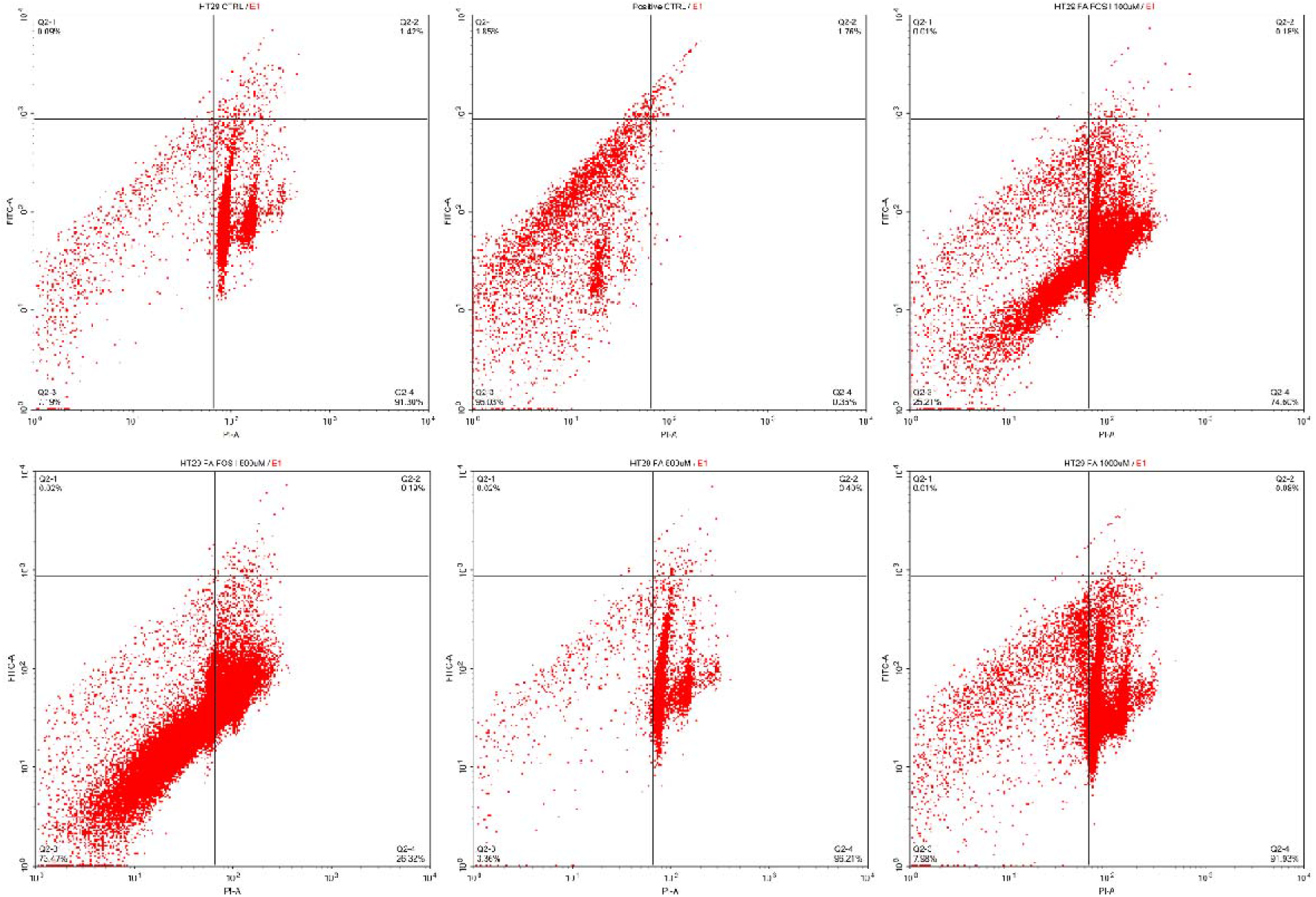
FACS analysis of TUNEL labelling of HT-29 cells (a) Negative Ctrl (un-treated cell), (b) Positive Ctrl-oxaliplatin 40µg/ml (100µM), (c) FA FOS I 198 µg/ml (100 µM) treatment, (d) FA FOS I 984 µg/ml (500 µM) treatment, (e) FA 97 µg/ml (500 µM) treatment, (f) FA 194 µg/ml (1000 µM) treatment.

**Fig. 3.**
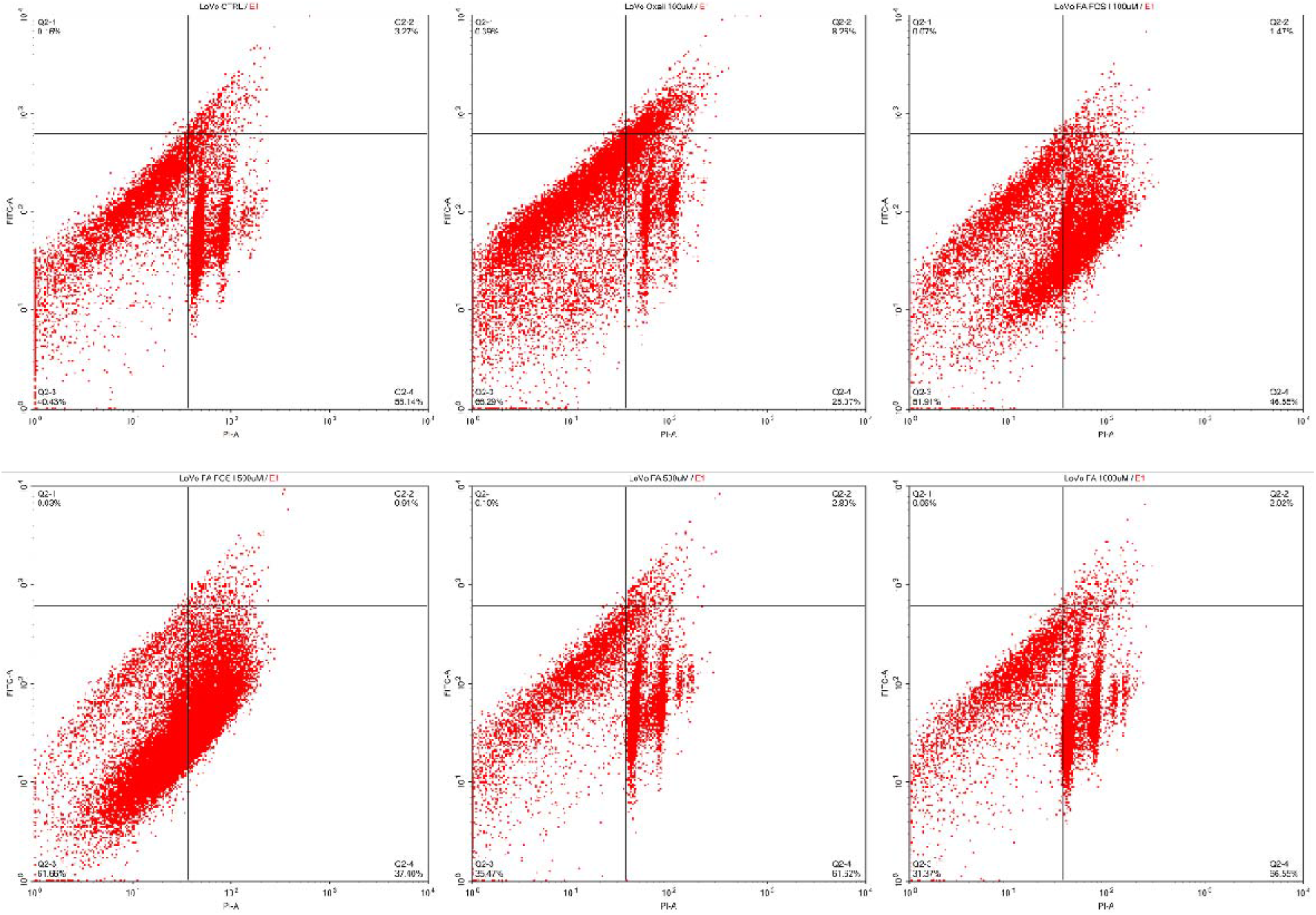
FACS analysis of TUNEL labelling in LoVo cells (a) Negative Ctrl (un-treated cell), (b) Positive Ctrl, (c) FA FOS I 198 µg/ml treatment, (d) FA FOS I 984 µg/ml treatment, (e) FA 97 µg/ml treatment, (f) FA 194 µg/ml treatment Figure 3 it depicts that in LoVo cell line also FA FOS I is exhibiting a greater effect on inducing apoptosis in a dose dependent manner than FA. FA FOS I at 198 µg/ml treatment showed 51.91 % of cell population undergoing DNA fragmentation (apoptosis), while FA FOS I at 984 µg/ml concentration showed 61.66% of cell population undergoing apoptosis. The treatment with FA at 97 µg/ml (equivalent to free FA present in 984 µg/ml of FA FOS I) showed only 35.47% apoptosis.

For testing the toxicity or apoptosis induction in non-cancerous cells, the TUNEL assay was performed in normal primary colon fibroblast cells CCD18 Co and treatment with the compounds as described above and the DNA fragmentation/apoptosis is quantitatively determined by using FACS. It was found that there was no detrimental effects noticed in the normal colon cells by treatment with FA FOS I but instead an enhanced proliferation of cells was observed with FA FOS I and FA treatment(Fig. 4). FA FOS I at 198 µg/ml showed 39.79% apoptotic dead cells and 63.07% cell proliferation compared to un-treated control which showed 42.69% dead cells and 57% proliferating cells. FA FOS I at 984 µg/ml showed an enhanced proliferating cells of 65.04% which shows a dose dependent response.

**Fig. 4.**
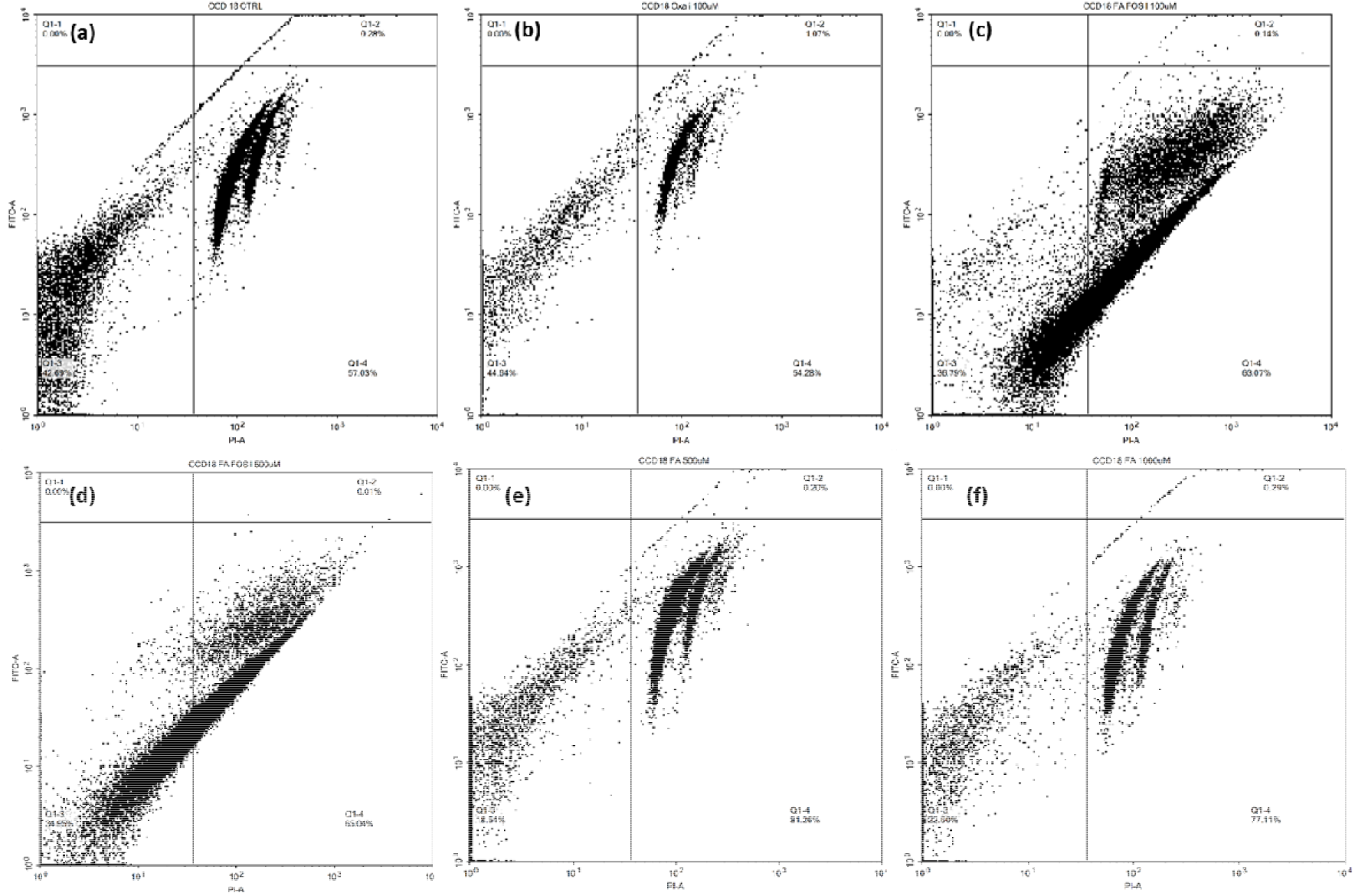
FACS analysis of TUNEL labelling in CCD18 Co cells (a) Negative Ctrl (un-treated cell), (b) Positive Ctrl oxaliplatin 40 µg/ml treatment, (c) FA FOS I 198 µg/ml treatment, (d) FA FOS I 984 µg/ml treatment, (e) FA 97 µg/ml treatment, (f) FA 194 µg/ml treatment.

The treatment with FA at the dose of 97 and 194 µg/ml showed 81.26% proliferating cells and 77.11% proliferating cells respectively showing that the proliferation rate decreases as the dosage of FA increases beyond 1000µM. There was no notable toxicity to normal cells observed from the FACS analysis.

### 3.2 Gene expression analysis to find out apoptosis in LoVo cell line

The gene expression profiling of LoVo prior to undergo apoptosis on treatment with FA FOS I at 198 µg/ml was determined after 12 h of exposure and compared with that of un-treated control. The gene expression profiles above the threshold of 1.5-fold were filtered out and summarized in table 1.

**Table: 1.** Gene expression profiling of apoptosis related genes in LoVo cell line

| Genes | Function | Fold Regulation |
| --- | --- | --- |
| TNFRSF9 | TNF-receptor superfamily- enhance T cell mediated apoptosis | 32.94 |
| CD40 | TNF-receptor superfamily receptor in antigen-presenting cells of immune system | 4.38 |
| IL10 | essential immune regulator in the intestinal tract,<br>anti-inflammatory function, inhibiting T cell<br>activation, block NF-kappa B activity | 3.79 |
| CD40LG | engages CD40, co-stimulates T-cell proliferation and<br>cytokine production, enhances production of IL4 and<br>IL10, activation of NF-kappa-B, anti-apoptotic<br>signaling | 2.97 |
| LTA | Positive role in apoptosis, cytokine produced by<br>lymphocytes are cytotoxic for a wide range of tumor<br>cells in vitro and in vivo, produced in response to<br>xenobiotic stimulus | 2.62 |
| TNFRSF11B | TNF-receptor superfamily, acts as a decoy for<br>TNFSF10/TRAIL and protect against apoptosis | 1.95 |
| FASLG | induction of apoptosis triggered by binding to FAS-also activate T-cell | 1.79 |
| CD27 | TNF-receptor superfamily- important role in the<br>apoptosis | 1.58 |
| CASP 1 | induce cell apoptosis- activate the inactive precursor<br>of interleukin-1, | 1.56 |
| CIDEA | activate apoptosis | 1.49 |
| TNFSF8 | Cytokine which binds to TNFRSF8/CD30 induce<br>proliferation of T-cells | 1.49 |
| BCL2 | Integral outer mitochondrial membrane protein-blocks the apoptotic death by BAX but not BAK | -1.50 |
| CASP5 | cysteine-aspartic acid protease (caspase) family | -1.50 |
| CRADD | a death domain (DD) motif | -1.69 |
| TNFRSF25 | regulate apoptosis | -1.81 |
| BCL2L10 | suppress apoptosis | -2.49 |
| BCL2A1 | reduce the release of pro-apoptotic cytochrome c | -33.55 |

**Table: 2.**
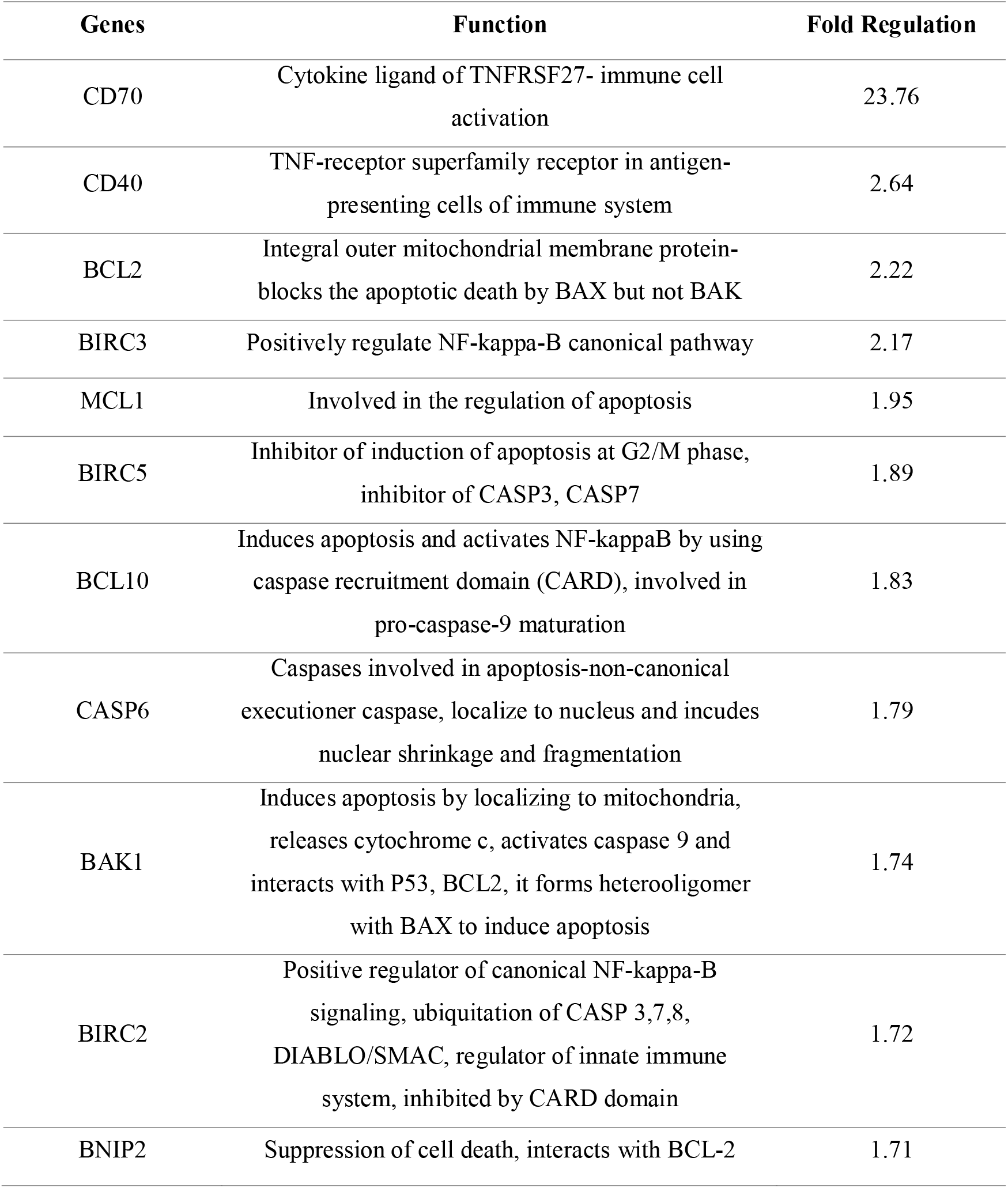
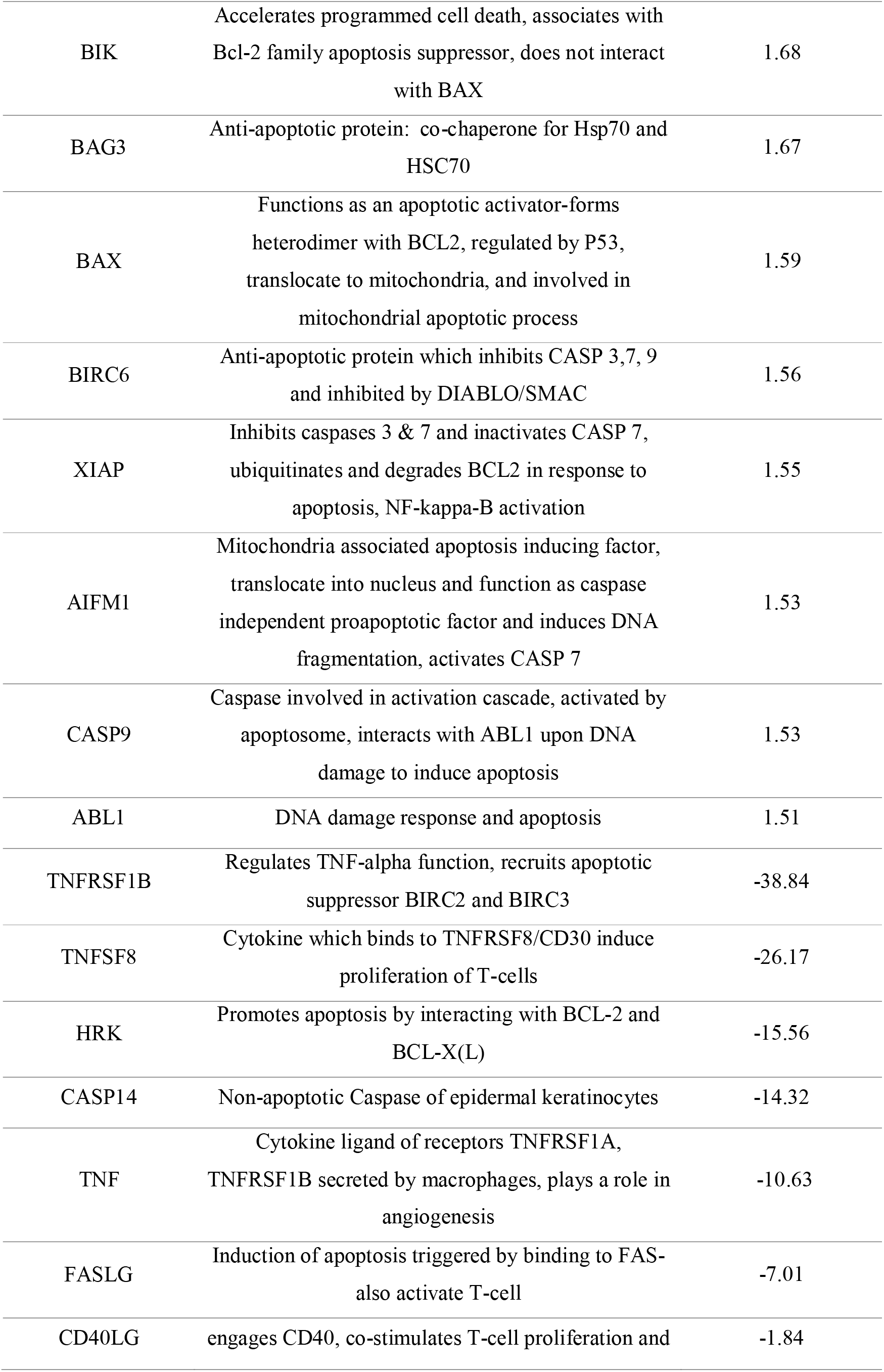
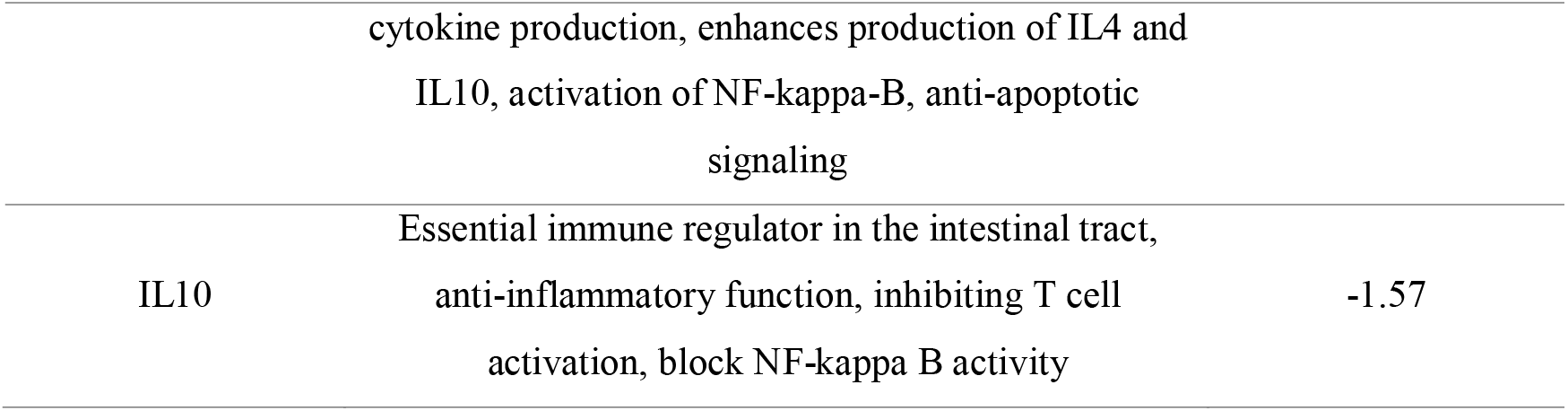
Gene expression profile of apoptosis related gene in HT-29 cell line

**Table: 3.** Gene expression profiling of cell cycle related genes in LoVo cell line

| Genes | Function | Fold Regulation |
| --- | --- | --- |
| BRCA2 | Tumor suppressor, DNA repair mechanism | 2.09 |
| BRCA1 | Plays a role in maintaining genomic stability and it also acts as a tumor suppressor | 1.67 |
| CCNT1 | Associates with cyclin dependent kinase 9 | 1.66 |
| ATR | Involved in the inhibition of DNA replication and mitosis | 1.63 |
| BCL2 | Binds to tumor suppressor p53 protein and involved in transcription-independent apoptosis of p53 | 1.61 |
| CDK 6 | Protein kinase complex that is important for cell cycle G1 phase progression | 1.54 |
| CCNG2 | Negative regulation of cell cycle progression | 1.48 |
| CDK5RAP1 | Inhibits CDK5 function which is involved in cell proliferation | 1.41 |
| BCCIP | Important cofactor of BRCA2 in tumor suppression, modulates CDK2 kinase activity through p21 | 1.40 |
| CDKN2B | Cyclin dependent kinase inhibitor complex with CDK6 and control cell cycle G1 progression | 1.40 |
| ATM | Master controller of cell cycle checkpoint signaling pathway | 1.39 |
| MCM4 | Essential for eukaryotic genome replication | -1.13 |
| SERATD1 | Apoptosis inhibitor, inhibits DNA damage response | -1.13 |
| E2F1 | Transcriptional control over cell cycle and DNA damage response | -1.08 |
| CDC34 | Ubiquitin mediated degradation of cell cycle G1 regulator | -1.04 |

From the table 1 it is apparent that 11 genes were up-regulated, and 06 genes were down-regulated on treatment with FA FOS I. In the above table it is noteworthy that TNFRSF9 (also known as CD137), which belongs to the TNF-receptor superfamily involved in the clonal expansion, survival, and development of T-cells (immune cells) was up-regulated 32.94-fold. They act as an immune stimulatory check point molecule. These molecules evoke an immune response by T-cells by activating them, induce the secretion of IL-2 (Interleukin 2) and infiltration of immune cells and eliminate tumours. This gives a clue about the possible immune cell mediated elimination of tumour/cancer cells in-vivo. There was an increase in the expression of CD40 molecule and CD40LG (ligand) which is also an immune stimulatory checkpoint molecule. IL-10 expression was also enhanced pointing towards possible anti-tumour immunity leading to control of primary tumour growth and inhibition of tumour cell metastasis in-vivo. Among the genes which are down-regulated BCL2A1 (B-cell lymphoma protein) is highly down regulated 33.55 fold, these proteins are involved in the tumour progression and are over expressed in a variety of cancer cells, down regulation of this gene sensitizes the tumour cells for apoptosis and improve the efficacy of anticancer therapy (19). Another noteworthy molecule BCL2L10 which is an anti-apoptotic protein was down-regulated 2.49- fold on treatment with FA FOS I.

The gene expression data were analysed using reactome pathway analysis (20) and top five most significant pathways were elucidated from the gene expression profile : (i) TNF binds to their physiological receptor (Fig.5), (ii) Signalling by interleukins, (iii) BH3 only proteins associate with and in-activate anti-apoptotic BCL2 members (Fig.6), (iv) activation of BAD and translocation to mitochondria, (v) Foxo mediated transcription of cell death genes. The members of the TNFSF (tumour necrosis factor superfamily) and TNFRSF (TNF receptor superfamily) play a vital role in both innate and adaptive immunity by positively regulating immune system (21). Interleukins are low molecular weight proteins that binds on to the cell surface receptors and act in a paracrine and/or autocrine mode. Initially they were known to be expressed by leukocytes but later found to be produced by many other cell types throughout the body which regulates tissue growth and repair, hematopoietic homeostasis and multiple levels of defence against pathogens where they are an integral part of the immune system (22). The Bcl-2 is found to interact with tBid, BIM, PUMA, NOXA, BAD and BMF resulting in inactivation of BCL2 which results in the prevention of the binding of BCL2 to tBid which in-turn causes the release of cytochrome C from mitochondria and causing apoptosis. BH3 only proteins associate with and inactivate the anti-apoptotic BCL-XL gene promoting apoptosis (23,24). The intrinsic (Bcl-2 inhibitable or mitochondrial) pathway of apoptosis functions in response to intracellular stress including withdrawal of growth factor, DNA damage, death receptor stimulation, protein unfolding stresses in the endoplasmic reticulum etc., Following the reception of stress signals, proapoptotic BCL-2 family proteins are activated and anti-apoptotic BCL-2 proteins are inactivated by interaction with them. This leads to the destabilization of the mitochondrial membrane and release of apoptotic factors which induces proteolytic caspase cascade, chromatin condensation and DNA fragmentation ultimately leading to cell death. The key players in the intrinsic pathway are the Bcl-2 family of proteins that are the key death regulators residing immediately upstream of mitochondria (25). The switching on/off the BAD activity happens through its de-phosphorylation by calcineurin and phosphorylation by Akt1 respectively. The activated BAD translocate in to mitochondria and induces apoptosis (26). FOXO transcription factors promote the expression of several pro-apoptotic genes such as FASLG (27), PINK1, BIM, BCL6, PUMA and contributes towards tumour suppressive role of FOXO factors (28).

**Fig. 5.**
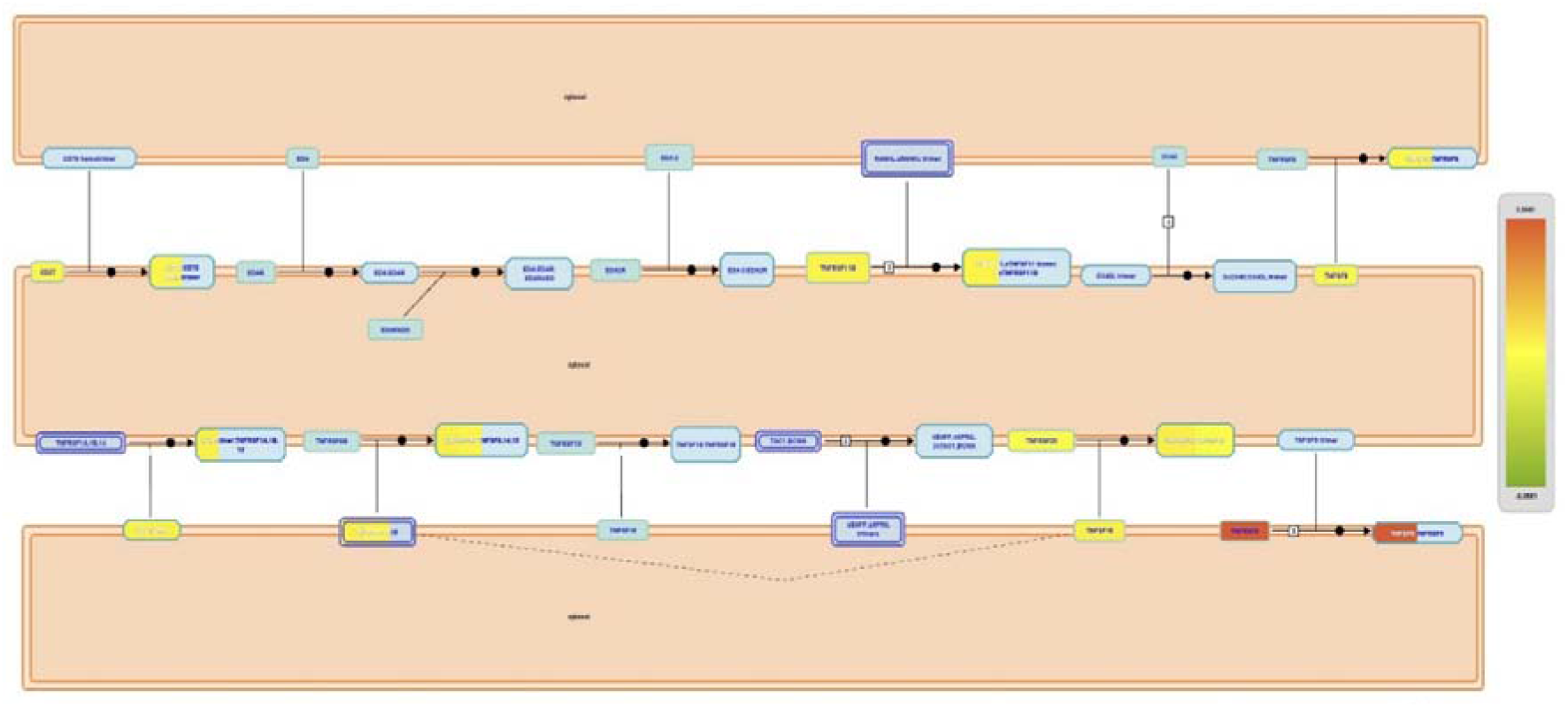
TNF binding to their physiological receptor. Pathway derived from reactome analysis showing relevant gene expression in the pathway; coloured according to the expression level

**Fig. 6.**
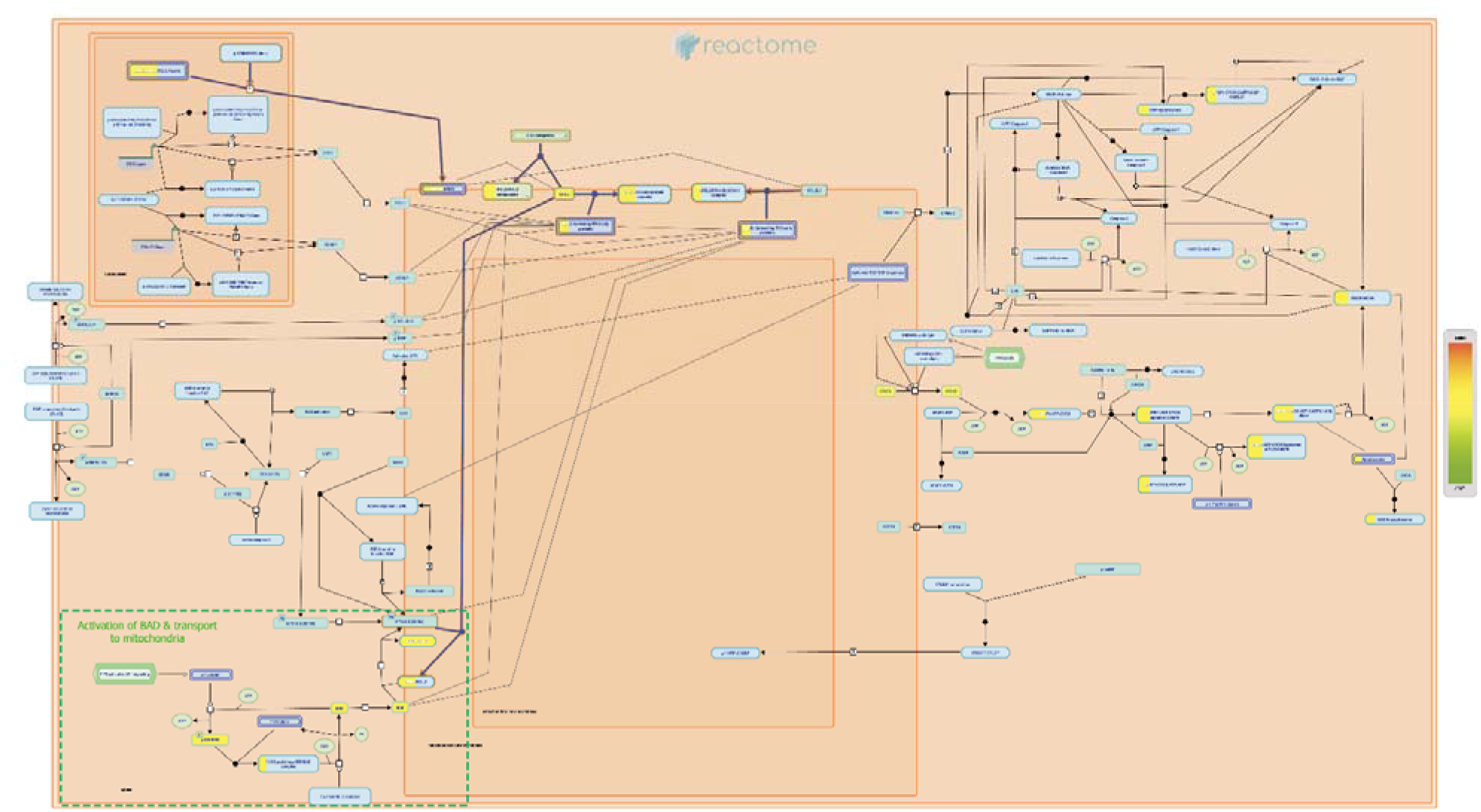
BH3 only proteins associate with and in-activate anti-apoptotic BCL-2 members, Marked region- Activation of BAD and translocation to mitochondria. Pathway derived from reactome analysis showing relevant gene expression in the pathway coloured according to the expression level

### 3.3 Gene expression analysis for the determination of molecules associated with apoptosis induction in HT-29 cell line

The gene expression profile of HT-29 cell line revealed that 19 genes were up-regulated and about eight genes were significantly down-regulated on treatment with FA FOS I. Among the up-regulated genes most of them are related to immune stimulation, NF κ-B activation and mitochondrial pathway of apoptosis. When the differentially expressed genes were fed to reactome pathway analysis, it revealed four significant pathways according to the context of the gene expression profile: (i) TP53 regulated transcription of cell death genes, (ii) TNF receptor superfamily mediated NF-κB pathway, (iii) Mitochondrial apoptotic factor mediated response, (iv) SMAC (DIABLO) binds to IAPs (Inhibitor of Apoptotic Proteins).

The tumour suppressor TP53 imparts its tumour suppressive role partly by regulating the transcription of several genes involved in apoptotic cell death. Most of the apoptotic genes are the transcriptional targets of TP53 but there are also several TP53 target genes which inhibit apoptosis providing opportunity for the cells to attempt to repair the damage and recover from stress. Even though the expression of TP53 is highly dis-regulated in HT-29 cell line because of the inherent mutation in them, the treatment with FA FOS I has tilted the TP53 pathway towards apoptosis. The transcriptional targets of TP53 which are involved in the intrinsic pathway of apoptosis triggered by cellular stress are BAX, BID, AIFM1, TP53AIP1 which regulate the permeability of mitochondrial membrane and/or release cytochrome C release. Other pro-apoptotic genes which are either involved in the intrinsic or extrinsic apoptosis pathway and pyroptosis (inflammation-related cell death) are transcriptionally regulated by TP53 are cytosolic caspase activators, APAF1 and caspases such as CASP1, CASP6 and CASP10. TP53 is stabilized in response to cellular stress by phosphorylation at the serine residues S15 and S20 followed by the activation of cell death genes, the TP53 tetramer phosphorylated at S15 and S20 is known to act as a regulator of pro-apoptotic/pro-cell death genes. Activation of the pro-apoptotic TP53 target such as BAX, FAS, BBC3 require the presence of the complex between TP53 and ASPP protein either ASPP1 or ASPP2, depending on which the interaction with the specific co-factors modulates the cellular response/outcome (29).

**Fig. 7.**
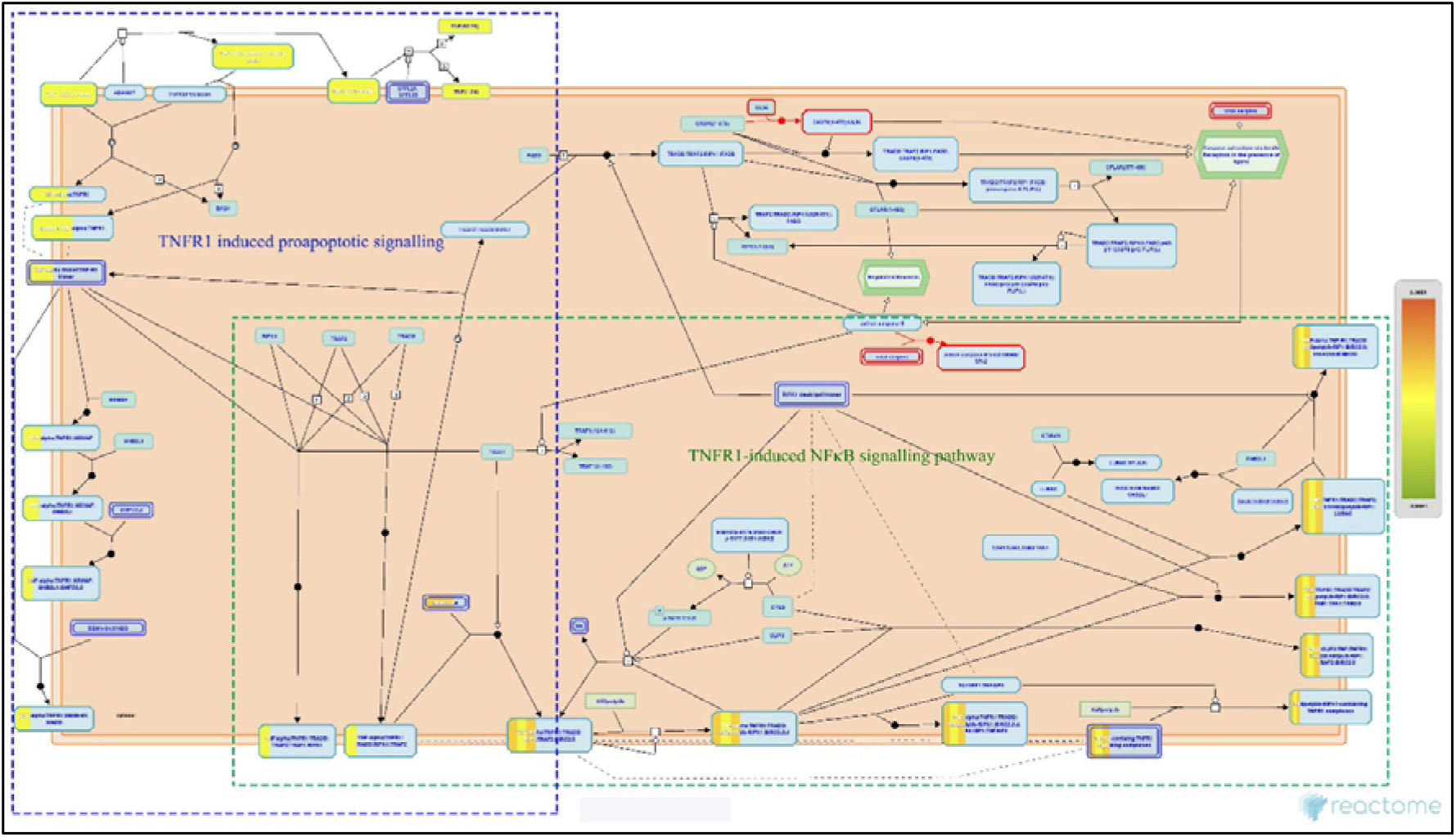
TNF receptor superfamily (TNFSF) members mediating NF-κB (nuclear factor kappa B) signalling pathway

Activation of NF-κB is fundamental for signal transduction by members of TNFRSF, the expression of NF-κB target genes is essential for mounting innate immune responses against infectious agents. NF- κB transcription factor family are activated by two distinct pathways: canonical pathway involving NF- κB1 and the non-canonical pathway involving NF- κB2. Unlike NF- κB1 signalling which can be activated by a wide variety of receptors the NF- κB2 is typically activated by a subset of receptor and ligand pairs belonging to TNFRSF superfamily members. Activation of tumour necrosis factor receptor 1 (TNFR1) can trigger multiple signal transduction pathway to induce inflammation, cell proliferation, cell survival or cell death. Whether TNF-alpha stimulated cell would survive, or die is dependent on the cellular context. TNF-alpha induced signals lead to the activation of transcription factors such as NF-κB and activator protein-1 (AP1). The binding of TNF-alpha to TNFR1 leads to recruitment of the adapter protein TNFR1-associated death domain (TRADD) and of receptor interacting protein 1 (RIPK1). TRADD subsequently also recruits TNF receptor-associated factor 2 (TRAF2). RIPK1 is promptly K63-polyubiquitinated which results in the recruitment of the TAB2:TAK1 complex and the IkB kinase (IKK) complex to TNFR1. The activated IKK complex mediates phosphorylation of the inhibitor of NF-κB (IkB), which targets IkB for ubiquitination and subsequent degradation. Released NF-κB induces the expression of a variety of genes including inflammation-related genes and anti-apoptotic genes encoding proteins such as inhibitor of apoptosis proteins cIAP1/2, Bcl-2, Bcl-xL or cellular FLICE-like inhibitory protein (FLIP). While pro-survival signalling is initiated and regulated via the activated TNFR1 receptor complex at the cell membrane, cell death signals are induced by internalization-associated fashion upon the release of RIPK1 from the membrane complex (30). This non-ubiquitylated RIPK1 induces either apoptosis or necroptosis. The action of caspase inactivates RIPK1 and RIPK3 by cleavage (30).

**Fig. 8.**
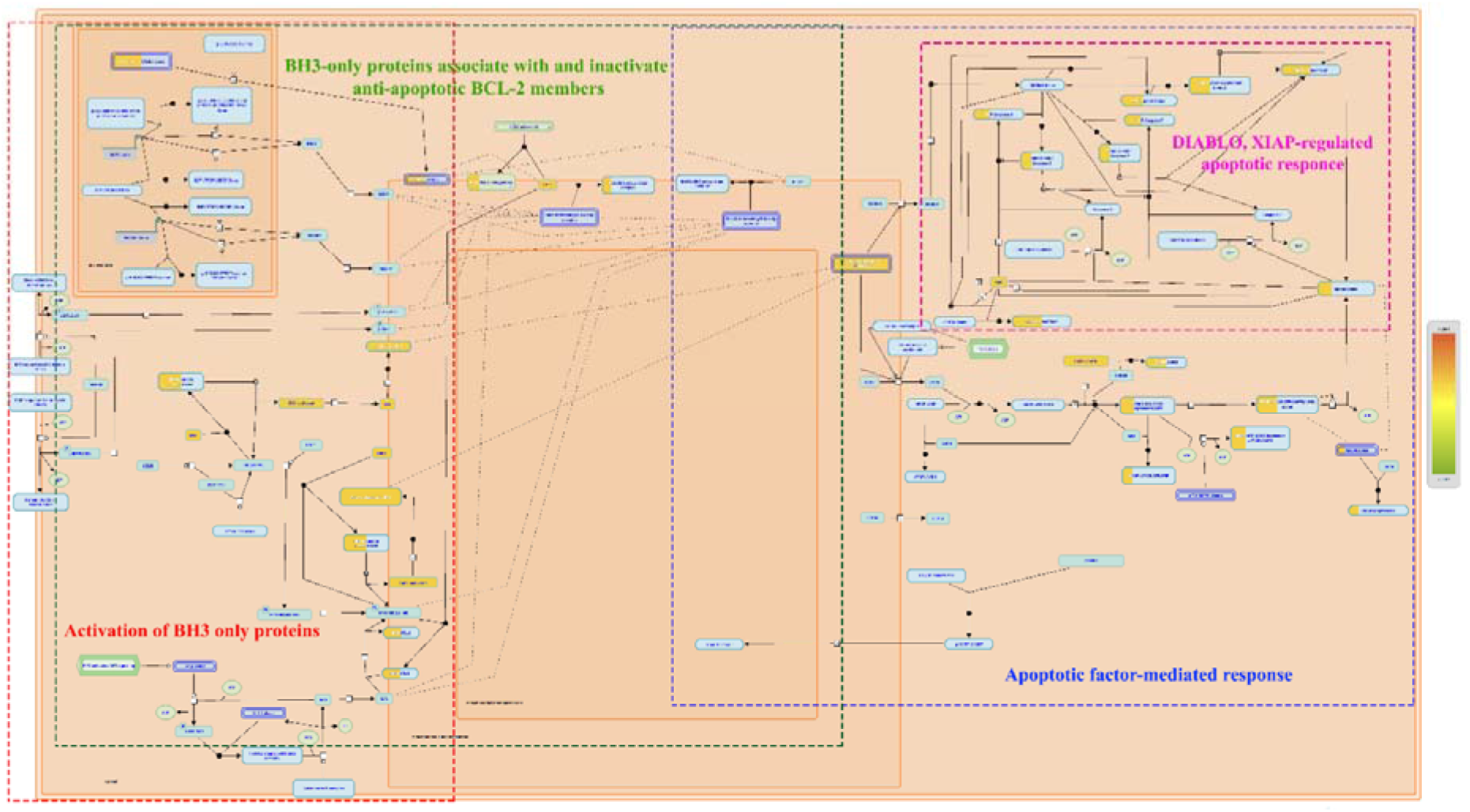
Mitochondrial pathway of apoptosis in HT-29 cell line on treatment with FA FOS I

In response to apoptotic signals, mitochondrial proteins are released into the cytosol and activate both caspase-dependent and -independent cell death pathways. Cytochrome c induces apoptosome formation, AIF and endonuclease G function in caspase independent apoptotic nuclear DNA damage. Smac/DIABLO and HtrA2/OMI promote both caspase activation and caspase-independent cytotoxicity (31). The intrinsic (Bcl-2 inhibitable or mitochondrial) pathway of apoptosis functions in response to various types of intracellular stress including growth factor withdrawal, DNA damage, unfolding stresses in the endoplasmic reticulum and death receptor stimulation. Following the reception of stress signals, proapoptotic BCL-2 family proteins are activated and subsequently interact with and inactivate antiapoptotic BCL-2 proteins. This interaction leads to the destabilization of the mitochondrial membrane and release of apoptotic factors. These factors induce the caspase proteolytic cascade, chromatin condensation, and DNA fragmentation, ultimately leading to cell death. The key players in the Intrinsic pathway are the Bcl-2 family of proteins that are critical death regulators residing immediately upstream of mitochondria. The Bcl-2 family consists of both anti- and proapoptotic members that possess conserved alpha-helices with sequence conservation clustered in BCL-2 Homology (BH) domains. Proapoptotic members are organized as follows:

1. "Multidomain" BAX family proteins such as BAX, BAK etc. that display sequence conservation in their BH1-3 regions. These proteins act downstream in mitochondrial disruption.
2. "BH3-only" proteins such as BID, BAD, NOXA, PUMA, BIM, and BMF have only the short BH3 motif.

These act upstream in the pathway, detecting developmental death cues or intracellular damage. Anti-apoptotic members like Bcl-2, Bcl-XL and their relatives exhibit homology in all segments BH1-4. One of the critical functions of BCL-2/BCL-XL proteins is to maintain the integrity of the mitochondrial outer membrane.

Second mitochondria derived activator of caspases protein (SMAC, also known as direct IAP binding protein with low pI or DIABLO) in its dimeric form interacts and antagonizes X linked inhibitor of apoptosis protein (XIAP) by concurrently targeting both BIR2 and BIR3 domains of XIAP (32). XIAP inhibits apoptosis by binding to and inhibiting the effectors caspase 3 and 7 and an initiator caspase 9 (33). During apoptosis, SMAC (DIABLO) is released from the mitochondria. In the cytosol, SMAC binds to XIAP displacing it from caspase: XIAP complexes liberating the active caspases. SMAC regulates XIAP function and potentiates caspase 3, -7 and -9 activity by disrupting the interaction of caspases with XIAP. Residues 56-59 of SMAC (DIABLO) are homologous to the amino-terminal motif that is used by caspase-9 (CASP 9) to bind to the BIR3 domain of XIAP. SMAC (DIABLO) competes with CASP 9 for binding to BIR3 domain of XIAP promoting the release of XIAP from the CASP9: apoptosome complex (33). The binding of SMAC to the BIR2 and BIR3 regions of XIAP creates a steric hindrance that is essential for preventing binding of XIAP linker region with effector caspases CASP3 and CASP7 thus achieving neutralization of XIAP inhibition. The strong affinity for XIAP allows SMAC (DIABLO) to displace caspase 3, -7 from the XIAP: caspase complexes.

**Fig. 9.**
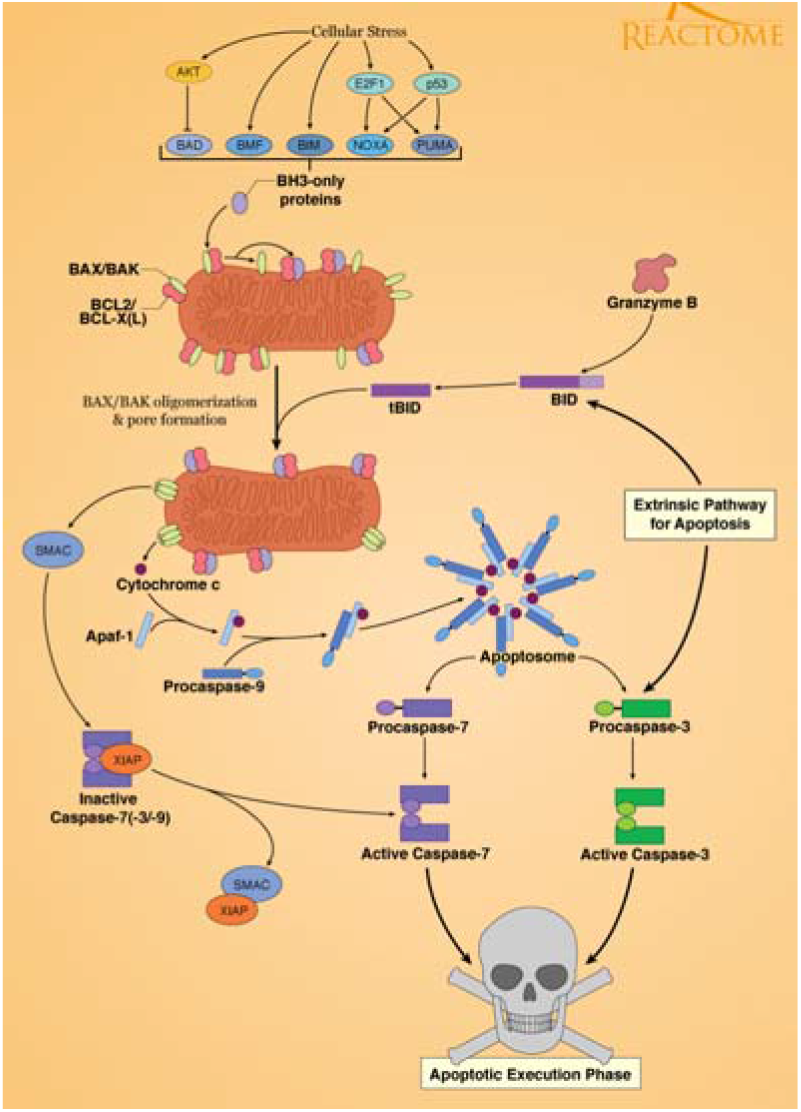
Intrinsic pathway of apoptosis, *Source-Reactome database

In both the cases of LoVo and HT-29 cell line the induction of apoptosis is mediated by mitochondria, which is a phylogenetically and evolutionarily ancient mode of cell death or apoptosis. Aberrant cell death is one of the traits of cancer cells and tumor progression which develop dysregulation of mitochondrial pathway leading to apoptotic resistance and survival. Various therapeutics have been developed to target Bcl-2 family which are pro-survival proteins resisting apoptosis (34) either by blocking at the translational level using oligonucleotides such as in the case of oblimersen sodium (35), another strategy is by developing mimetics of BH3 which could sequester or block the anti-apoptotic Bcl2 allowing pro-apoptotic proteins to induce apoptotic cell death such as in ABT-263 developed by Abott (36)and JY-1-106 which is also a BH3 mimetic to induce apoptosis in colon cancer, mesothelioma and lung cancer (37). Therapeutics with SMAC mimetics which could inhibit IAPs are also under clinical trial to induce apoptosis (38), anti-sense oligonucleotide against XIAP has also been developed for soft tissue cancers (39). Various senescent cells are found to be resistant to apoptosis by over expressing BCL-xL and other BCL-2 family members and not heeding to apoptotic signals. They are found to cause inflammation and damage to nearby cells by their secretions. Among the 46 drugs and natural compounds Quercetin (flavonoid), curcumin, luteolin, piper longumine were found to be having senolytic properties and inducing apoptosis (40) while not destroying neighbouring normal cells. Here we report the apoptosis induction in cancer cell line by ferulic acid conjugated fructo oligosaccharide microparticles which could tip the balance towards apoptosis induction.

### 3.4 Cell cycle arrest by FA FOS I microparticle

The function of FA-FOS I in the cell cycle arrest was performed in LoVo and HT-29 cell line. FA- FOS I arrested the cell cycle at G_1_ phase of the cell cycle in a dose dependent manner. In HT-29, FA- FOS I at 198 µg/ml and 988 µg/ml, the cell population which was stalled at G_1_ phase was 91.22% and 94.16% respectively (Fig 10a). While in the case of LoVo cell line FA FOS I at 198 µg/ml and 988 µg/ml 62.74% and 79.52 % of cells were found at G_1_ phase compared to 48.64% of cells in the population of cells at control group (Fig 10b).

**Fig. 10.**
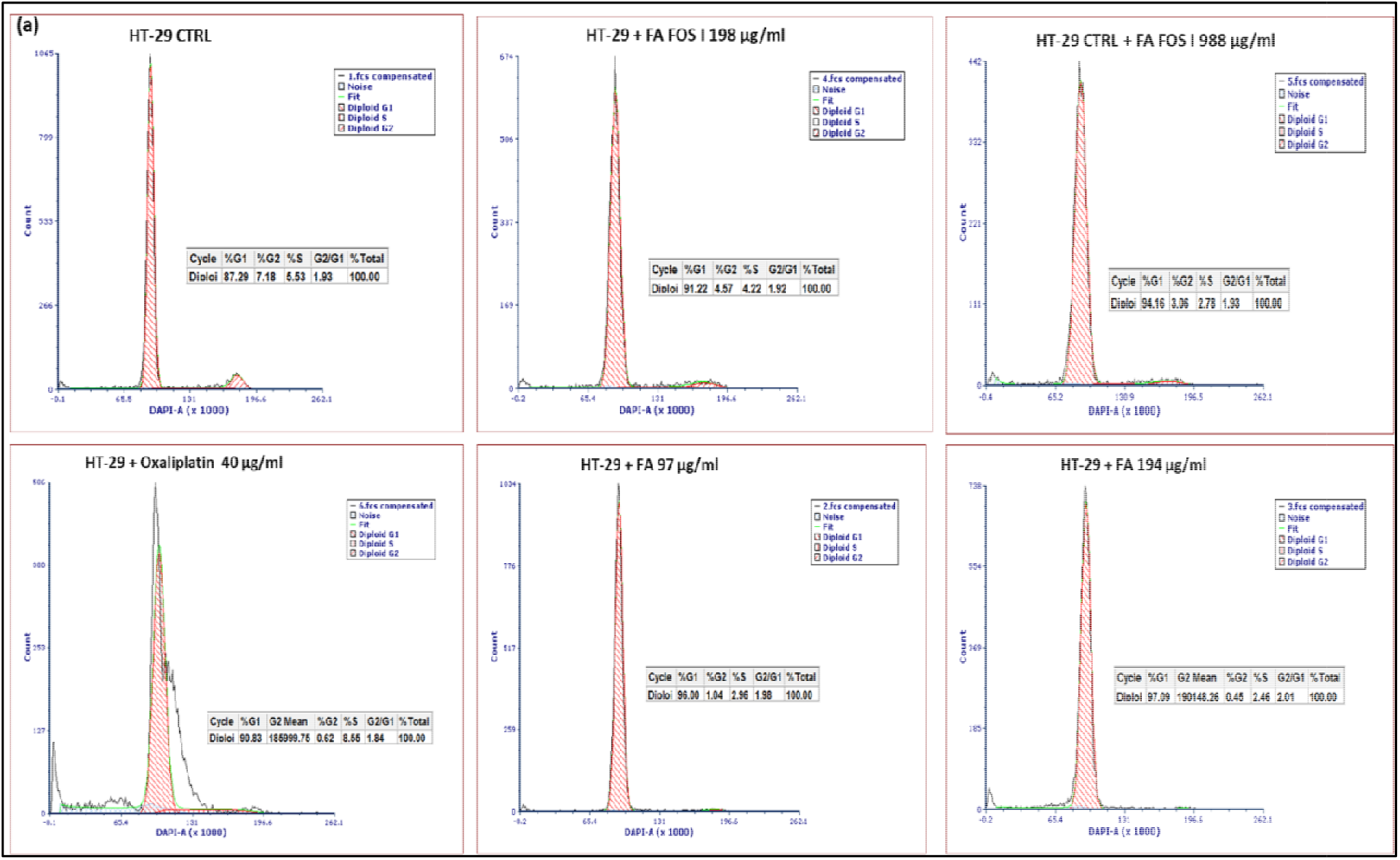
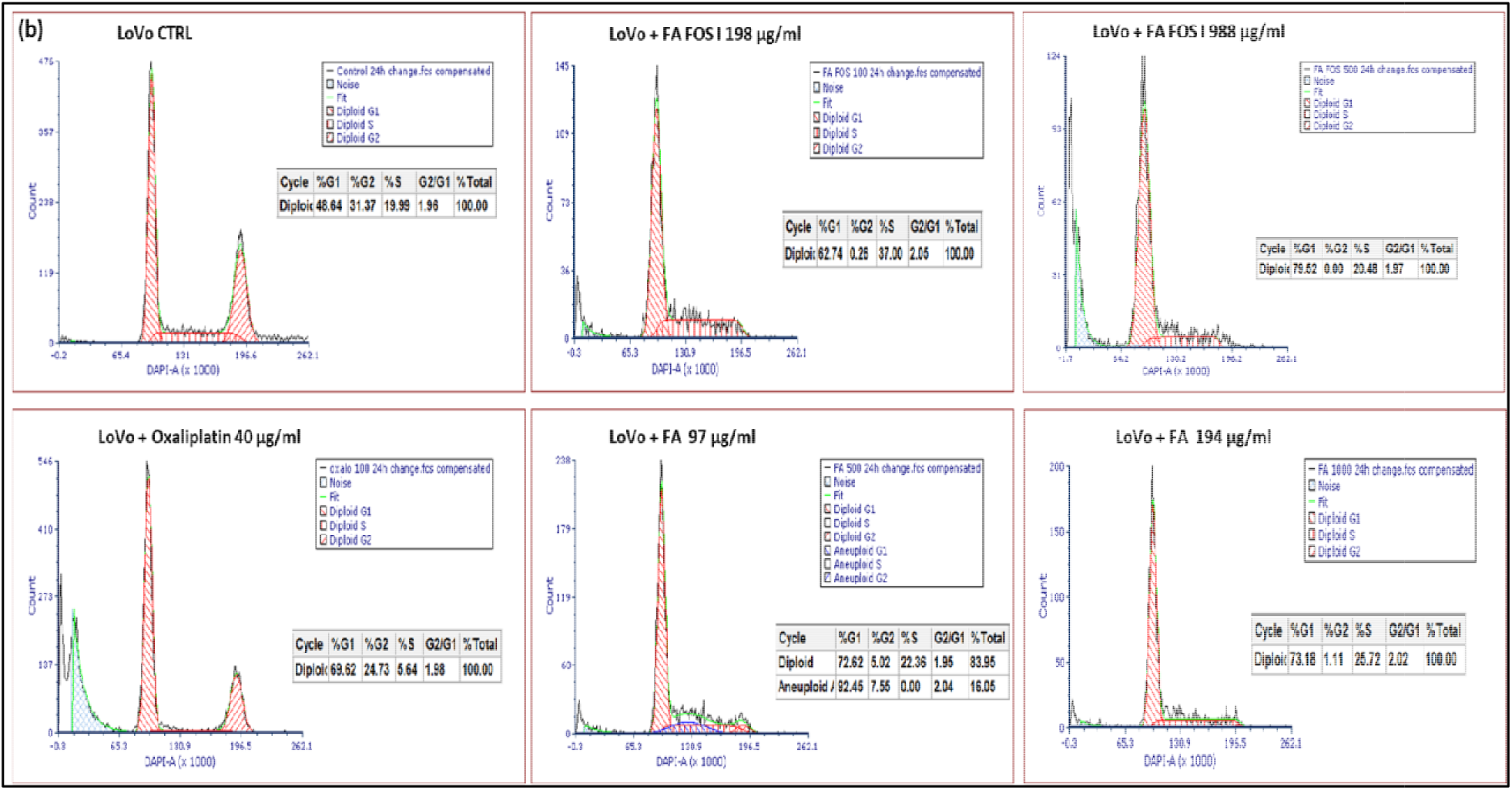
**a** Determination of cell cycle stage in HT-29 cell line by using FACS analysis of cellular DNA content **b** Determination of cell cycle stage in LoVo cell line by using FACS analysis of cellular DNA content

To further validate the process of cell cycle arrest expression profiling of cyclins B1, E and cyclin dependent kinase inhibitor protein p21 is determined by using the total protein extract from LoVo cell line treated as in the above condition and densitometry analysis were performed by using Image J software.

**Fig. 11.**
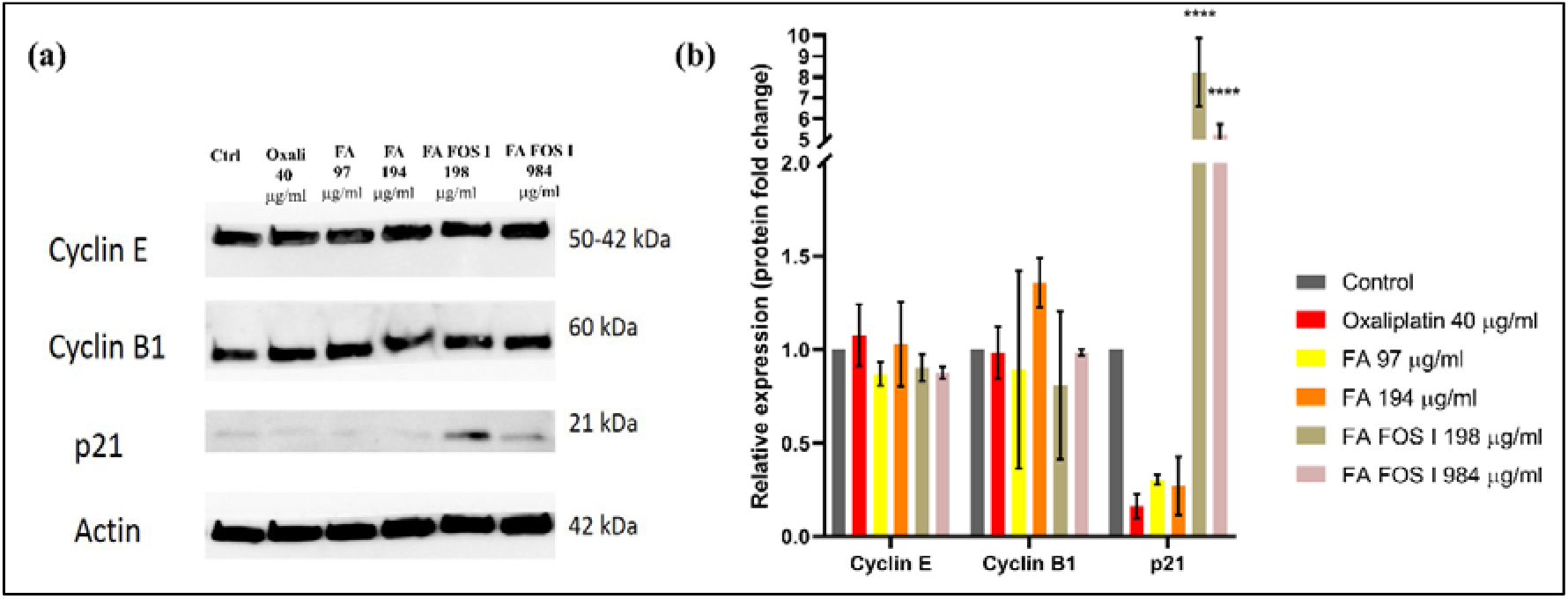
(a) Western blot of the total protein fraction of the treated LoVo cell, (b) relative expression of cyclin B1, (c) relative expression of cyclin E, (d) relative expression of p21 over control (un-treated sample).

It was observed from the bivariate analysis of DNA content and western blot analysis that FA FOS I induce cell cycle arrest by up-regulating the expression of p21 which inhibits all the cyclins which regulate the cell cycle and down regulation of cyclin B1 which are involved in the progression of cells from G_2_ phase to mitosis and a slight down regulation of Cyclin E which are involved in the progression of the cells from G_0_/G_1_ phase to S phase thus locking in the cells at G_1_ – S phase and preventing further progression of the cell cycle hindering the proliferation of the cancer cells.

### 3.5 Gene expression profiling of cell cycle related genes on treatment with FA FOSI

It has been observed that there is an upregulation of 11 key genes and a slight downregulation of 04 key genes which are directly involved in the cell cycle progression and proliferation.

The gene expression profile suggests that the compound FA FOS I causes DNA damage in the cells by double stranded and single stranded breaks which is acted upon by BRACA1 tumour suppressor protein along with other tumour suppressor proteins like BRCA2 which attempts the single stranded DNA repair and forms a complex with its cofactor BCCIP and modulates cyclin dependent kinase 2 via p21 tumour suppressor, thereby stalling the cell proliferation at the G_1_ phase (41). The Ataxia telangiectasia and Rad3-related protein (ATR) is also activated in response to single strand breaks by sensing DNA damage and activates DNA damage checkpoint leading to cell cycle arrest. ATR is also related to the second checkpoint-activating kinase ATM (serine/threonine protein kinase) which responds to double strand breaks in DNA and chromatin disruption which activates DNA damage checkpoint leading to cell cycle arrest, DNA repair or apoptosis (42). The cyclin G2 (CCNG2) are involved in growth regulation and in negative regulation of cell cycle progression, it’s under expression or repression was involved in un-controlled cell proliferation, migration and invasion. The cyclin T1 protein (CCNT1) tightly regulates the Cyclin Dependent kinases (CDK) and especially is highly associated with CDK9 kinase which is a major subunit in the transcription elongation factor p- TEFb. The cyclin dependent kinase CDK6 plays an important role in the accumulation of apoptosis proteins p53 and p130 which stops the cells from entering cell division in the presence of DNA damage at G1 phase by activating pro-apoptotic pathways. CDK6 interacts with CD1, CD2 and CD3 and their complex acts as a point of switch to commit to cell division in response to external signals like mitogen or growth factors (43). The cyclin-dependent kinase inhibitor (CDI) protein are proteins which inhibits cyclin-dependent kinase, several of which acts as tumour suppressor proteins. Cell progressions are negatively controlled by such kinase inhibitors, which arrest the cell cycle at G_1_ phase. One such CDI is CDKN2B which is a cyclin-dependent kinase 4 (CDK4) inhibitor B also known as multiple tumour suppressor 2 (MTS-2) or p15. They form complex with CDK4 or CDK6 and prevents the activation of CDK kinases by cyclin D thus functions as a cell growth regulator which inhibits the cell cycle G_1_ progression (44). The CDK5 regulatory subunit-associated protein 1 (Cdk5rap1) is the regulator of CDK5 activity, they inhibit CDK5 function which is involved in cell proliferation in invasive cancers by apparently reducing the activity of the actin regulatory protein caldesmon, thus preventing cancer cell migration and invasion (45). The anti-apoptotic BCL-2 family protein play an important role in transcription-independent apoptosis of p53 by binding to the transactivation domain p53TAD which is a highly conserved mechanism, shared among Bcl-XL/Bcl- 2, MDM2 and CBP/p300 (24).

The DNA replication licensing factor MCM4 is a protein which is one of the highly conserved mini-chromosome maintenance proteins (MCM) that are essential for the initiation of eukaryotic genome replication in the S phase of the cell cycle (46). This gene expression was down regulated 1.13-fold on FA FOS I treatment for 24h. SERTA domain-containing protein 1 is a protein which renders activity to cyclin D1/CDK4 inhibited by CDI like CDKN2A/p16INK4A and imparts a positive regulation of cell proliferation, this gene expression was down-regulated 1.13-fold on treatment with FA FOS I which might be responsible for stalling of the cell cycle at G1/S transition. The CDC34 is a gene which encodes for protein which has a ubiquitin conjugating activity, they are a part of the large multiprotein complex which are required for ubiquitin-mediated degradation of cell cycle G1 regulators involved in the initiation of DNA replication (47), this gene was found to be slightly downregulated on treatment with FA FOS I. The E2F1 belongs to the E2F family of transcription factors coordinating the expression of key genes involved in the cell cycle regulation and progression. They are mainly active during G1 to S transition, which includes the regulatory elements of the cell cycle such as CDC2, CDC25A and cyclin E and essential components of DNA replication machinery. In response to DNA damage E2F1 is stabilized by phosphorylation at S31 by ATM/ATR kinases (which are upregulated by treatment with FA FOS I) which is an integral component of the DNA damage signaling pathway. In response to agents which causes DNA double strand breaks this phosphorylation by ATM/ATR seems to prime E2F1 for acetylation at specific lysine residues, these acetylations are a prerequisite for targeting of the P73 gene promoter by E2F1 which ultimately leads to apoptosis (48).

The gene expression data were fed to the Reactome database, which revealed the significant pathways as follows: (i) RB1 binding to E2F1, (ii) Regulation of mitotic G1/S transition due to RB1, (iii) Tp53 regulates the transcription of DNA repair genes, (iv) Activation of ATR in response to replication stress, (v) G2/M checkpoint and inhibition of cell cycle.

RB1 protein, also known as pRB or retinoblastoma protein, is a nuclear protein that plays a major role in the suppression of G1/S transition during mitotic cell cycle in multicellular eukaryotes. RB1 performs this function by binding to activating E2Fs (E2F1, E2F2 and E2F3), and preventing transcriptional activation of E2F1/2/3 target genes, which include several genes involved in DNA synthesis. RB1 binds E2F1/2/3 through the so-called pocket region, which is formed by two parts, pocket domain A (amino acid residues 373-579) and pocket domain B (amino acid residues 640-771). Besides intact pocket domains, RB1 requires an intact nuclear localization signal (NLS) at its C-terminus (amino acid residues 860-876) to be fully functional(49).

**Fig. 12.**
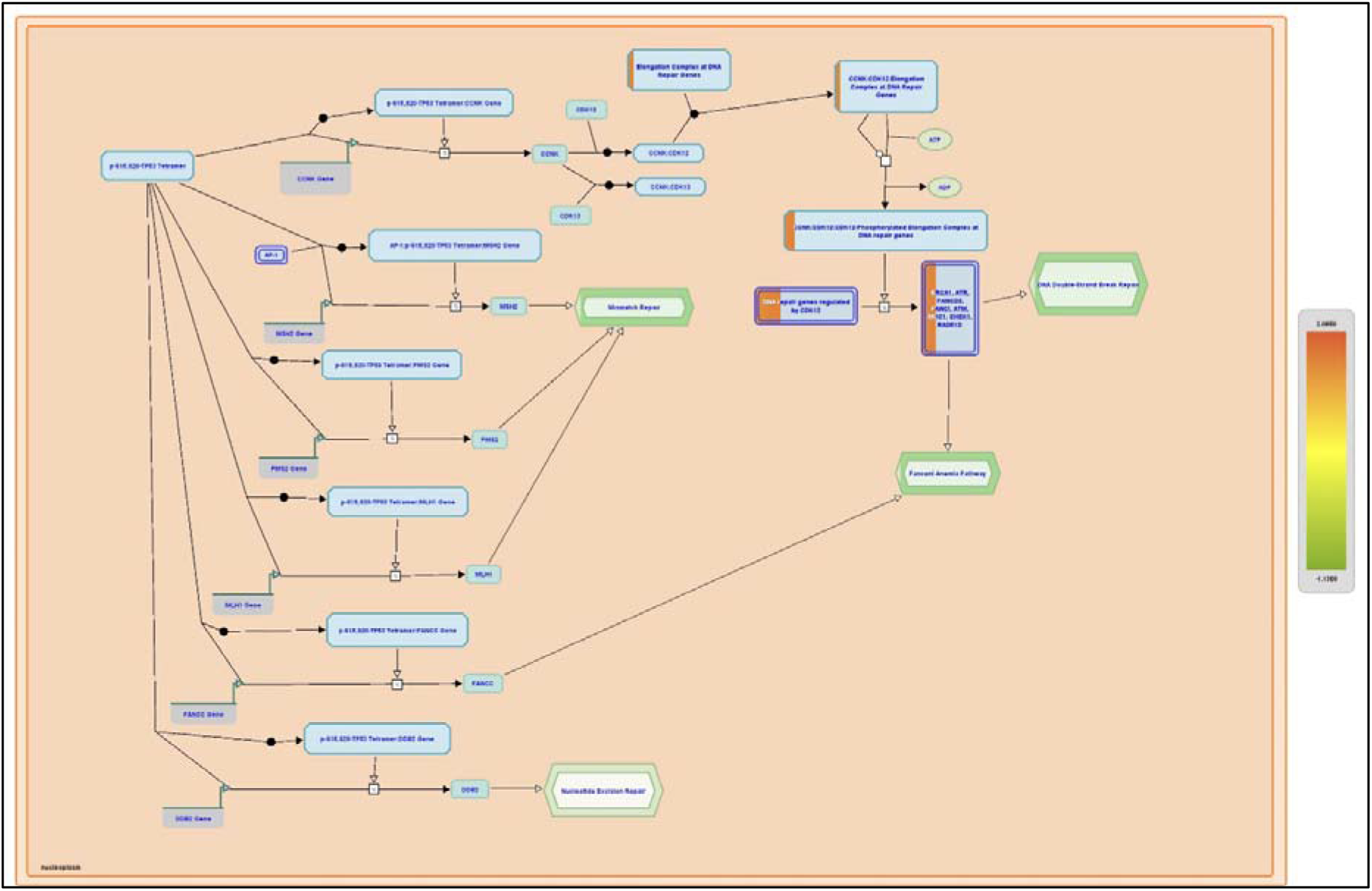
Regulation of the transcription of DNA repair genes by TP53 from reactome pathway analysis

Several DNA repair genes contain p53 response elements and their transcription is positively regulatedby TP53 (p53). TP53-mediated regulation probably ensures increased protein level of DNA repair genes under genotoxic stress. TP53 directly stimulates transcription of several genes involved in DNA mismatch repair, including MSH2, PMS2 and MLH1. TP53 also directly stimulates transcription of DDB2, involved in nucleotide excision repair, and FANCC, involved in the Fanconi anemia pathway that repairs DNA interstrand crosslinks. Other p53 targets that can influence DNA repair functions are RRM2B, XPC, GADD45A, CDKN1A and PCNA. Contrary to the positive modulation of nucleotide excision repair (NER) and mismatch repair (MMR), p53 can negatively modulate base excision repair (BER), by downregulating the endonuclease APEX1 (APE1), acting in concert with SP1 (49). Expression of several DNA repair genes is under indirect TP53 control, through TP53-mediated stimulation of cyclin K (CCNK) expression. CCNK is the activating cyclin for CDK12 and CDK13. The complex of CCNK and CDK12 binds and phosphorylates the C-terminal domain of the RNA polymerase II subunit POLR2A, which is necessary for efficient transcription of long DNA repair genes, including BRCA1, ATR, FANCD2, FANCI, ATM, MDC1, CHEK1 and RAD51D. Genes whose transcription is regulated by the complex of CCNK and CDK12 are mainly involved in the repair of DNA double strand breaks and/or the Fanconi anemia pathway (50).

Genotoxic stress caused by DNA damage or stalled replication forks can lead to genomic instability. To guard against such instability, genotoxically stressed cells activate checkpoint factors that halt or slow cell cycle progression. Among the pathways affected are DNA replication by reduction of replication origin firing, and mitosis by inhibiting activation of cyclin-dependent kinases (Cdks). A key factor involved in the response to stalled replication forks is the ATM- and rad3-related (ATR) kinase, a member of the phosphoinositide-3-kinase-related kinase (PIKK) family. Rather than responding to lesions in DNA, ATR and its binding partner ATRIP (ATR-interacting protein) sense replication fork stalling indirectly by associating with persistent ssDNA bound by RPA. These structures would be formed, for example, by dissociation of the replicative helicase from the leading or lagging strand DNA polymerase when the polymerase encounters a DNA lesion that blocks DNA synthesis. Along with phosphorylating the downstream transducer kinase Chk1 and the tumor suppressor p53, activated ATR modifies numerous factors that regulate cell cycle progression or the repair of DNA damage. In response to replication stress, Chk1 phosphorylates CDC25B and CDC25C leading to Cdc25B/C complex formation with 14-3-3 proteins. As these complexes are sequestered in the cytoplasm, they are unable to activate the nuclear CDK1-cyclin B complex for mitotic entry. Persistent single-stranded DNA associated with RPA binds claspin and ATR : ATRIP, leading to claspin phosphorylation. In parallel, the same single stranded DNA: RPA complex binds RAD17: RFC, enabling the loading of RAD9 : HUS1: RAD1 (9-1-1) complex onto the DNA. The resulting complex of proteins can then repeatedly bind and phosphorylate CHK1, activating multiple copies of CHK1 (51). G2/M checkpoints include the checks for damaged DNA, un-replicated DNA, and checks that ensure that the genome is replicated once and only once per cell cycle. If cells pass these checkpoints, they follow normal transition to the M phase. However, if any of these checkpoints fail, mitotic entry is prevented by specific G2/M checkpoint events. The G2/M checkpoints can fail due to the presence of un-replicated DNA or damaged DNA. In such instances, the cyclin-dependent kinase, CDC2(Cdk1), is maintained in its inactive, phosphorylated state, and mitotic entry is prevented. Events that ensure that origins of DNA replication fire once and only once per cell cycle are also an example of a G2/M checkpoint. In the event of high levels of DNA damage, the cells may also be directed to undergo apoptosis (52).

### 3.6 FA FOS conjugates prevents AOM-DSS mediated colitis-associated tumorigenesis in mice model

It was observed that there was a 60.83 % (p < 0.001) reduction in FA FOS I treated group compared to control which was significantly higher tumour reduction compared to Oxaliplatin which showed only 43.5 % reduction (p<0.005). FA FOS I has much higher efficiency (p< 0.01) than FA administration which showed only 36.4% reduction (p< 0.01) compared to control. There was no significant difference in the colon index of mice among the control group and the treatment group.

There was no statistically significant difference in the body weight of mouse after four weeks of treatment when compared to the control group, while there was a reduction of 18.46% in the body weight of FA FOS I treated group (p<0.001) when compared to naïve group. There was a reduction in 14.43% in the body weight of oxaliplatin treated group (p<0.01) when compared to naïve group. There was no statistically significant difference in the body weight gain rate of mice compared to AOM DSS control. There was no statistically significant difference in the length of the colon in the treatment groups compared to AOM DSS control and the naïve group (Fig. 13).

**Fig. 13.**
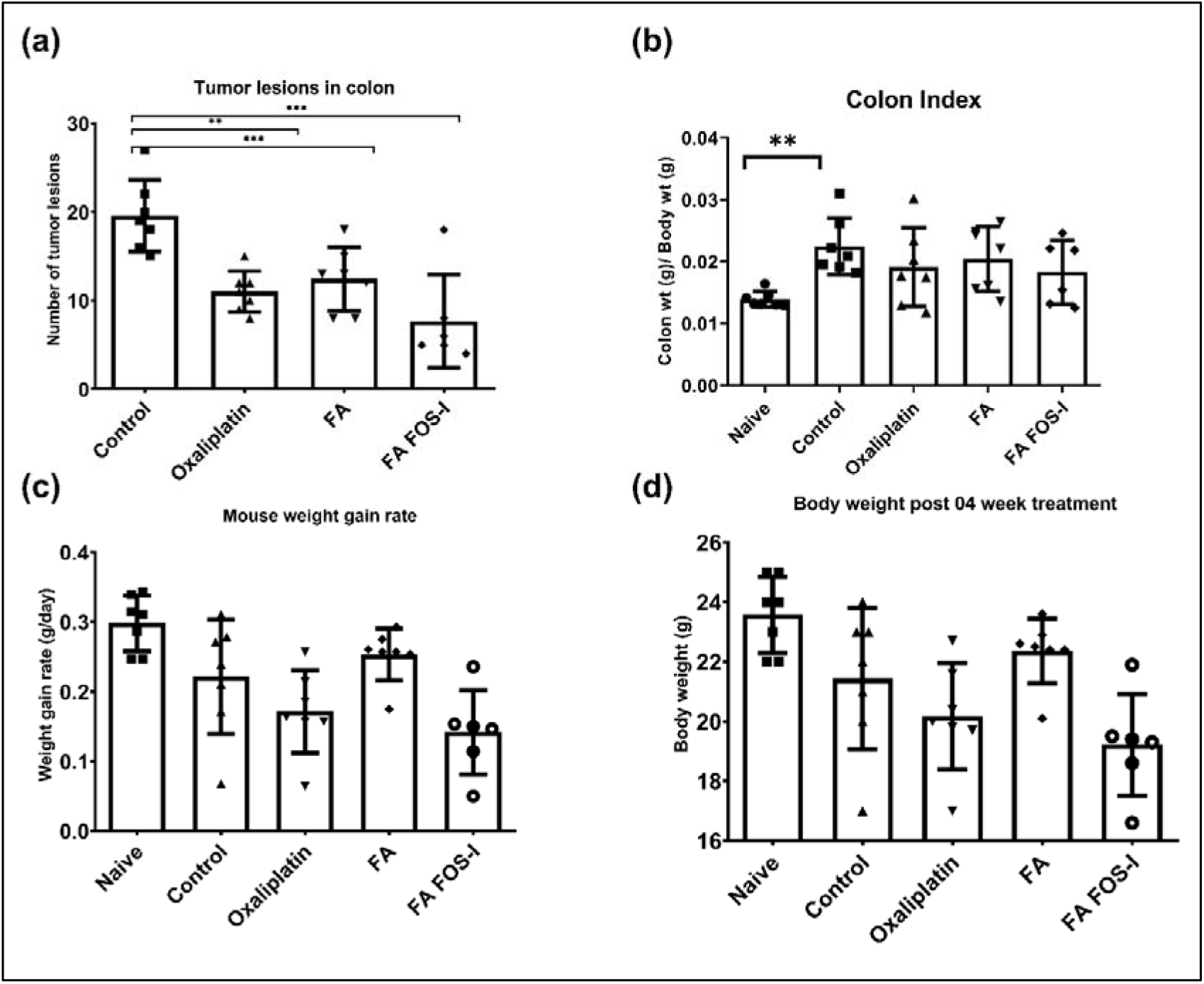
Macroscopic parameters in AOM DSS induced colitis associated cancer model (a) Number of tumour lesions in the colon/polyp per colon, (b) Colon index: the ratio of weight of colon to body weight of mice in different treatment group, (c) The weight gain rate of mice in each treatment group, (d) The final body weight of mice after four weeks of administration of drug/vehicle control. Data was acquired from three independent experiments and is expressed as a mean (n=6) ±SD. * p<0.01, **p<0.001, ***p<0.0001 vs control group.

**Fig. 14.**
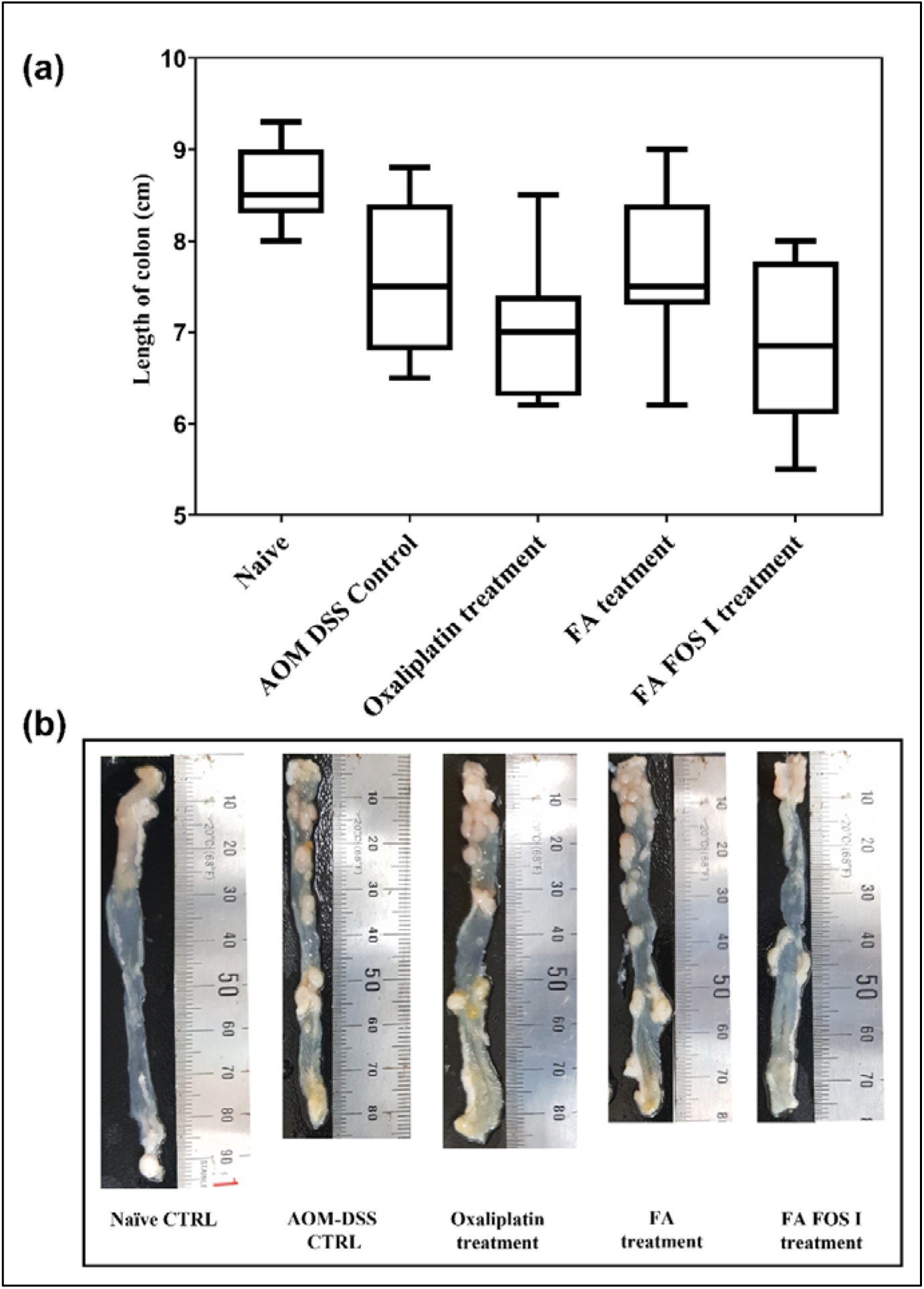
**(a)** Average length of the colon of Naïve and AOM-DSS mediated colitis associated colon cancer mice treated for four weeks with the indicated compound/vehicle control. Data was acquired and is expressed as a mean (n=6) ±SD.

**Fig. 15.**
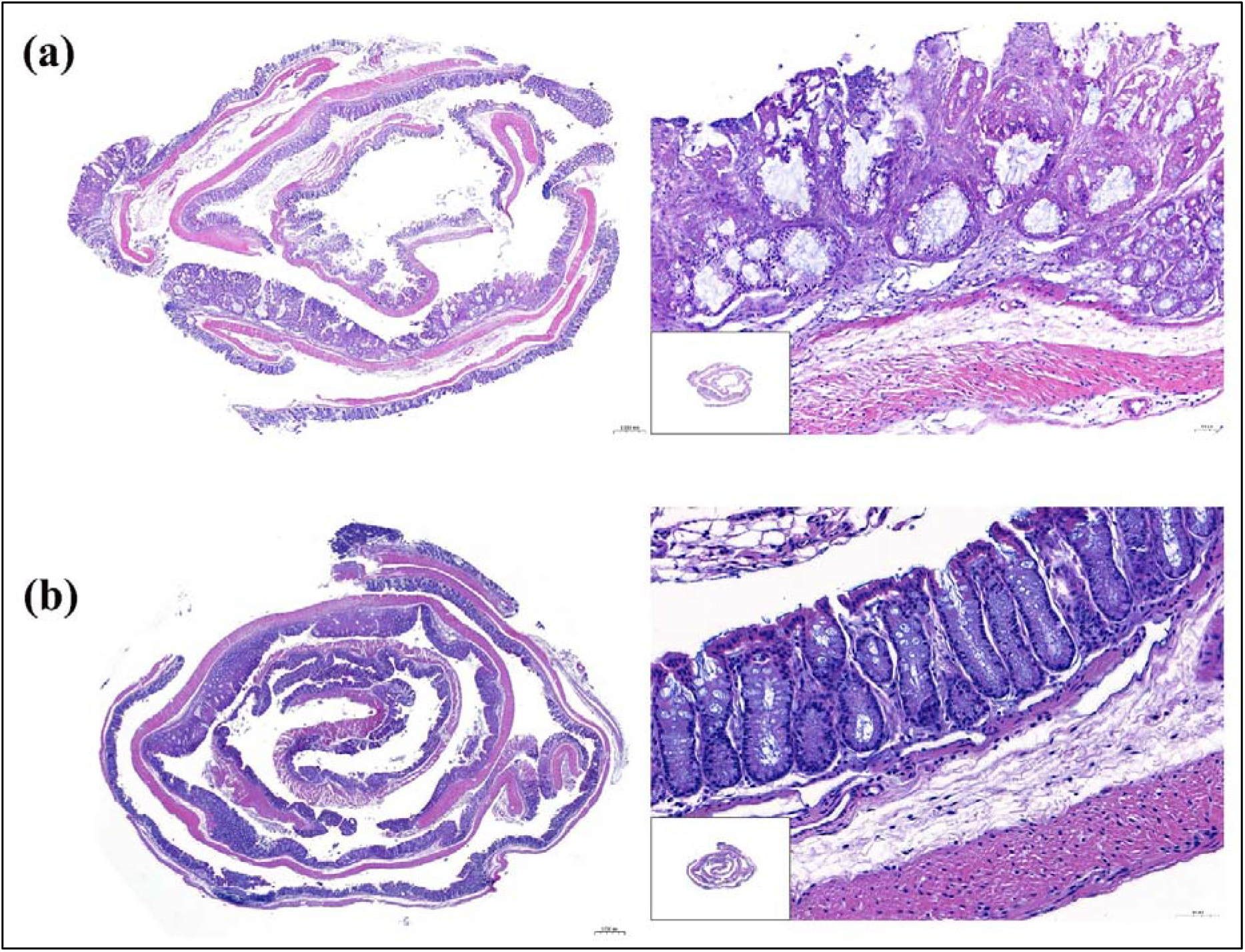
The representative H&E stained swiss-roll whole colon tissue of AOM DSS mediated colitis induced colon cancer mice model at 40x, Scale bar =0.5 mm, magnified region to the right, 200x, Scale bar =50 µm (a) Colitis associated colon cancer control, (b) FA FOS I 500 mg/Kg treatment group

The H&E histopathology of the colon of the mice showed that there was a predominant destruction of the intestinal lining in the AOM DSS mediated colitis associated colon cancer (CAC) mice leading to the compromise of the intestinal barrier and tight junction, in Fig. 12 (a) at several places of the colon the stroma has invaded into the mucosa and submucosa region and the arrangement of columnar epithelial cells were greatly distorted. The mucin layer was also found to be diminished and damage to goblet cells and crypt cells were found in the control intestinal tissue (Fig.17 (b)) compared to the naïve group. In the FA FOS I treatment group all these abnormalities were ameliorated to a significant extent on four weeks of oral administration. The H&E scoring of the Swiss roll colon tissue of the mice revealed that the percentage of tumour/malignant cells in the intestine of vehicle control group was 53.41% while the FA FOS I treatment group showed 27.62% tumour/malignant cell population which is 48.27% reduction for FA FOS I treatment group compared to the vehicle control group. The area occupied by the malignant cells were also drastically reduced in the FA FOS I treatment group (Fig. 16).

**Fig. 16.**
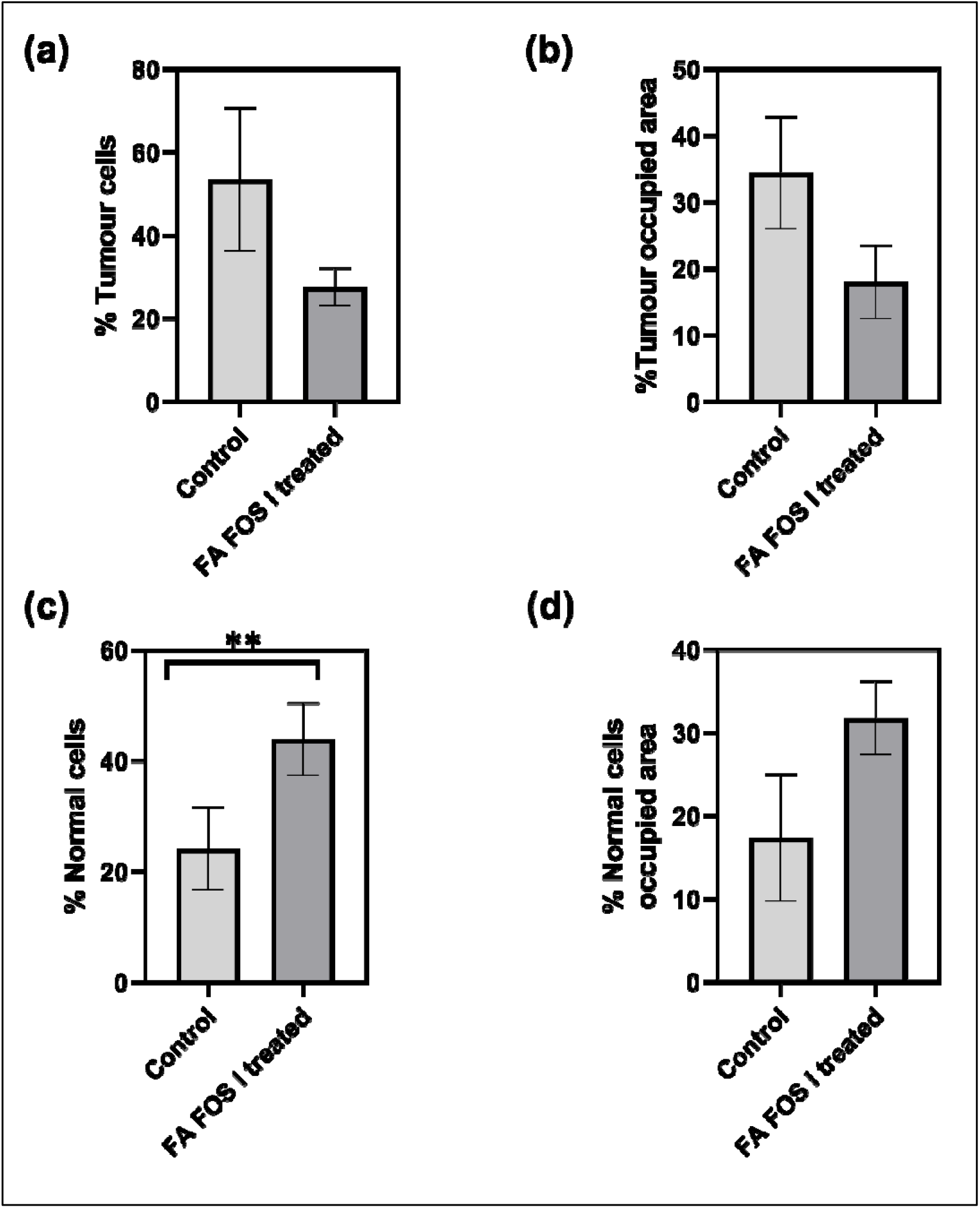
The proportion of tumour/malignant cells and normal cells determined from the H&E-stained tissue sections (histopathology) determined by using Qpath. (a) Percentage of tumour cells, (b) Percentage of area occupied by tumour cells, (c) Percentage of normal cells, (d) Area occupied by normal/non-malignant cells. Data was acquired from three independent experiments and is expressed as a mean (n=3) ±SD. * p<0.01, **p<0.001, ***p<0.0001 vs control group.

**Fig. 17.**
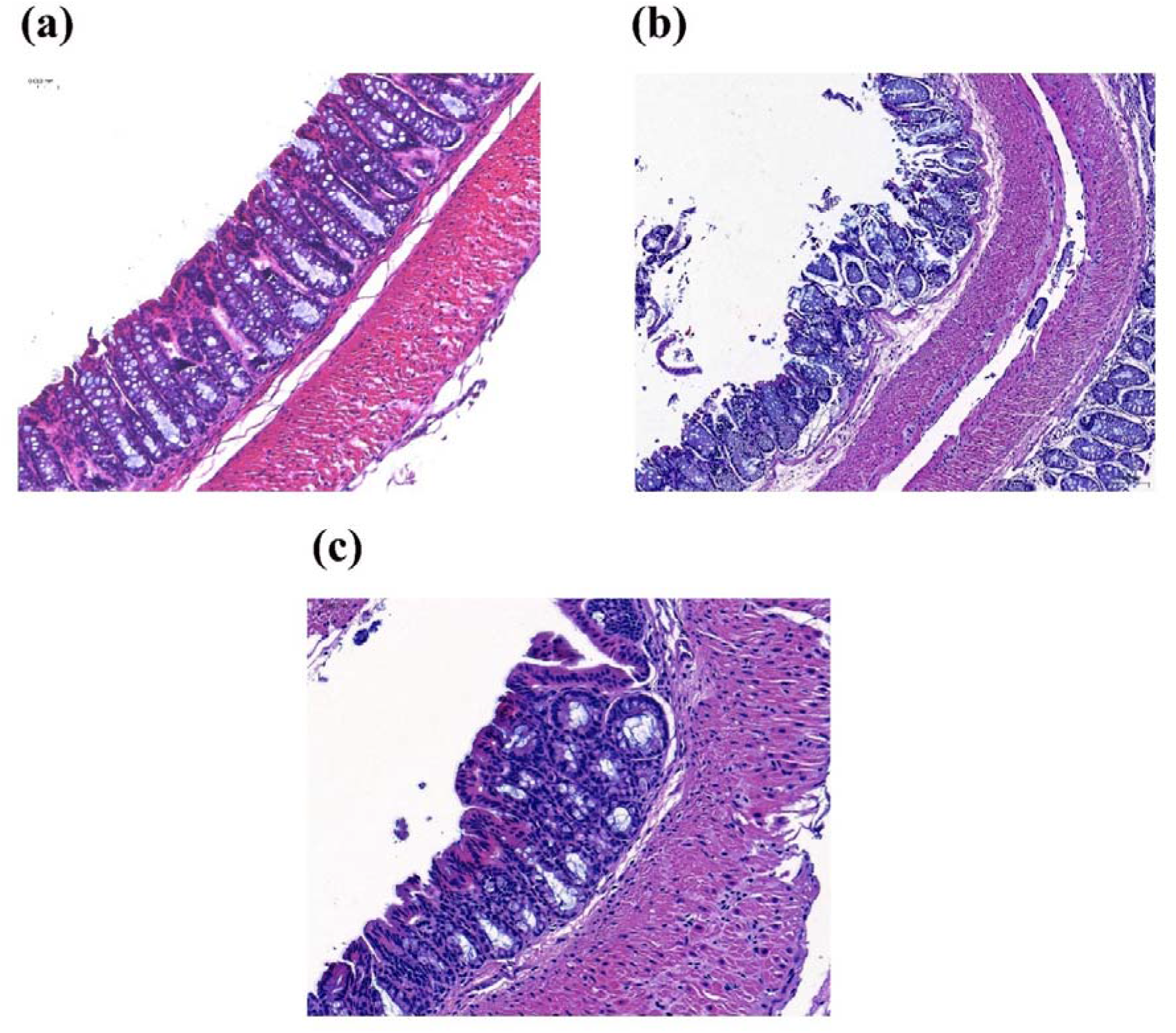
The histopathology of intestinal lining showing the mucin stained blue and neutral mucin stained pink-red, the goblet cells covered in mucin is also stained blue with Gill’s B hematoxylin and eosin-phloxine,200x, Scale bar = 0.050 mm (a) Naïve without colitis induced colon cancer, (b) Colitis associated colon cancer mouse model showing disrupted intestinal barrier, where the integrity of the columnar epithelial cells are lost showing irregular arrangement and invasion of the stoma in to the mucosa and submucosa. (c) Colitis associated colon cancer (CAC) mouse treated with FA FOS I for 4 weeks, they show intact and functional mucosal barrier and intestinal lining with uniform margin and intact goblet cells in blue producing mucin. They show proper intestinal integrity with tight junction, the submucosa and serosa does not show any sign of invasion.

The histological analysis of colon tissues from CAC control showed severe inflammation, destruction of the crypts, superficial ulceration, damage to the mucosa and infiltration of leucocytes to the epithelium, lamina propria and submucosa, whereas the administration of FA FOS I was found to be alleviating the colonic damage, inflammation, and epithelial damages in the colonic tissues (Fig. 54 and 56). These clearly reveal that FA FOS administration reduced the pathological symptoms caused by AOM/DSS which also includes reduction of glandular hyperplasia, extinction of nuclear polarity, hyperchromatic nucleus and enlarged nucleoplasmic ratio (Fig 54).

### 3.7 FA FOS conjugates regulates CDX2-**β** Catenin axis in ameliorating AOM/DSS mediated CAC colorectal cancer model

The cyclooxygenase-2 (COX-2) are considered to be the prime enzyme, epoxide hydrolase, necessary for the synthesis of prostaglandins (PGs), they play an important role in the induction of inflammation and in tumorigenesis, many clinical studies have found that high expression of COX-2 is an early event in the development of tumorigenesis in human colorectal adenoma and adenocarcinomas (53). In this study it was found that the expression of COX-2 was elevated in CAC control but was lowered in FA FOS I treated cells. The CAC vehicle control group showed 9.84% ±1.71 positive cells vs 4.45% ± 1.2 for FA FOS I which was 54.7% reduction in positive cells compared to CAC vehicle control. β-Catenin plays a significant role in Wnt/β-catenin signal transduction system and the nuclear accumulation of β-Catenin is associated with colon carcinogenesis (54). Reports suggests that constitutive activation or deregulation of Wnt/β-catenin pathway has been known to contribute to the genesis and progression of different kind of disorders in humans which include early-stage sporadic colorectal cancer, irritable bowel disease (IBD) associated carcinogenesis etc., (55,56). Under normal conditions most of the β-Catenin are found bound to cadherin of the cytomembrane which are found to regulate cell to cell adhesion, where the β-Catenin present in the cytoplasm is rapidly phosphorylated by glycogen synthase kinase-3 beta (GSK3β) at Ser33, Ser37 and Thr41 residues by the adenomatous polyposis coli (APC), axis inhibition protein 2 (AXIN2), GSK-3β destruction complex and is finally degraded by the proteasome. In cancer cells the inactivation/mutation in the APC/AXIN/GSK-3β complex causes the accumulation of β-catenin in the cytosol followed by translocation into the nucleus, where it acts as a T-cell factor/lymphoid enhancer factor (TCL/LEF) mediated transcription co-activator and modulates cell survival, proliferation, and differentiation (55,56). In this study the CAC control group had 63.25% ±15.8 of cell population positive for nuclear localization of β-Catenin (Fig. 18 & 19), while the FA FOS I showed only 13.98 %±7.5 positive for β- Catenin nuclear localization which was a drastic reduction of 77.9% (p<0.01) for FA FOS I treatment group. The caudal-related homeobox transcription factor 2 (CDX2) is a prime transcription factor controlling the critical balance between cell proliferation and differentiation of the intestinal epithelium (57). It was reported that CDX2 is required for the proper cytodifferentiation and villus morphology of the intestinal lining (58–60). The decreased expression of CDX2 was often reported in invasive tumours of the colon and in tumour buddings/polyps (61,62). Since CDX2 gene are less prone to mutation it can be considered that a regulatory mechanism rather than genetic alteration contributes to the downregulation of CDX2. It was also acclaimed that key inflammatory transcription factor NF-κB is one of the chief regulators of CDX2 in colon cancer (63). CDX2 was also found to upregulate the expression of APC and AXIN2 (64,65) leading to the stabilization of the degradation complex of cytoplasmic β-catenin hence the possibility of elevated tumorigenicity of CAC caused by the down-regulation of CDX2 and activation of β- catenin is undeniable. In this present research the AOM/DSS mediated CAC control showed very low expression of CDX2 while the treatment with FA FOS I upregulated the expression of CDX2 13.4- fold (p<0.01). It has been reported that CDX2 plays a role in the regulation of the expression of APC and AXIN2 the components of β-catenin degradation complex (66) and also the increasing experimental evidences supports the notion that CDX2 inhibits β-catenin and thereby the progression of colon cancer (67,68), the loss CDX2 was also reported to be observed in invasive colorectal cancers (69,70). The present study also supports the fact that CDX2 over expression indeed reduced the expression of β-catenin and reduced the tumour proliferation. The translocation of β-catenin from cytoplasm to nucleus was reported to serve as an transcription factor for the activation and upregulation of its downstream target genes Cyclin D1 and C-Myc (71). Early reports suggests that in the process of development of colitis associated colon cancer, the upregulation of TNF-α induces the phosphorylation of Iκb-α and nuclear localization of p65 which inhibits the expression of CDX2 which inactivates the destruction complex of β-catenin and blocks the phosphorylation of β-catenin at Ser37 causing the β-catenin induced expression of cell proliferation signal cascade involving PCNA, Cyclin D1 and c-Myc promoting the CAC tumour development (72). In agreements with the previous reports in our study the expression of Cyclin D1 and c-Myc in the FA FOS treatment was reduced compared to CAC control group. The CAC control group showed an elevated Cyclin D1 and c-Myc which showed 51.36% ±12.7 cell population positive for Cyclin D1 and 56.98% ±0.9 cell population positive for c-Myc. While the FA FOS I and II treatment group showed and 92.36% reduction and 87.8% reduction (p<0.001) in population of Cyclin D1. The c-Myc for FA FOS I and II showed an 35.3% reduction and 40.8% reduction (p<0.001) vs the CAC control group (Fig 19).

**Fig. 18.**
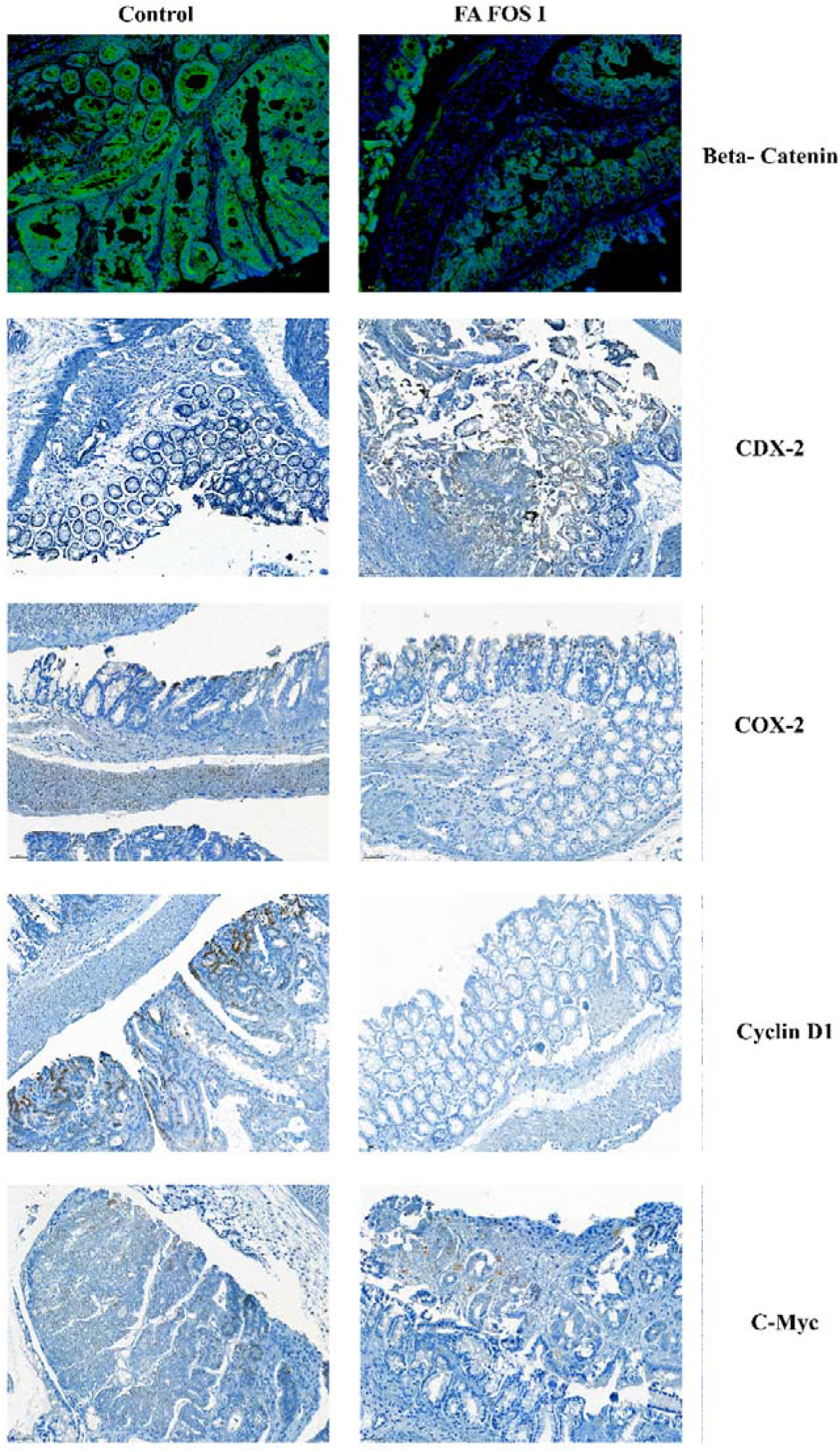
The photomicrograph of large intestine of mouse at x200, Scale bar= 50 µm, showing Immuno fluorescence (IF) of β-catenin localized (Green fluorescence) into nucleus stained with DAPI, Immuno histochemistry (IHC) of the intestine stained with anti-mouse CDX-2 antibody diluted 3200t, and counter stained with DAB. COX-2 antibody diluted 3200T, Cyclin D1 antibody diluted 800t, c-Myc antibody diluted 400t.

**Fig. 19.**
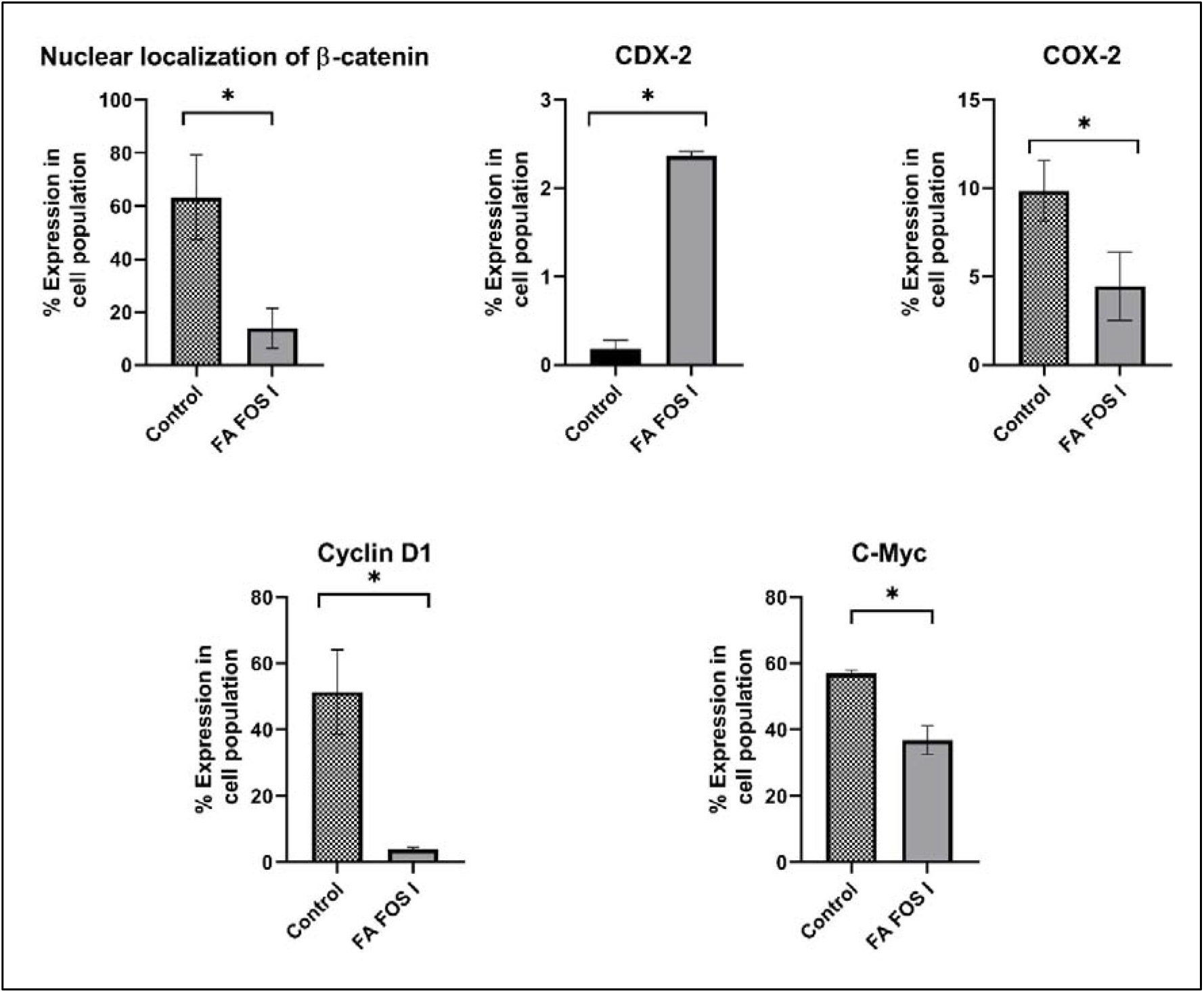
Scoring of the IF and IHC histopathological slides using Qpath, the bar graph shows the proportion of positive staining of cells among the total cell nuclei detected. Data was acquired from three independent experiments and is expressed as a mean (n=3) ±SD. * p<0.01, **p<0.001, ***p<0.0001 vs control group.

### 3.8 FA FOS conjugates inhibits the proliferation and enhanced apoptosis of the tumour cells in AOM DSS CAC model

The proliferation of cell is an important function in the growth, repair, and normal functioning of an organism but the un-checked proliferation of the malignant cells will lead to tumorigenesis. Several reports suggests that the aberrant proliferation of colorectal cells would lead to the development of neoplasm (73,74). In the event of cell proliferation the proliferative markers Ki67 and PCNA are highly expressed in the colorectal cancer tissues (75,76). In our study the cell proliferation marker Ki67, PCNA were highly expressed throughout the tumours in the CAC control group, but FA FOS conjugate treatment drastically reduced the nuclear expression of the proliferation markers. Nuclear Ki67 positive cell population in the CAC control group was 79% ±6.1 vs 43.77% ±4.02 for FA FOS I treatment group. In the PCNA marker the percentage of cells positive in CAC control group were 88.32% ±2.12 vs 54.72% ±3.2 for FA FOS I. Apoptosis which is described as programmed cell death is a cell stabilizing mechanism in organisms to regulate and cull the un-controlled proliferation of cells and play a pivotal role in the control of tumorigenesis and acts as an barrier for oncogenesis (77,78). The DNA fragmentation is considered as the hallmark of apoptosis and TUNEL staining is used to detect apoptotic progress. In the CAC control group, there were detected only 21.95% ±5.1 TUNEL positive cells while the FA FOS I treatment group showed 59% ±5.6 (p<0.001) TUNEL positive cell population among the malignant or cancerous cells in the intestine.

**Fig. 20.**
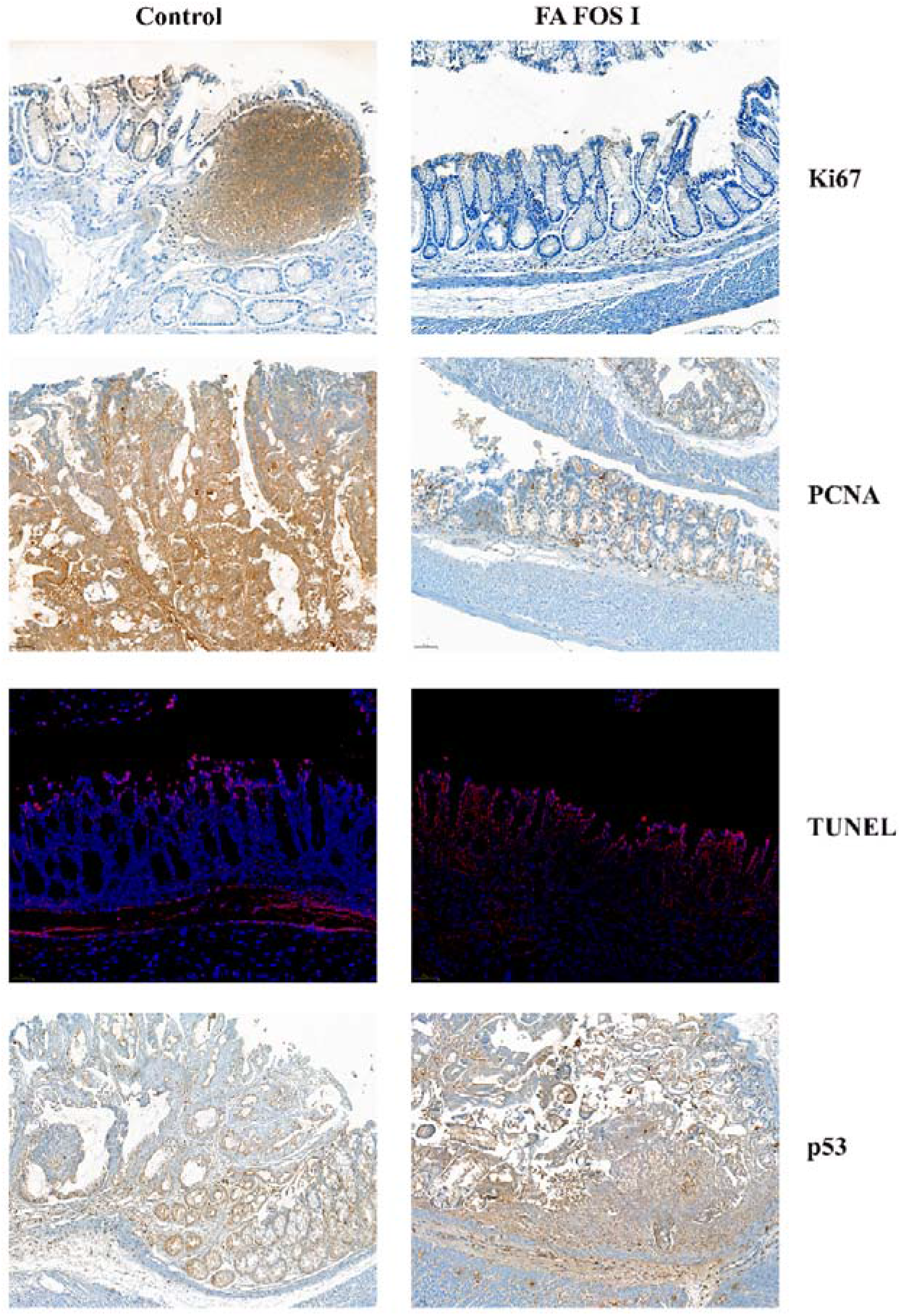
Photomicrograph of the large intestine of CAC mice treated with the indicated compounds or vehicle control for four weeks, 200x, Scale bar = 50 µm. IHC of proliferation marker antibodies Ki67 nuclear localization, antibody diluted 400t, nuclear expression of PCN, antibody diluted 51200t, IF of Terminal deoxynucleotidyl transferase dUTP nick end labelling (TUNEL) showing DNA fragmentation, nuclear expression of p53, antibody diluted 51200t.

**Fig. 21.**
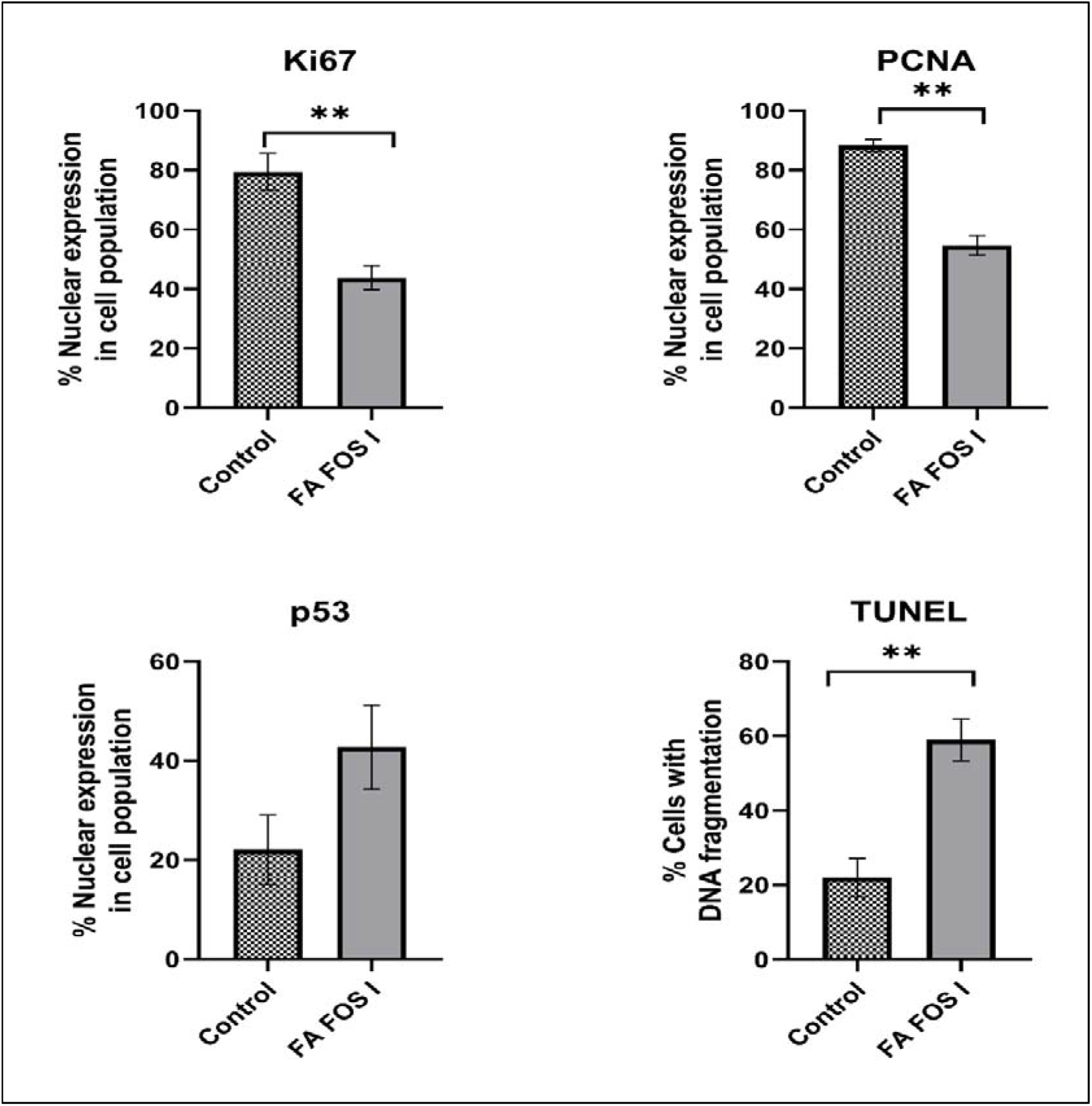
Scoring of the IHC histopathology slides using Qpath, the bar graph depicts the expression of the indicated antigens and labelling of fragmented DNA quantified with respect to the number of positive cells in the tissue section. Data was acquired from three independent experiments and is expressed as a mean (n=3) ±SD. * p<0.01, **p<0.001, ***p<0.0001 vs control group.

The functional p53 is considered to be a tumour suppressor and anti-oncogenic protein mediating apoptosis and considered as a central mechanism in the inhibition of tumour in colorectal cancers (527,528). It was found that the FA FOS conjugate treatment could considerably upregulate the expression of p53 in the cell nucleus which stood at an 93.7% (p<0.001) and 98.05% (p<0.001) enrichment of cells positive for nuclear expression of p53 marker vs CAC control group which showed an average of 22.05% ±7.03 cells positive for p53 among the tumour cell population.

### 3.9 FA FOS treatment ameliorates inflammation in colitis-associated colorectal cancer in mice model

Chronic inflammation of the gastrointestinal tract induced by poor dietary habits are found to be the main risk factors in colorectal cancers(79). Inflammation is found to initiate tumorigenesis by inducing DNA damage in the absence of carcinogens attributed by increased oxidative stress caused by nearby tissues of the sight of inflammation or by the recruited immune system such as macrophages and neutrophils which release high levels of reactive oxygen and nitrogen species at the site of recruitment. The intestinal inflammation is also known to affect the epithelial barrier function and could expose the intestinal stem cells to environmental mutagen or inflammatory immune cells causing genotoxicity to the exposed stem cells and initiation of tumorigenesis (80). Nitric oxide (NO) is a short-lived regulator involved in various pathophysiological function and macrophage mediated immunity. Sustained induction of iNOS during the chronic inflammatory phase leads to reactive intermediates of NO which are mutagenic, alter cell signalling promoting proinflammatory properties contributing to carcinogenesis (81). iNOS are also found to regulate antiapoptotic molecules in tumour and enhance invasive and migratory properties. NO and its reactive metabolite peroxynitrite are involved in the stimulation of cyclooxygenase-2 (COX-2) enhancing the production of prostaglandins and aggravating tumorigenesis. In the present study the CAC control showed 6.96% ±1.35 cell population positive for iNOS vs 3.80% ±0.34 for FA FOS I which is a 45.36% (p<0.01) reduction. NF-κB activaction are reported to accelerate APC loss and colorectal cancer through the upregulation of iNOS (82). NF-κB are known to be the transcription factor regulating the inflammatory and immune responses modulating tumorigenesis most importantly in inflammation driven cancers (83). NF-κB are found to be activated in intestinal epithelial cells and in lamina propria macrophages and plays a pivotal role in development of CAC (84). In the intestinal epithelial cells, the pro-tumorigenic function of NF-κB is found to be mediated by its anti-apoptotic effects on the premalignant cells by obstructing its elimination, clearing and by the activation of β-catenin. Thus NF-κB are found to be acting as tumour promotor(84). In this study the CAC control group shows an elevated expression of NF-κB which accounts for 60.6% ±2.12 of the cells positive for NF-κB antigen seems to be migrating from the lamina propria on to the mucosa and sub mucosa region of the intestine (Fig. 22). There was found to be a reduction of up to 33.24% in reduction (p<0.001) of NF- κB positive cells in FA FOS I treatment group. The pro-inflammatory cytokine TNF-α is reported to be an prime mediator of inflammatory diseases and are found to activate signalling pathway involved in tumorigenesis (85). TNF-α are found to be activating Wnt/β-catenin pathway in colorectal cancer cells and in CAC mouse models (85). In the development of CAC the inflammatory cytokines such as TNF-α induces phosphorylation of IκB-α and found to promote the nuclear translocation of p65 inhibiting the protein expression of CDX2 which in turn inactivates the destruction complex of β- catenin. This cascade of events induces the upregulation of PCNA, Cyclin D1and c-Myc which further promotes development of CAC (85). In our study the expression of TNF-α was found to be elevated in CAC control group which had an 8.09% ±0.74 of cells positive for the cytokine while the treatment with FA FOS I are found to drastically reduce (p<0.01) the expression of the cytokine which was 4.58% ±0.70. It has been reported that the innate immune cells present in the stroma such as tumour associated macrophages play an major role in the tumour growth, neovascularization and invasion (86). The secretion of the pro-inflammatory cytokine TNF-α is elevated during acute inflammatory response and in sustained inflammation, where the levels of TNF-α is found to be increased in the colonic mucosa contributing to the invasiveness of adenocarcinoma (87). The TNF-α along with the activation of NF-κB pathway are involved in colorectal cancers (84) and reported to activate Wnt/β- catenin pathway in gastric tumour cells and in colitis-associated cancer (70). The pro-inflammatory cytokines IL-6 and IL-1β are found to be elevated in colorectal cancer and responsible for tumorigenesis (88,89). Reports suggests that the secretion of IL-6 by macrophages and T cells present in lamina propria is involved in the promotion of CAC in mice (90). It was reported that in AOM+DSS mediated mouse model of CAC the IL-6 was involved in trans-signalling in intestinal epithelial cells with downstream activation of STAT3 (91). STAT3 are found to be involved directly in the regulation of cell cycle progression and other mechanisms of inducing the expression of vascular endothelial growth factor receptor 2 (VEGFR2) in intestinal epithelial cells and enables an feedback loop (paracrine/auto) promoting the proliferation of tumour cells in CAC (92). In the present study the IL-6 was found to be elevated in CAC control group 6.91%± 1.30 vs FA FOS I which showed a decreased profile (p<0.01) with 4.55% ±0.21. IL-1β was found to be inducing cell proliferation in human colon cancer cells through NF-κB by repressing phosphatase and tensin homolog (PTEN) and induction of proliferation (93). Reports also suggest that the promotion of colon carcinogenesis by IL-1β was also mediated by inactivation of glycogen synthase kinase (GSK)3β leading to the activation of the Wnt pathway and tumour growth (94). In the present study the CAC control group showed and enhanced population of cells positive for IL-1β cytokine which depicted 9.39% ±0.72 of tumour cells in the mucosa and submucosal region while the FA FOS I showed 7.56% ±0.47 of cell population positive for IL-1β expression.

**Fig. 22.**
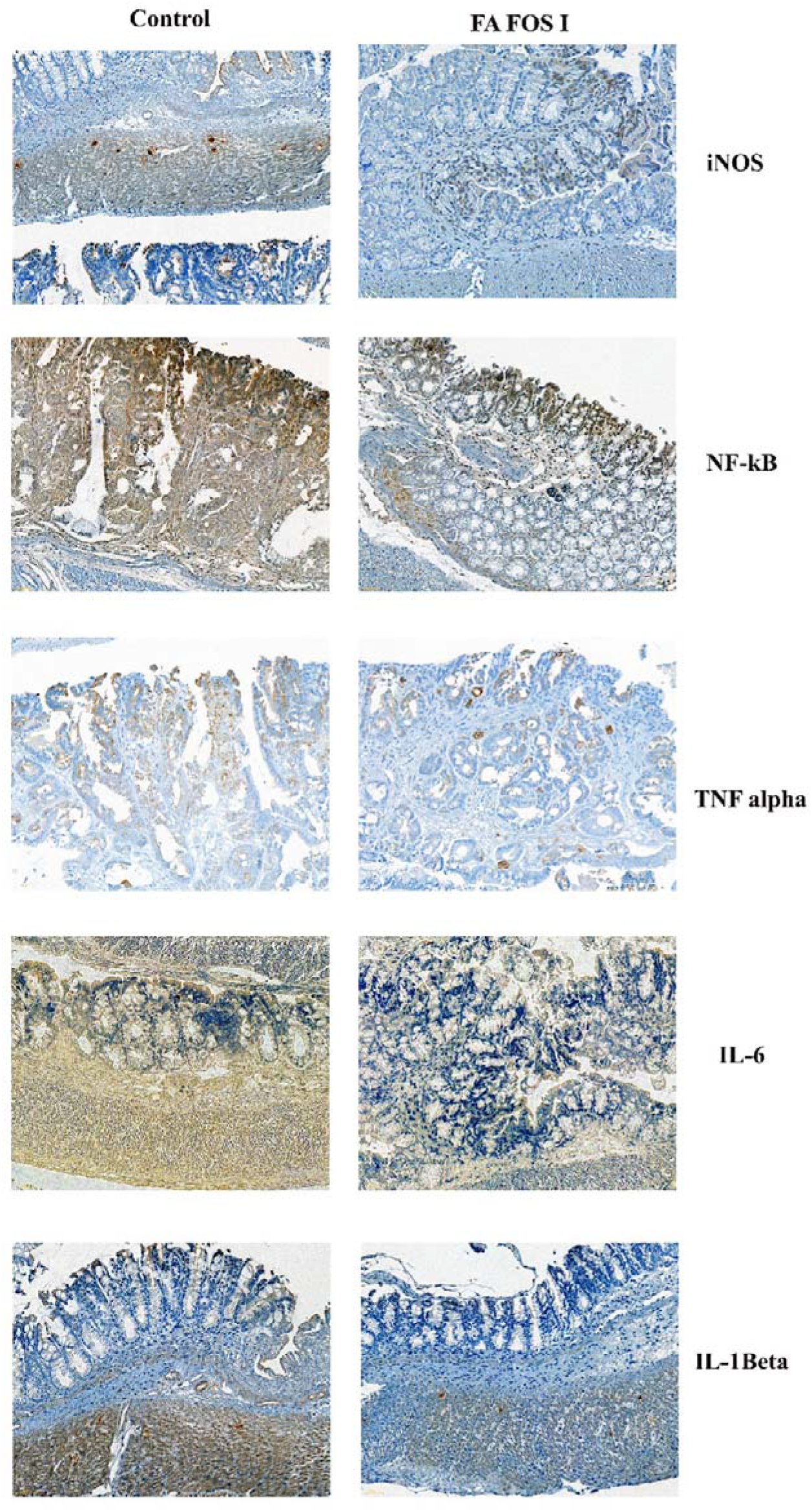
The photomicrograph depicts IHC of inflammation panel, the tissue sections are labelled with antibodies related to inflammatory response: iNOS anti-mouse antibody diluted 800t, NF-κB diluted 12800t, TNF α diluted 3200t, IL-6 diluted 3200t, IL-1β diluted 800t. Magnification 200x, Scale bar = 50 µm

**Fig. 23.**
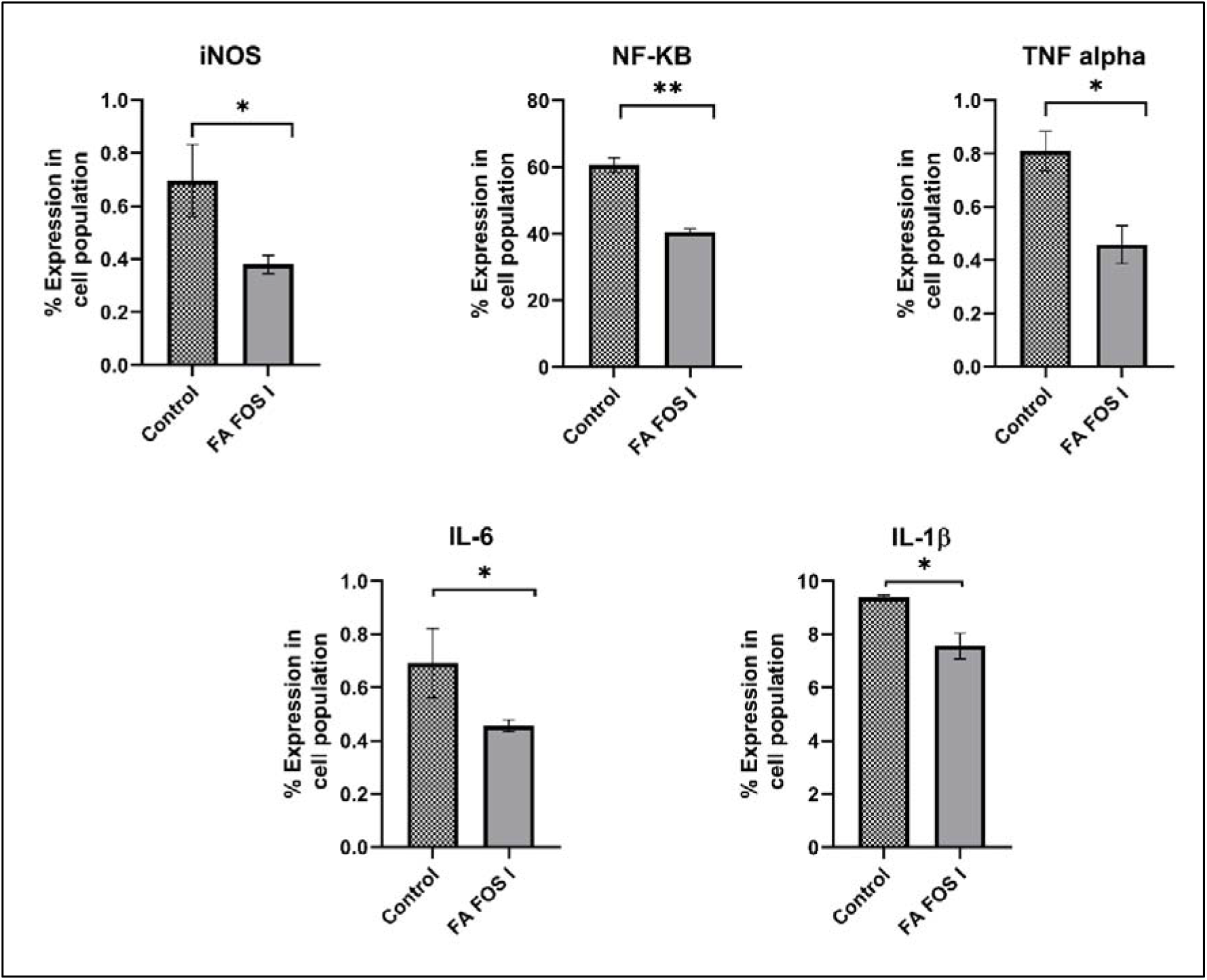
Scoring of IHC of the intestinal tissue with inflammation panel using Qpath, the bar graphs represent the proportion of cells labelled positive for the indicated antigen. Data was acquired from three independent experiments and is expressed as a mean (n=3) ±SD. * p<0.01, **p<0.001, ***p<0.0001 vs control group.

### 3.10 Immune modulation and induction of immune cells towards tumour microenvironment and sensitization of tumour cells to immune action by FA FOS conjugates

From the immunological parameters studied in the mice (Fig. 24) there was a 9% increase in the WBC count in FA FOS I group with a mean of 5.93x 10^3^ cells/ml ±0.216 (p<0.001) compared to control group which had 5.44± 0.20 x 10^3^ cells/ml. This may be due to the activation and recruitment of the immune cells to the tumour site for the eradication of cancer cells. There was no immune suppression induced by the compound FA FOS I and II which is evident from the bone marrow cell count however there was an 16.37% increase in the bone marrow cell count with a mean of 46.2 ±4.07 x10^5^ cells/ml in FA FOS I treatment group. The evaluation of thymus index and spleen index showed no statistically significant difference in the FA FOS I treatment group compared to the control group. In the case of IL-3 there was a 15.54% increase with a mean of 59.33 ±3.27 pg/ml (p<0.05) in the IL- 3 concentration present in the blood plasma of mice in FA FOS I treated group compared to the control group which had 51.35 ±2.11 pg/ml, showing a good prognosis towards tumour regression. IL-3 is a multipotent hematopoietic growth factor produced by activated T cells, macrophages and stroma cells and involved in the survival and proliferation of hematopoietic progenitor cells. They are the signal (cytokine) which improves the body’s natural defences against diseases as a part of immune system. They are produced by T cells only after stimulation with other antigens or other specific impulses, also they are known to have beneficial therapeutic effects in solid cancers in combination with chemotherapy(95,96). The IL-3 was also found to be involved in the stimulation and differentiation of immature myelomonocytic cells activating macrophages and granulocyte cell population (97). IFN-γ is an effector cytokine which plays a pivotal role in anti-cancer immunity by activation of CD4 T helper cells (type 1) and CD8 cytotoxic T cells, dendritic cells (DCs), macrophages, natural killer (NK) cells and aid in the antigen presentation (98). They are considered to be a pleiotropic molecule associated with antiproliferative, antitumour and pro-apoptotic mechanism. The administration of FA FOS I conjugate to the mice increased the plasma INF-γ levels in mice 19.56% (p<0.01) compared to AOM DSS CAC control group. The naïve group showed 63.2±8.27 pg/ml of INF-γ in the plasma while AOM DSS CAC group showed 50.6±3.89 pg/ml vs FA FOS I 60.5± 2.83 pg/ml, of INF-γ in the plasma. This evidence suggests the stimulation/sensitization of T cells and other immune cells to tumour antigens by FA FOS conjugate.

**Fig. 24.**
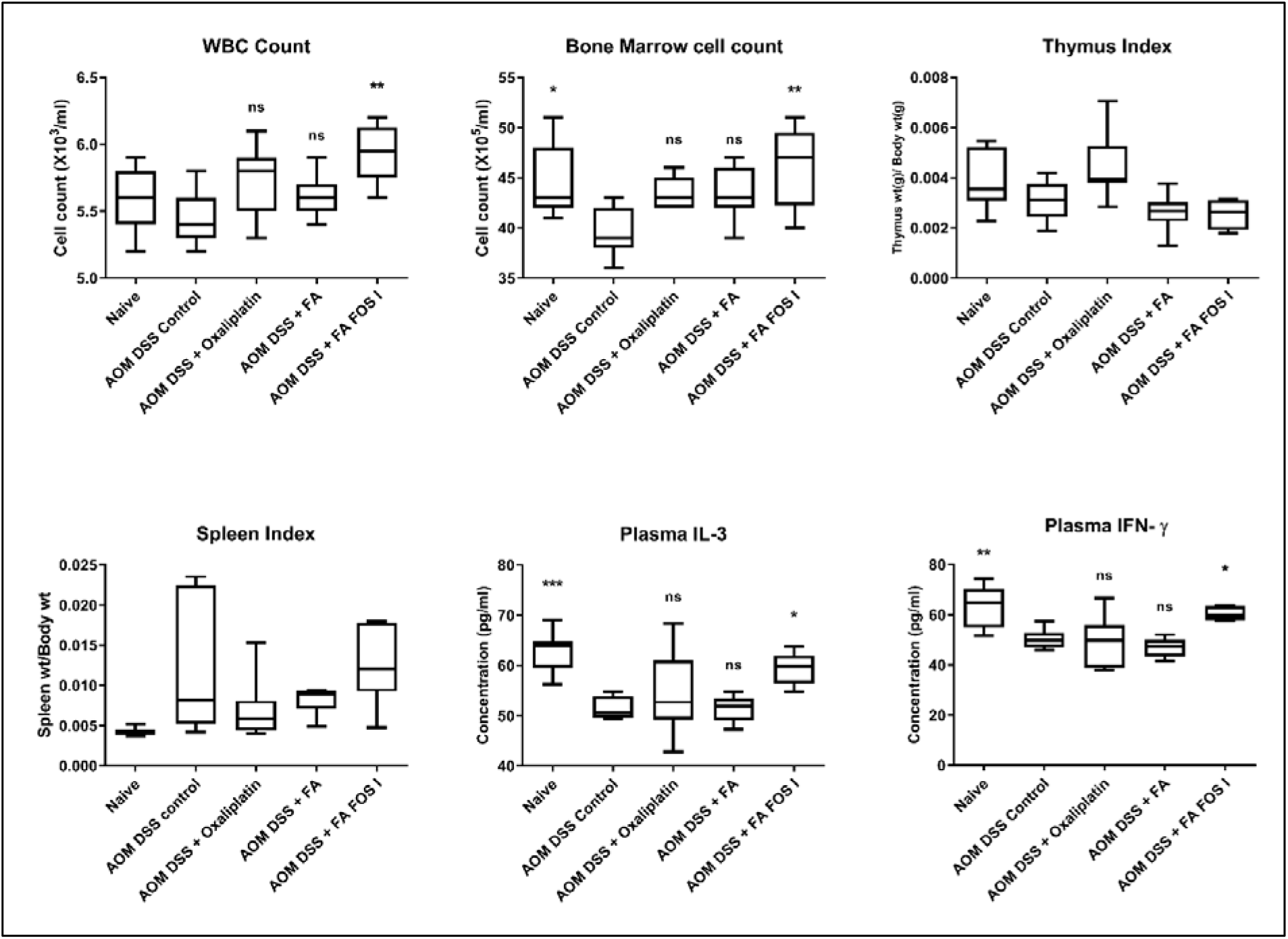
Immune status of mice group in AOM DSS induced colitis associated colon cancer mice model treated with Oxaliplatin, FA, FA FOS I and FA FOS II for four weeks. The box whisker plot represents the mean of six independent replicate animals (n=6) ± SD. * p<0.01, **p<0.001, ***p<0.0001, ns= non-significant vs AOM DSS control group.

**Fig. 25.**
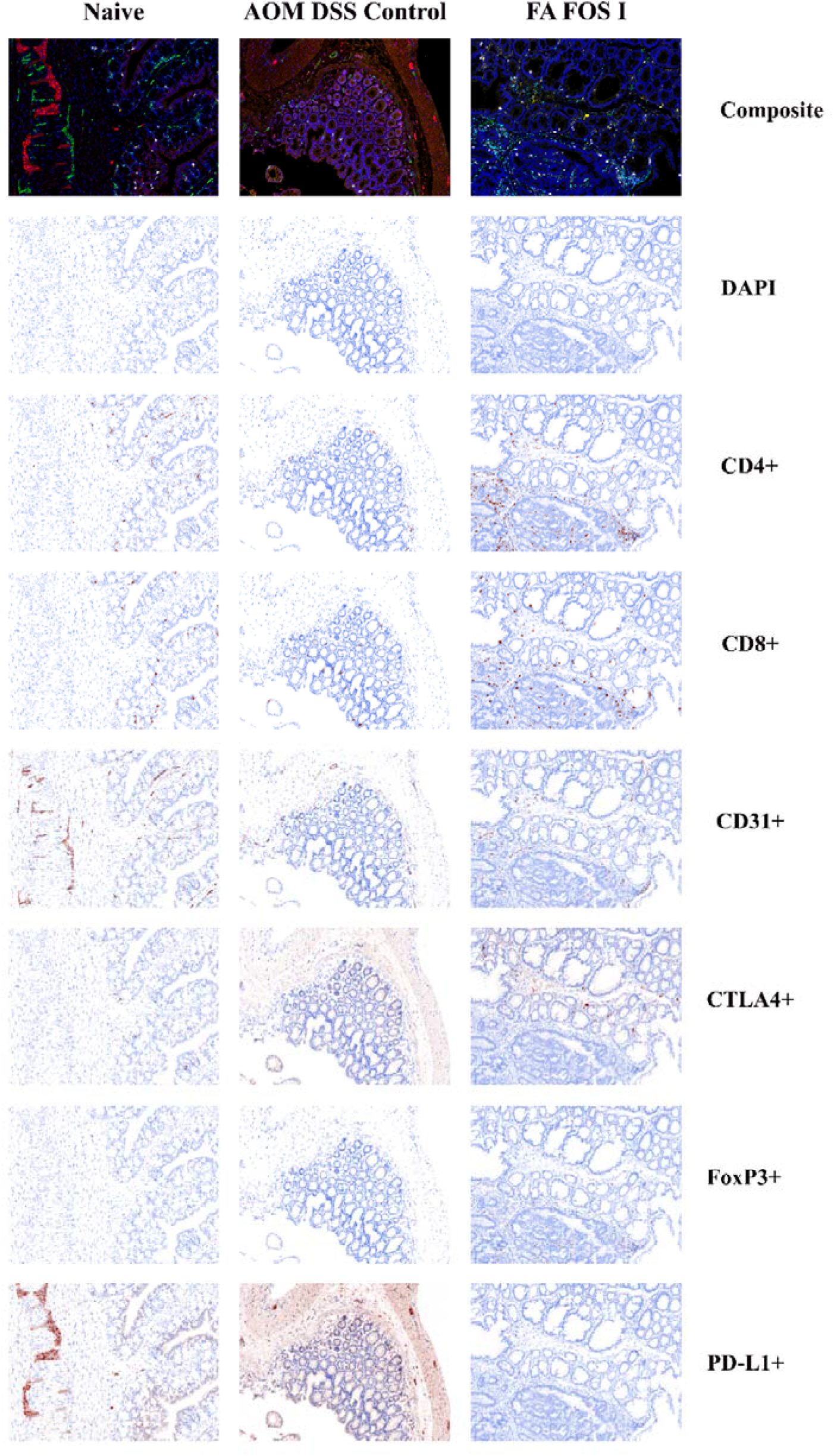
Representative multiplex immunofluorescence photomicrograph of colon tissue from the experimental group of AOM DSS induced colitis associated colon cancer model treated with indicated compound for four weeks: Top row- composite image showing all the markers in the context of host immune surveillance/status- CD4 = blue, CD8 = maroon, CD31 = green, FoxP3 = orange, PD-L1 = red, CTLA4 = yellow. Photomicrographs below the composite IF displaying the un-mixed channel of each indicated immune markers, its localization and distribution in the tumour tissue or tumour microenvironment (TME).

**Fig. 26.**
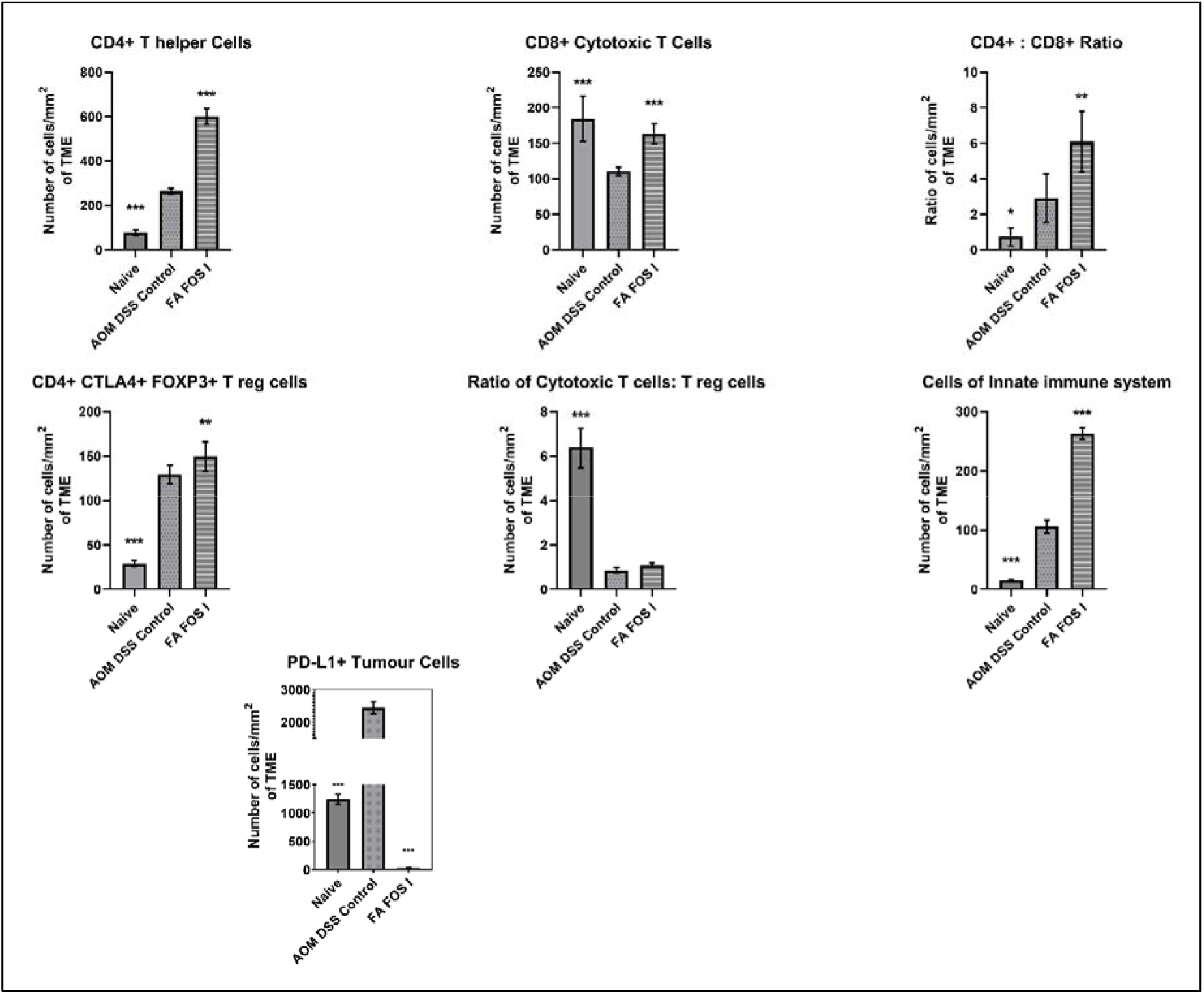
Counts of the specific immune cells found infiltrating the tumour tissue or in the TME of the colon of CAC mice in different experimental group as indicated. The data is derived from three independent experiments of annotation of multiplex IF histochemistry of three animals from each experimental group, (n=3) ± SD and which is a mean of six different field of view in each colon tissue. * p<0.01, **p<0.001, ***p<0.0001, ns= non-significant vs AOM DSS control group.

The infiltration of the immune cells into the tumour tissues or tumour microenvironment (TME) is determined by multiplex immunofluorescence imaging and inform tissue finder analysis. The TME with infiltrated immune cells such as CD4+ helper T cells and CD8+ activated or cytotoxic T cells are considered to be an good prognostic factor in several cancers (99,100). Apart from the adaptive immune system the innate immune cells are also known to play a vital role in improvement of therapeutic outcomes by infiltrating TME with cells of the innate immune wing like dendritic cells (101), macrophages (101)and natural killer cells (102). CD4+ T cells are considered to be highly polyfunctional and versatile cells of the adaptive T cell immunity along with their familial lineage of CD8+ cytotoxic T cells. The CD4+ are differentiated into various sub types of various function which are context dependent based on the signals received, basically their role is to provide help to the appropriate effector immune cells and play a role of primary co-ordination of immune response. They are also involved in the secretion of effector cytokines such as interferon- γ (INF-γ) and tumour necrosis factor- α (TNF-α). In certain conditions direct cytotoxicity against the tumour cells were also reported (103). Recent reports and clinical trials also suggest that in the absence of the CD4+ T cells, the anti-tumour response of CD8+ cytotoxic T cells were not observed (104,105). In order to determine the tumour sensitizing function of the FA FOS conjugate to the adaptive wing of the immune system, the recruitment of the CD4+ T helper cells and CD8+ cytotoxic cells in to the TME was determined by multiplex immune fluorescence analysis of the colon tissue of the AOM DSS CAC model experimental groups. It was observed that the Naïve group depicted very less CD4+ T cells in the colon tissue compared to AOM DSS CAC group which was a 3.4-fold (p<0.0001) increase compared to naïve group. The naïve group showed an average of 76.83± 12.1 cells/mm^2^ in the colon tissue while the AOM DSS CAC group TME showed 265± 13 cells/mm^2^. In the FA FOS I treatment group there was observed a 2.26-fold increase (p<0.0001) in CD4+ T cells compared to AOM DSS CAC control group with 599.5 ±33.9 cells/mm^2^. The CD8+ Cytotoxic T cells were found to be decreased in AOM DSS CAC control group when compared to naïve group, the naïve group had 184.5 ±31.8 cells/mm^2^ while the AOM DSS CAC control had only 110.66 ±5.8 cells/mm^2^ which was an 40.02% (p<0.0001) decrease compared to the naïve group. In the FA FOS I treatment group there was observed 163 ±14.2 cells/mm^2^ which was an 47.29% (P<0.0001) increase for FA FOS I compared to the AOM DSS CAC control group. The ratio of CD4+: CD8+ T cells were determined for the experimental group which showed an decreased profile for naïve group with 0.726± 0.5 compared to AOM DSS CAC control which showed 2.9± 1.3. The FA FOS I treatment group showed a ratio of 6.08± 1.69 which was a 2.09-fold (p<0.001) increase. These results also concurs with the previous reports on enhanced T cell response on treatment with certain selective chemotherapeutics (106). The T-reg cells which are usually CTLA4+ Foxp3+ are found to be associated with reducing the inflammatory response produced by the cytotoxic T lymphocytes and T helper cells, macrophages and granulocytes (106) which may be an essential role to preventing the un-controlled inflammation fuelling the cell proliferation and colorectal tumorigenesis by means of inflammation-driven cellular plasticity (8,107,108). In the present study the naïve group showed the presence of only 29± 3.4 cells/mm^2^ T-reg cells in the colon tissue but there was observed an increase of 4.45-fold CTLA4+ and Foxp3+ T-reg cells in the TME with an average of 129.33 ±10.3 cells/mm^2^ in AOM DSS CAC control group. This may be due to the inflammatory condition prevailing in the colon due to the administration of AOM and DSS causing colitis and inflammation related morbidities. In FA FOS I treatment there was observed an 15.59% (p<0.001) increase in the T-reg cells with 149.5 ±16.8 cells/mm^2^ compared to AOM DSS CAC control group. This active response of FA FOS I may be due to their physico mechanical properties and physiological properties involving interaction with the mucosal membrane of the large intestine eliciting a positive modulatory response. The ratio between the cytotoxic T cells and T-reg cells in the TME infiltrate may play an important role in abatement of tumour cells or suppression of immune mediated attack on the tumour tissue depending on the variability in stage and type of cancer and also depend on the cell type and functionality in the periphery of the TME (109). In this study there was an decrease in the ratio of cytotoxic T cells: T reg cells by 86% (p<0.0001) with naïve showing a ratio of 6.36± 0.88 vs AOM DSS CAC group with 0.85± 0.13. In FA FOS I there was observed an 27.42% increase in the ratio compared to AOM DSS CAC control with 1.09± 0.083. The CD31 marker which is also known as platelet endothelial cell adhesion molecule-1 (PECAM-1) are transmembrane homophilic receptor expressed by endothelial cells, platelets, granulocytes, dendritic cells, natural killer cells and dendritic cells. In this study the CD31+ CD4- CD8- Ki67- phenotype was used to recognize the tumour infiltrating immune cells from the innate immune system in the TME (110). In this study there was an increase in 7.06-fold (p<0.0001) of cells from the innate immune response infiltrating the TME in the AOM DSS CAC control compared to the naïve control group. The naïve control group showed only 15 ±1.38 cells/mm^2^ in the colon tissue compared to 106± 10.8 cells/mm^2^ in the AOM DSS CAC control TME. There was observed an 148.20% and 228.92% increase in the cells from the innate immune wing infiltrating the tumour in FA FOS I treatment group compared to the AOM DSS CAC control group (p<0.0001) with 263.1 ±10.1 cells/mm^2^. The programmed death ligand 1 (PD-L1) are considered to be a crucial modulator of the immune mediated killing of tumour cells by activated T cells. They are check point proteins which are produced in the surface of the tumour cells which interacts with the PD-1 receptor of the T cells and suppress their activity. Thus, this cloaking of the tumour cells with PD-L1 would mislead the activated T-cells in the TME as a harmless self-cell and evade immune mediated killing and engagement of T-cells. PD-L1 activity are also known to induce apoptosis in T-cells and the enhanced expression of the same is associated with poor patient prognosis and reduced patient survival (111). In the present study there was an increase in expression of PD-L1 96.91% (p<0.0001) in the tumour cells of AOM DSS CAC control compared to Naïve group which had only 1241.66 ±84.4 cells/mm^2^ expressing PD-L1 in the normal colon vs an average of 2445± 182.4 cells/mm^2^ in AOM DSS CAC control group. On treatment with FA FOS I there was observed an 98.95% decrease (p<0.0001) in the PD-L1 expression in the tumour cells with only 25.66 ±3.9 cells/mm^2^. There is present and multifunctional regulator named COP9 signalosome 5 (CSN5) initially identified in *Drosophila* as development related protein. Later it was discovered that NF-κb- P65-induced COP9 signalosome 5 (CSN5) is required for the TNF-α mediated PD-L1 stabilization in cancer cells (112). In this present study the FA FOS conjugates might be acting on the NF-κb-P65 axis and thereby inhibiting CSN5 leading to the downregulation of PD-L1 expression, this is also evident from the inflammatory panel analysis of the colon (Fig. 23) it can be observed that the treatment with FA FOS I conjugate to AOM DSS CAC mice drastically reduced the presence of TNF- α in the colon. Similar evidences has been reported on curcumin which is an di-ferulate family of ferulic acid which is known to inhibit CSN5 and thereby diminishing the expression of PD-L1 in tumour cells (111,112). Curcumin treatment was also reported to be modulating the immunosuppressive tumour microenvironment into immune active hot tumours by inhibiting the expression of PD-L1 in the tumour cells and also inhibiting p-STAT3Y705. Studies also supported the antitumour immune response efficacy of curcumin in oral squamous cell carcinoma (113), bladder carcinoma (114) and melanoma (115). Several flavonoids, especially apigenin are also reported to supress PD-L1 expression by their inhibitory action on STAT1 phosphorylation and thereby sensitizing the cancer cells to immune system and engagement of anti-tumour immune response (112).

## 4 Conclusion

The in-vitro evaluation of the anti-cancer properties of the FA FOS I conjugates revealed that they induced apoptosis selectively in the cancer cells by the fragmentation of DNA and activation of mitochondrial mediated apoptosis. The FA FOS I also caused cell cycle arrest at G1/S phase by mediating DNA damage-based activation of checkpoint factors and cyclins, stalling the cell cycle and preventing the cell proliferation. It was found that the FA FOS I microparticle ameliorated the AOM DSS mediated colitis associated colon cancer by reducing the inflammation, inhibition of the aberrant proliferation of cells and by inducing apoptosis in the tumour cells by the translocation of β-catenin into the nucleus, inhibition cyclin D1 and COX-2 and supressing the proliferative proteins such as Ki67, PCNA and upregulating the tumour suppressor protein p53. The FA FOS I microparticle were also found to be stimulating the immune system and enhance the immune mediated killing of the cancer cells in-vivo by sensitizing the cancer cells to immune cell engagement and killing, by down regulating the expression of PD-L1 in the tumour cells and recruiting both the innate immune cells and adaptive immune wing of the immune system to the TME. Several carbohydrate molecules are known to act as an adjuvants in stimulating the immune system (116–118). Here the immune stimulating function could be due to the synergistic action of neutral fructo-oligosaccharide and negatively charged ferulic acid (phenolic acid) conjugated to it (119). The physiological antioxidant property of FA FOS I was evaluated, and their antioxidant activity in-vivo was also evaluated in AOM DSS CAC model. There was observed an enhanced antioxidant profile on treatment with FA FOS conjugate in-vivo. The inflammation ameliorating property of the FA FOS combined with its anti-tumour and immune modulatory function prospects for the evaluation of the pharmacokinetics, pharmacodynamics, metabolomics, and host microbiota modulation activity of the FA FOS conjugates. The investigation of the gene expression profile and delineation of the molecular pathway associated with the administration of FA FOS conjugate in the in-vivo system would shed light on the mechanistic action of the FA FOS conjugates in ameliorating colorectal carcinogenesis.

## Author contribution

**Eldin M Johnson and Rasu Jayabalan** conceptualized the research idea.

**Eldin M. Johnson:** Conceptualization, Design of study, Methodology, Investigation, Formal analysis, Data curation, Software, Writing - original draft and editing.

**Joo-Won Suh:** Resources, Project administration, Funding acquisition, Supervision, Editing.

**Samir Kumar Patra:** Supervision, Manuscript review and editing.

## Acknowledgments Funding

This work is carried out with the support of the “Cooperative Research Program for Agriculture Science & Technology Development (Project No. PJ01319101 assigned to Joo-Won Suh)”, the Rural Development Administration of the Republic of Korea.

## References

1. Lopez-Lazaro M. A Simple and Reliable Approach for Assessing Anticancer Activity In Vitro. Curr Med Chem. 2015 Mar 30;22(11):1324–34.

2. López-Lázaro M. Two preclinical tests to evaluate anticancer activity and to help validate drug candidates for clinical trials. Oncoscience. 2015;2(2):91.

3. Johnson EM, Lee H, Jayabalan R, Suh J-W. Ferulic acid grafted self-assembled fructo- oligosaccharide micro particle for targeted delivery to colon. Carbohydr Polym. 2020;247(June):116550.

4. Barbier A. Rôle de la pharmacologie in vivo en recherche et développement. In: Therapie. Therapie; 2004. p. 45–50.

5. Deb M, Sengupta D, Kar S, Rath SK, Roy S, Das G, et al. Epigenetic drift towards histone modifications regulates CAV1 gene expression in colon cancer. Gene. 2016 Apr 25;581(1):75–84.

6. Kar S, Sengupta D, Deb M, Shilpi A, Parbin S, Rath SK, et al. Expression profiling of DNA methylation-mediated epigenetic gene-silencing factors in breast cancer. Clin Epigenetics. 2014 Oct 13;6(1):1–13.

7. Deb M, Sengupta D, Rath SK, Kar S, Parbin S, Shilpi A, et al. Clusterin gene is predominantly regulated by histone modifications in human colon cancer and ectopic expression of the nuclear isoform induces cell death. Biochim Biophys Acta - Mol Basis Dis. 2015 Aug 1;1852(8):1630–45.

8. Schmitt M, Greten FR. The inflammatory pathogenesis of colorectal cancer. Vol. 21, Nature Reviews Immunology. 2021. p. 653–67.

9. Roy A, Patra SK. Lipid Raft Facilitated Receptor Organization and Signaling: A Functional Rheostat in Embryonic Development, Stem Cell Biology and Cancer. Vol. 19, Stem Cell Reviews and Reports. Springer; 2022. p. 2–25.

10. Bhattacharyya A, Chattopadhyay R, Mitra S, Crowe SE. Oxidative stress: An essential factor in the pathogenesis of gastrointestinal mucosal diseases. Physiol Rev. 2014;94(2):329–54.

11. Greten FR, Grivennikov SI. Inflammation and Cancer: Triggers, Mechanisms, and Consequences. Vol. 51, Immunity. Immunity; 2019. p. 27–41.

12. Bożyk A, Wojas-Krawczyk K, Krawczyk P, Milanowski J. Tumor Microenvironment—A Short Review of Cellular and Interaction Diversity. Vol. 11, Biology. Multidisciplinary Digital Publishing Institute (MDPI); 2022.

13. Neufert C, Becker C, Neurath MF. An inducible mouse model of colon carcinogenesis for the analysis of sporadic and inflammation-driven tumor progression. Nat Protoc. 2007 Aug;2(8):1998–2004.

14. De Robertis M, Massi E, Poeta M, Carotti S, Morini S, Cecchetelli L, et al. The AOM/DSS murine model for the study of colon carcinogenesis: From pathways to diagnosis and therapy studies. J Carcinog. 2011;10.

15. Parang B, Barrett CW, Williams CS. AOM/DSS Model of Colitis-Associated Cancer. Methods Mol Biol. 2016;1422:297–307.

16. Liu X, Quan N. Immune Cell Isolation from Mouse Femur Bone Marrow. BIO-PROTOCOL. 2015 Oct 10;5(20).

17. Le Naour J, Montégut L, Joseph A, Garbin K, Vacchelli E, Kroemer G, et al. Improved Swiss- rolling method for histological analyses of colon tissue. MethodsX. 2022 Jan 1;9:101630.

18. Jansen KA, Donato DM, Balcioglu HE, Schmidt T, Danen EHJ, Koenderink GH. A guide to mechanobiology: Where biology and physics meet. Vol. 1853, Biochimica et Biophysica Acta - Molecular Cell Research. Elsevier; 2015. p. 3043–52.

19. Vogler M. BCL2A1: The underdog in the BCL2 family. Cell Death Differ. 2012;19(1):67–74.

20. Fabregat A, Sidiropoulos K, Viteri G, Forner O, Marin-Garcia P, Arnau V, et al. Reactome pathway analysis: A high-performance in-memory approach. BMC Bioinformatics. 2017 Mar 2;18(1):1–9.

21. So T, Lee SW, Croft M. Tumor necrosis factor/tumor necrosis factor receptor family members that positively regulate immunity. Vol. 83, International Journal of Hematology. Int J Hematol; 2006. p. 1–11.

22. Akdis M, Aab A, Altunbulakli C, Azkur K, Costa RA, Crameri R, et al. Interleukins (from IL- 1 to IL-38), interferons, transforming growth factor β, and TNF-α: Receptors, functions, and roles in diseases. Vol. 138, Journal of Allergy and Clinical Immunology. 2016. p. 984–1010.

23. Happo L, Strasser A, Cory S. BH3-only proteins in apoptosis at a glance. J Cell Sci. 2012 Mar 3;125(5):1081.

24. Cory S, Adams JM. The BCL2 family: Regulators of the cellular life-or-death switch. Vol. 2, Nature Reviews Cancer. Nat Rev Cancer; 2002. p. 647–56.

25. Saelens X, Festjens N, Vande Walle L, Van Gurp M, Van Loo G, Vandenabeele P. Toxic proteins released from mitochondria in cell death. Vol. 23, Oncogene. Oncogene; 2004. p. 2861–74.

26. So YS, Chen YB, Ivamovska I, Ranger AM, Hong SJ, Dawson VL, et al. BAD is a pro- survival factor prior to activation of its pro-apoptotic function. J Biol Chem. 2004 Oct 1;279(40):42240–9.

27. Li J, Hu L, Tian C, Lu F, Wu J, Liu L. microRNA-150 promotes cervical cancer cell growth and survival by targeting FOXO4. BMC Mol Biol. 2015 Dec 29;16(1).

28. Arimoto-Ishida E, Ohmichi M, Mabuchi S, Takahashi T, Ohshima C, Hayakawa J, et al. Inhibition of phosphorylation of a Forkhead transcription factor sensitizes human ovarian cancer cells to cisplatin. Endocrinology. 2004 Apr;145(4):2014–22.

29. 29. Kruiswijk F, Labuschagne CF, Vousden KH. P53 in survival, death and metabolic health: A lifeguard with a licence to kill. Vol. 16, Nature Reviews Molecular Cell Biology. Nature Publishing Group; 2015. p. 393–405.

30. Yu H, Lin L, Zhang Z, Zhang H, Hu H. Targeting NF-κB pathway for the therapy of diseases: mechanism and clinical study. Vol. 5, Signal Transduction and Targeted Therapy. Nature Publishing Group; 2020. p. 1–23.

31. 31. Green DR. The Mitochondrial Pathway of Apoptosis: Part I: MOMP and Beyond. Vol. 14, Cold Spring Harbor perspectives in biology. Cold Spring Harbor Laboratory Press; 2022. p. a041038.

32. Burke SP, Smith JB. Monomerization of cytosolic mature Smac attenuates interaction with IAPS and potentiation of caspase activation. PLoS One. 2010;5(10):e13094.

33. Salvesen GS, Duckett CS. IAP proteins: Blocking the road to death’s door. Vol. 3, Nature Reviews Molecular Cell Biology. Nat Rev Mol Cell Biol; 2002. p. 401–10.

34. Xiong S, Mu T, Wang G, Jiang X. Mitochondria-mediated apoptosis in mammals. Protein Cell. 2014 Oct 31;5(10):737–49.

35. Herbst RS, Frankel SR. Oblimersen sodium (genasense bcl-2 antisense oligonucleotide): A rational therapeutic to enhance apoptosis in therapy of lung cancer. In: Clinical Cancer Research. Clin Cancer Res; 2004.

36. Tse C, Shoemaker AR, Adickes J, Anderson MG, Chen J, Jin S, et al. ABT-263: A potent and orally bioavailable Bcl-2 family inhibitor. Cancer Res. 2008 May 1;68(9):3421–8.

37. Cao X, Yap JL, Newell-Rogers MK, Peddaboina C, Jiang W, Papaconstantinou HT, et al. The novel BH3 α-helix mimetic JY-1-106 induces apoptosis in a subset of cancer cells (lung cancer, colon cancer and mesothelioma) by disrupting Bcl-xL and Mcl-1 protein-protein interactions with Bak. Mol Cancer. 2013;12(1):42.

38. Fulda S, Vucic D. Targeting IAP proteins for therapeutic intervention in cancer. Vol. 11, Nature Reviews Drug Discovery. Nat Rev Drug Discov; 2012. p. 109–24.

39. Mahadevan D, Chalasani P, Rensvold D, Kurtin S, Pretzinger C, Jolivet J, et al. Phase i trial of AEG35156 an antisense oligonucleotide to xiap plus gemcitabine in patients with metastatic pancreatic ductal adenocarcinoma. Am J Clin Oncol Cancer Clin Trials. 2013 Jun;36(3):239– 43.

40. Kirkland JL, Tchkonia T. Senolytic drugs: from discovery to translation. Vol. 288, Journal of Internal Medicine. Wiley-Blackwell; 2020. p. 518–36.

41. Fan J, Wray J, Meng X, Shen Z. BCCIP is required for the nuclear localization of the p21 protein. Cell Cycle. 2009 Sep 9;8(18):3023–8.

42. Helt CE, Cliby WA, Keng PC, Bambara RA, O’Reilly MA. Ataxia telangiectasia mutated (ATM) and ATM and Rad3-related protein exhibit selective target specificities in response to different forms of DNA damage. J Biol Chem. 2005 Jan 14;280(2):1186–92.

43. Tigan AS, Bellutti F, Kollmann K, Tebb G, Sexl V. CDK6-a review of the past and a glimpse into the future: From cell-cycle control to transcriptional regulation. Vol. 35, Oncogene. Nature Publishing Group; 2016. p. 3083–91.

44. Xia Y, Liu Y, Yang C, Simeone DM, Sun TT, DeGraff DJ, et al. Dominant role of CDKN2B/p15INK4B of 9p21.3 tumor suppressor hub in inhibition of cell-cycle and glycolysis. Nat Commun. 2021 Apr 6;12(1):1–15.

45. Reiter V, Matschkal DMS, Wagner M, Globisch D, Kneuttinger AC, Müller M, et al. The CDK5 repressor CDK5RAP1 is a methylthiotransferase acting on nuclear and mitochondrial RNA. Nucleic Acids Res. 2012 Jul;40(13):6235–40.

46. Sheu YJ, Kinney JB, Lengronne A, Pasero P, Stillman B. Domain within the helicase subunit Mcm4 integrates multiple kinase signals to control DNA replication initiation and fork progression. Proc Natl Acad Sci U S A. 2014 May 6;111(18):E1899–908.

47. Goebl MG, Goetsch L, Byers B. The Ubc3 (Cdc34) ubiquitin-conjugating enzyme is ubiquitinated and phosphorylated in vivo. Mol Cell Biol. 1994 May;14(5):3022–9.

48. Wu Z, Zheng S, Yu Q. The E2F family and the role of E2F1 in apoptosis. Vol. 41, International Journal of Biochemistry and Cell Biology. Int J Biochem Cell Biol; 2009. p. 2389–97.

49. Dick FA. Structure-function analysis of the retinoblastoma tumor suppressor protein - Is the whole a sum of its parts? Vol. 2, Cell Division. Cell Div; 2007.

50. Blazek D, Kohoutek J, Bartholomeeusen K, Johansen E, Hulinkova P, Luo Z, et al. The cyclin K/Cdk12 complex maintains genomic stability via regulation of expression of DNA damage response genes. Genes Dev. 2011 Oct 15;25(20):2158–72.

51. Zou L, Elledge SJ. Sensing DNA damage through ATRIP recognition of RPA-ssDNA complexes. Science (80-). 2003 Jun 6;300(5625):1542–8.

52. Stark GR, Taylor WR. Analyzing the G2/M checkpoint. Vol. 280, Methods in molecular biology (Clifton, N.J.). Humana Press; 2004. p. 51–82.

53. Eberhart CE, Coffey RJ, Radhika A, Giardiello FM, Ferrenbach S, Dubois RN. Up-regulation of cyclooxygenase 2 gene expression in human colorectal adenomas and adenocarcinomas. Gastroenterology. 1994 Oct 1;107(4):1183–8.

54. Brabletz T, Jung A, Kirchner T. β-Catenin and the morphogenesis of colorectal cancer. Vol. 441, Virchows Archiv. Springer; 2002. p. 1–11.

55. Nelson WJ, Nusse R. Convergence of Wnt, β-Catenin, and Cadherin pathways. Vol. 303, Science. 2004. p. 1483–7.

56. Herr P, Hausmann G, Basler K. WNT secretion and signalling in human disease. Vol. 18, Trends in Molecular Medicine. 2012. p. 483–93.

57. Olsen AK, Boyd M, Danielsen ET, Troelsen JT. Current and emerging approaches to define intestinal epithelium-specific transcriptional networks. Vol. 302, American Journal of Physiology - Gastrointestinal and Liver Physiology. 2012. p. G277–86.

58. Gao N, White P, Kaestner KH. Establishment of Intestinal Identity and Epithelial- Mesenchymal Signaling by Cdx2. Dev Cell. 2009 Apr 1;16(4):588–99.

59. Crissey MAS, Guo R, Funakoshi S, Kong J, Liu J, Lynch JP. Cdx2 levels modulate intestinal epithelium maturity and Paneth cell development. Gastroenterology. 2011 Nov 13;140(2):517–528.e8.

60. Verzi MP, Shin H, He HH, Sulahian R, Meyer CA, Montgomery RK, et al. Differentiation-Specific Histone Modifications Reveal Dynamic Chromatin Interactions and Partners for the Intestinal Transcription Factor CDX2. Dev Cell. 2010 Nov 1;19(5):713–26.

61. Gross I, Duluc I, Benameur T, Calon A, Martin E, Brabletz T, et al. The intestine-specific homeobox gene Cdx2 decreases mobility and antagonizes dissemination of colon cancer cells. Oncogene. 2008 Jun 25;27(1):107–15.

62. Brabletz T, Spaderna S, Kolb J, Hlubek F, Faller G, Bruns CJ, et al. Down-regulation of the homeodomain factor Cdx2 in colorectal cancer by collagen type I: An active role for the tumor environment in malignant tumor progression. Cancer Res. 2004 Oct 1;64(19):6973–7.

63. Kim S, Domon-Dell C, Wang Q, Chung DH, Di Cristofano A, Pandolfi PP, et al. PTEN and TNF-α regulation of the intestinal-specific Cdx-2 homeobox gene through a PI3K, PKB/Akt, and NF-κB-dependent pathway. Gastroenterology. 2002 Oct 1;123(4):1163–78.

64. Gregorieff A, Pinto D, Begthel H, Destrée O, Kielman M, Clevers H. Expression pattern of Wnt signaling components in the adult intestine. Gastroenterology. 2005 Aug 1;129(2):626–38.

65. Clevers H. Wnt/β-Catenin Signaling in Development and Disease. Vol. 127, Cell. 2006. p. 469–80.

66. Olsen AK, Coskun M, Bzorek M, Kristensen MH, Danielsen ET, Jørgensen S, et al. Regulation of APC and AXIN2 expression by intestinal tumor suppressor CDX2 in colon cancer cells. Carcinogenesis. 2013 Feb 7;34(6):1361–9.

67. Guo RJ, Funakoshi S, Lee HH, Kong J, Lynch JP. The intestine-specific transcription factor Cdx2 inhibits β-catenin/TCF transcriptional activity by disrupting the β-catenin-TCF protein complex. Carcinogenesis. 2010 Sep 4;31(2):159–66.

68. Hinkel I, Duluc I, Martin E, Guenot D, Freund J, Gross I. Cdx2 controls expression of the protocadherin Mucdhl, an inhibitor of growth and β-catenin activity in colon cancer cells. Gastroenterology. 2012 Dec 24;142(4):875–885.e3.

69. Thiagarajan DG, Bennink MR, Bourquin LD, Kavas FA. Prevention of precancerous colonic lesions in rats by soy flakes, soy flour, genistein, and calcium. In: American Journal of Clinical Nutrition. American Society for Nutrition; 1998. p. 1394S–1399S.

70. Coskun M, Olsen AK, Bzorek M, Holck S, Engel UH, Nielsen OH, et al. Involvement of CDX2 in the cross talk between TNF-α and Wnt signaling pathway in the colon cancer cell line Caco-2. Carcinogenesis. 2014 Feb 5;35(5):1185–92.

71. Vermeulen L, De Sousa E Melo F, Van Der Heijden M, Cameron K, De Jong JH, Borovski T, et al. Wnt activity defines colon cancer stem cells and is regulated by the microenvironment. Nat Cell Biol. 2010 Apr 25;12(5):468–76.

72. Du Q, Wang Y, Liu C, Wang H, Fan H, Li Y, et al. Chemopreventive activity of GEN-27, a genistein derivative, in colitis-associated cancer is mediated by p65-CDX2-β-catenin axis. Oncotarget. 2016 Apr 1;7(14):17870–84.

73. Deschner EE, Lipkin M. Proliferative patterns in colonic mucosa in familial polyposis. Cancer. 1975;35(2):413–8.

74. Biasco G, Minarini A, Miglioli M, Barbara L, Lipkin M, Higgins P. Proliferative and Antigenic Properties off Rectal Cells in Patients with Chronic Ulcerative Colitis. Cancer Res. 1984 Nov 1;44(11):5450–4.

75. Kubben FJGM, Peeters-Haesevoets A, Engels LGJB, Baeten CGMI, Schutte B, Arends JW, et al. Proliferating cell nuclear antigen (PCNA): A new marker to study human colonic cell proliferation. Gut. 1994 Apr 1;35(4):530–5.

76. Suzuki R, Fukui T, Kishimoto M, Miyamoto S, Takahashi Y, Takeo M, et al. Smad2/3 linker phosphorylation is a possible marker of cancer stem cells and correlates with carcinogenesis in a mouse model of colitis-associated colorectal cancer. J Crohn’s Colitis. 2015 Jul 1;9(7):565– 74.

77. 77. Adams JM, Cory S. The Bcl-2 apoptotic switch in cancer development and therapy. Vol. 26, Oncogene. Nature Publishing Group; 2007. p. 1324–37.

78. 78. Lowe SW, Cepero E, Evan G. Intrinsic tumour suppression. Vol. 432, Nature. Nature Publishing Group; 2004. p. 307–15.

79. 79. Schmitt M, Greten FR. The inflammatory pathogenesis of colorectal cancer. Vol. 21, Nature Reviews Immunology. Nature Publishing Group; 2021. p. 653–67.

80. Canli Ö, Nicolas AM, Gupta J, Finkelmeier F, Goncharova O, Pesic M, et al. Myeloid Cell- Derived Reactive Oxygen Species Induce Epithelial Mutagenesis. Cancer Cell. 2017 Dec 11;32(6):869–883.e5.

81. Janakiram NB, Rao C V. Chemoprevention of colon cancer by iNOS-selective inhibitors. In: Forum on Immunopathological Diseases and Therapeutics. NIH Public Access; 2012. p. 155–67.

82. Shaked H, Hofseth LJ, Chumanevich A, Chumanevich AA, Wang J, Wang Y, et al. Chronic epithelial NF-κB activation accelerates APC loss and intestinal tumor initiation through iNOS up-regulation. Proc Natl Acad Sci U S A. 2012 Aug 28;109(35):14007–12.

83. Karin M. Nuclear factor-κB in cancer development and progression. Vol. 441, Nature. Nature; 2006. p. 431–6.

84. Greten FR, Eckmann L, Greten TF, Park JM, Li ZW, Egan LJ, et al. IKKβ links inflammation and tumorigenesis in a mouse model of colitis-associated cancer. Cell. 2004 Aug 6;118(3):285–96.

85. Popivanova BK, Kitamura K, Wu Y, Kondo T, Kagaya T, Kaneko S, et al. Blocking TNF-α in mice reduces colorectal carcinogenesis associated with chronic colitis. J Clin Invest. 2008 Feb 1;118(2):560–70.

86. Pollard JW. Tumour-educated macrophages promote tumour progression and metastasis. Vol. 4, Nature Reviews Cancer. European Association for Cardio-Thoracic Surgery; 2004. p. 71–8.

87. Breese EJ, Michie CA, Nicholls SW, Murch SH, Williams CB, Domizio P, et al. Tumor necrosis factor α-producing cells in the intestinal mucosa of children with inflammatory bowel disease. Gastroenterology. 1994 Jun 1;106(6):1455–66.

88. Waldner MJ, Foersch S, Neurath MF. Interleukin-6 - A key regulator of colorectal cancer development. Vol. 8, International Journal of Biological Sciences. Ivyspring International Publisher; 2012. p. 1248–53.

89. Rébé C, Ghiringhelli F. Interleukin-1β and cancer. Vol. 12, Cancers. Multidisciplinary Digital Publishing Institute; 2020. p. 1–31.

90. Becker C, Fantini MC, Schramm C, Lehr HA, Wirtz S, Nikolaev A, et al. TGF-β suppresses tumor progression in colon cancer by inhibition of IL-6 trans-signaling. Immunity. 2004 Oct;21(4):491–501.

91. Grivennikov S, Karin E, Terzic J, Mucida D, Yu GY, Vallabhapurapu S, et al. IL-6 and Stat3 Are Required for Survival of Intestinal Epithelial Cells and Development of Colitis-Associated Cancer. Cancer Cell. 2009 Feb 3;15(2):103–13.

92. Waldner MJ, Wirtz S, Jefremow A, Warntjen M, Neufert C, Atreya R, et al. VEGF receptor signaling links inflammation and tumorigenesis in colitis-associated cancer. J Exp Med. 2010 Dec 20;207(13):2855–68.

93. Hai Ping P, Feng Bo T, Li L, Nan Hui Y, Hong Z. IL-1β/NF-kb signaling promotes colorectal cancer cell growth through miR-181a/PTEN axis. Arch Biochem Biophys. 2016 Aug 15;604:20–6.

94. Kaler P, Augenlicht L, Klampfer L. Macrophage-derived IL-1Β stimulates Wnt signaling and growth of colon cancer cells: A crosstalk interrupted by vitamin D"3. Oncogene. 2009 Aug 24;28(44):3892–902.

95. Dougan M, Dranoff G, Dougan SK. GM-CSF, IL-3, and IL-5 Family of Cytokines: Regulators of Inflammation. Vol. 50, Immunity. Elsevier; 2019. p. 796–811.

96. Hirst WJR, Buggins A, Darling D, Gäken J, Farzaneh F, Mufti GJ. Enhanced immune costimulatory activity of primary acute myeloid leukaemia blasts after retrovirus-mediated gene transfer of B7.1. Gene Ther. 1997;4(7):691–9.

97. Guthridge MA, Stomski FC, Thomas D, Woodcock JM, Bagley CJ, Berndt MC, et al. Mechanism of activation of the GM-CSF, IL-3, and IL-5 family of receptors. Vol. 16, Stem Cells. Oxford Academic; 1998. p. 301–13.

98. Castro F, Cardoso AP, Gonçalves RM, Serre K, Oliveira MJ. Interferon-gamma at the crossroads of tumor immune surveillance or evasion. Vol. 9, Frontiers in Immunology. Frontiers Media S.A.; 2018. p. 847.

99. Binnewies M, Roberts EW, Kersten K, Chan V, Fearon DF, Merad M, et al. Understanding the tumor immune microenvironment (TIME) for effective therapy. Nat Med. 2018 May 1;24(5):541–50.

100. Hegde PS, Karanikas V, Evers S. The where, the when, and the how of immune monitoring for cancer immunotherapies in the era of checkpoint inhibition. Clin Cancer Res. 2016 Apr 15;22(8):1865–74.

101. Barkal AA, Brewer RE, Markovic M, Kowarsky M, Barkal SA, Zaro BW, et al. CD24 signalling through macrophage Siglec-10 is a target for cancer immunotherapy. Nature. 2019 Aug 15;572(7769):392–6.

102. Zemek RM, De Jong E, Chin WL, Schuster IS, Fear VS, Casey TH, et al. Sensitization to immune checkpoint blockade through activation of a STAT1/NK axis in the tumor microenvironment. Sci Transl Med. 2019 Jul 17;11(501).

103. Tay RE, Richardson EK, Toh HC. Revisiting the role of CD4+ T cells in cancer immunotherapy—new insights into old paradigms. Vol. 28, Cancer Gene Therapy. Nature Publishing Group; 2021. p. 5–17.

104. Antony PA, Piccirillo CA, Akpinarli A, Finkelstein SE, Speiss PJ, Surman DR, et al. CD8 + T Cell Immunity Against a Tumor/Self-Antigen Is Augmented by CD4 + T Helper Cells and Hindered by Naturally Occurring T Regulatory Cells. J Immunol. 2005 Mar 3;174(5):2591– 601.

105. Aarntzen EHJG, De Vries IJM, Lesterhuis WJ, Schuurhuis D, Jacobs JFM, Bol K, et al. Targeting CD4+ T-helper cells improves the induction of antitumor responses in dendritic cell-based vaccination. Cancer Res. 2013 Jan 1;73(1):19–29.

106. Hodge JW, Garnett CT, Farsaci B, Palena C, Tsang KY, Ferrone S, et al. Chemotherapy- induced immunogenic modulation of tumor cells enhances killing by cytotoxic T lymphocytes and is distinct from immunogenic cell death. Int J Cancer. 2013 Aug 8;133(3):624–36.

107. Stromnes IM, Hulbert A, Pierce RH, Greenberg PD, Hingorani SR. T-cell localization, activation, and clonal expansion in human pancreatic ductal adenocarcinoma. Cancer Immunol Res. 2017 Nov 1;5(11):978–91.

108. Asaka S, Yen TT, Wang TL, Shih IM, Gaillard S. T cell-inflamed phenotype and increased Foxp3 expression in infiltrating T-cells of mismatch-repair deficient endometrial cancers. Mod Pathol. 2019 Apr 1;32(4):576–84.

109. Malka D, Lièvre A, André T, Taïeb J, Ducreux M, Bibeau F. Immune scores in colorectal cancer: Where are we? Vol. 140, European Journal of Cancer. Eur J Cancer; 2020. p. 105–18.

110. Marelli-Berg FM, Clement M, Mauro C, Caligiuri G. An immunologist’s guide to CD31 function in T-cells. J Cell Sci. 2013 Jun 1;126(11):2343–52.

111. Yang YCSH, Li ZL, Shih YJ, Bennett JA, Whang-Peng J, Lin HY, et al. Herbal medicines attenuate PD-L1 expression to induce anti-proliferation in obesity-related cancers. Vol. 11, Nutrients. Multidisciplinary Digital Publishing Institute (MDPI); 2019.

112. Lim SO, Li CW, Xia W, Cha JH, Chan LC, Wu Y, et al. Deubiquitination and Stabilization of PD-L1 by CSN5. Cancer Cell. 2016 Dec 12;30(6):925–39.

113. Liao F, Liu L, Luo E, Hu J. Curcumin enhances anti-tumor immune response in tongue squamous cell carcinoma. Arch Oral Biol. 2018 Aug 1;92:32–7.

114. Sshao Y, Zhu W, Da J, Xu M, Wang Y, Zhou J, et al. Bisdemethoxycurcumin in combination with α-PD-L1 antibody boosts immune response against bladder cancer. Onco Targets Ther. 2017 May 22;10:2675–83.

115. Xu L, Zhang Y, Tian K, Chen X, Zhang R, Mu X, et al. Apigenin suppresses PD-L1 expression in melanoma and host dendritic cells to elicit synergistic therapeutic effects. J Exp Clin Cancer Res. 2018 Oct 29;37(1).

116. Pifferi C, Fuentes R, Fernández-Tejada A. Natural and synthetic carbohydrate-based vaccine adjuvants and their mechanisms of action. Vol. 5, Nature Reviews Chemistry. Nature Publishing Group; 2021. p. 197–216.

117. Barry KC, Hsu J, Broz ML, Cueto FJ, Binnewies M, Combes AJ, et al. A natural killer– dendritic cell axis defines checkpoint therapy–responsive tumor microenvironments. Nat Med. 2018 Aug 1;24(8):1178–91.

118. Lin K, Kasko AM. Carbohydrate-based polymers for immune modulation. ACS Macro Lett. 2014 Jul 15;3(7):652–7.

119. Nishat S, Andreana PR. Entirely carbohydrate-based vaccines: An emerging field for specific and selective immune responses. Vol. 4, Vaccines. Multidisciplinary Digital Publishing Institute; 2016. p. 19.

